# Towards complete and error-free genome assemblies of all vertebrate species

**DOI:** 10.1101/2020.05.22.110833

**Authors:** Arang Rhie, Shane A. McCarthy, Olivier Fedrigo, Joana Damas, Giulio Formenti, Sergey Koren, Marcela Uliano-Silva, William Chow, Arkarachai Fungtammasan, Gregory L. Gedman, Lindsey J. Cantin, Francoise Thibaud-Nissen, Leanne Haggerty, Chul Lee, Byung June Ko, Juwan Kim, Iliana Bista, Michelle Smith, Bettina Haase, Jacquelyn Mountcastle, Sylke Winkler, Sadye Paez, Jason Howard, Sonja C. Vernes, Tanya M. Lama, Frank Grutzner, Wesley C. Warren, Christopher Balakrishnan, Dave Burt, Julia M. George, Mathew Biegler, David Iorns, Andrew Digby, Daryl Eason, Taylor Edwards, Mark Wilkinson, George Turner, Axel Meyer, Andreas F. Kautt, Paolo Franchini, H William Detrich, Hannes Svardal, Maximilian Wagner, Gavin J.P. Naylor, Martin Pippel, Milan Malinsky, Mark Mooney, Maria Simbirsky, Brett T. Hannigan, Trevor Pesout, Marlys Houck, Ann Misuraca, Sarah B. Kingan, Richard Hall, Zev Kronenberg, Jonas Korlach, Ivan Sović, Christopher Dunn, Zemin Ning, Alex Hastie, Joyce Lee, Siddarth Selvaraj, Richard E. Green, Nicholas H. Putnam, Jay Ghurye, Erik Garrison, Ying Sims, Joanna Collins, Sarah Pelan, James Torrance, Alan Tracey, Jonathan Wood, Dengfeng Guan, Sarah E. London, David F. Clayton, Claudio V. Mello, Samantha R. Friedrich, Peter V. Lovell, Ekaterina Osipova, Farooq O. Al-Ajli, Simona Secomandi, Heebal Kim, Constantina Theofanopoulou, Yang Zhou, Robert S. Harris, Kateryna D. Makova, Paul Medvedev, Jinna Hoffman, Patrick Masterson, Karen Clark, Fergal Martin, Kevin Howe, Paul Flicek, Brian P. Walenz, Woori Kwak, Hiram Clawson, Mark Diekhans, Luis Nassar, Benedict Paten, Robert H.S. Kraus, Harris Lewin, Andrew J. Crawford, M. Thomas P. Gilbert, Guojie Zhang, Byrappa Venkatesh, Robert W. Murphy, Klaus-Peter Koepfli, Beth Shapiro, Warren E. Johnson, Federica Di Palma, Tomas Margues-Bonet, Emma C. Teeling, Tandy Warnow, Jennifer Marshall Graves, Oliver A. Ryder, David Hausler, Stephen J. O’Brien, Kerstin Howe, Eugene W. Myers, Richard Durbin, Adam M. Phillippy, Erich D. Jarvis

## Abstract

High-quality and complete reference genome assemblies are fundamental for the application of genomics to biology, disease, and biodiversity conservation. However, such assemblies are only available for a few non-microbial species^1–4^. To address this issue, the international Genome 10K (G10K) consortium^5,6^ has worked over a five-year period to evaluate and develop cost-effective methods for assembling the most accurate and complete reference genomes to date. Here we summarize these developments, introduce a set of quality standards, and present lessons learned from sequencing and assembling 16 species representing major vertebrate lineages (mammals, birds, reptiles, amphibians, teleost fishes and cartilaginous fishes). We confirm that long-read sequencing technologies are essential for maximizing genome quality and that unresolved complex repeats and haplotype heterozygosity are major sources of error in assemblies. Our new assemblies identify and correct substantial errors in some of the best historical reference genomes. Adopting these lessons, we have embarked on the Vertebrate Genomes Project (VGP), an effort to generate high-quality, complete reference genomes for all ~70,000 extant vertebrate species and help enable a new era of discovery across the life sciences.

## Main

Reference genomes have served as the foundation of genomics since the generation of the first model organism sequences in the 1990s, culminating with the first human reference genome in 2003^3^. A chromosome-level reference genome underpins the study of functional, comparative, and population genomics within and across species. However, even the current human reference genome assembly (GRCh38.p13) remains incomplete, affecting downstream analyses^7,8^. This problem is far worse for species with lower quality reference genomes or, in most cases, with no reference at all. Previously, it has been expensive to assemble genomes, so the luxury of a high-quality reference has been reserved for only the most well-studied species^2,3,9,10^. However, advances in sequencing technology are now making it possible to produce high-quality reference genomes cost effectively and at scale.

The first high-quality human^1^, mouse^2^, and zebrafish^4^ reference genomes were put together hierarchically, by independently assembling thousands of 50–300 kbp large insert clones using 500–1,000 bp Sanger sequencing reads and then stitching the clone sequences together using multiple mapping methods. Tremendous manual effort and software engineering were required, leading to costly, decade-long projects. Whole-genome shotgun methods, initially introduced for viruses^11,12^, were scaled up in the 1990s to include bacteria^13^ and later, with the addition of paired insert reads, to the fruit fly^9^, human^14^, pufferfish^15^, and mouse^16^. The advent in the late 2000’s of less expensive massive parallel sequencing^17,18^ enabled many laboratories to shotgun sequence a multitude of new genomes (e.g.^19,20^). However, the much shorter sequencing reads (30–150 bp) and typically short insert sizes (e.g. 1 kbp) resulted in poor assemblies that were often fragmented into thousands of pieces because most repetitive sequences longer than the read length could not be resolved. Up to 20% of the genes in these genomes could be either missing, particularly in GC-rich regions, truncated, or incorrectly assembled, resulting in annotation and other errors^21–25^. Such errors impact functional studies, and can require months of manual effort to correct individual genes and years to correct an entire assembly.

Genomic heterozygosity poses additional problems, because homologous haplotypes in a diploid genome are forced together into a single consensus by standard assemblers. In regions of greater allelic diversity, assemblers can correctly separate each allele but often interleave them in the final assembly, creating false duplications^7,22,26–28^. In attempts to avoid these issues, nearly all early eukaryotic reference genomes were based on highly inbred lines of model organisms (e.g. worm, fruit fly, and mouse). However, such inbreeding is impractical when sequencing thousands of species; it also does not completely eliminate haplotype heterozygosity.

To address this situation, the Genome 10K (G10K) consortium^5,6^ initiated the Vertebrate Genomes Project (VGP) with the ultimate aim of producing complete and error-free assemblies for all vertebrate species. This aim includes the completion and public availability of at least one high-quality, near error-free, near gapless, chromosome-level, haplotype phased, and annotated reference genome assembly for the 71,657 extant named vertebrate species (http://vgpdb.snu.ac.kr/splist/ VGP species list). A second aim is to use these genomes to address fundamental questions in biology, disease and biodiversity conservation.

Towards this end, we first evaluated genome sequencing and assembly approaches extensively on a test species, the Anna’s hummingbird (*Calypte anna*). We then deployed the best performing method across 15 additional species representing six major vertebrate classes, with a wide diversity of genomic characteristics, and based on lessons learned improved these methods further. Here we present these lessons with the first 16 VGP reference genomes, our optimal assembly approaches developed thus far, and a proposal for revised standards and terminology for measuring and classifying genome assembly quality.

### Breadth of data types

A female Anna’s hummingbird was chosen for benchmarking because it has a relatively small genome size (~1 Gbp), it is the heterogametic sex (having both Z and W sex chromosomes), and there is a previous annotated reference of the same individual built from short-reads^19^. We obtained 12 new sequencing data types, including both short and long reads (80 bp to 100 kbp) and long-range linking information (40 kbp to > 100 Mbp), generated with eight technologies: Illumina short reads (Illumina SR); Pacific Biosciences continuous long reads (PacBio CLR); Oxford Nanopore 1D and 2D ligation long reads (ONT); Bionano Genomics optical maps with either two restriction enzyme labelling (Bionano Opt1 or Opt2) or single direct labelling (Opt3); 10X Genomics linked read clouds (10XG linked SR); Hi-C read pairs produced by the proximity ligation protocols of Dovetail Genomics, Phase Genomics, and Arima Genomics; and NRGene DeNovoMAGIC (NRGene; **Supplementary Table 1**). To determine the most effective assembly approach, we benchmarked all technologies and associated assembly algorithms (**Supplementary Table 2**) in isolation and in many combinations and orderings. The final assemblies were compared to a manually curated reference of the most complete assembly.

### Long-read assemblies are more continuous

The product of an assembly process is a collection of genomic sequence reads assembled into *contigs*, which are then ordered and oriented into larger *scaffolds* with gaps between the contigs. The aim is to generate scaffolds and ultimately gapless contigs that represent complete chromosomes. To quantify assembly continuity, we used NG50, defined as the contig or scaffold size above which 50% of the total ‘genome’ size is found, with the expected genome size estimated statistically from the unassembled sequencing reads (**Methods**). Unlike N50, the NG50 measurement is independent of assembly size, and thus not influenced by data and algorithm quality. For diploid assembly approaches that separate the two haplotypes, we refer to the more continuous pseudo-haplotype assembly as the ‘primary’ and the other as the ‘alternate’. Throughout the benchmark, we used the primary contigs for scaffolding and measured NG50s.

We found that primary contigs assembled from long reads were ~30- to 300-fold longer than those assembled from short reads, regardless of data type combination or assembly algorithm used (**Fig. 1a** and **Supplementary Table 3**). The highest contig NG50 obtained with short reads was 0.169 Mbp, using linked reads, and most others were ~0.025 Mbp, including the prior short-read (SR) based reference (**Fig. 1a**). The highest contig NG50 obtained with long reads was 7.66 Mbp using a combination of CLR and ONT reads, or ~4.6 Mbp with CLR reads only (**Fig. 1a**). These findings indicate that given current sequencing technology and assembly algorithms, it is not possible to achieve high contig continuity with short reads alone.

**Fig. 1.**
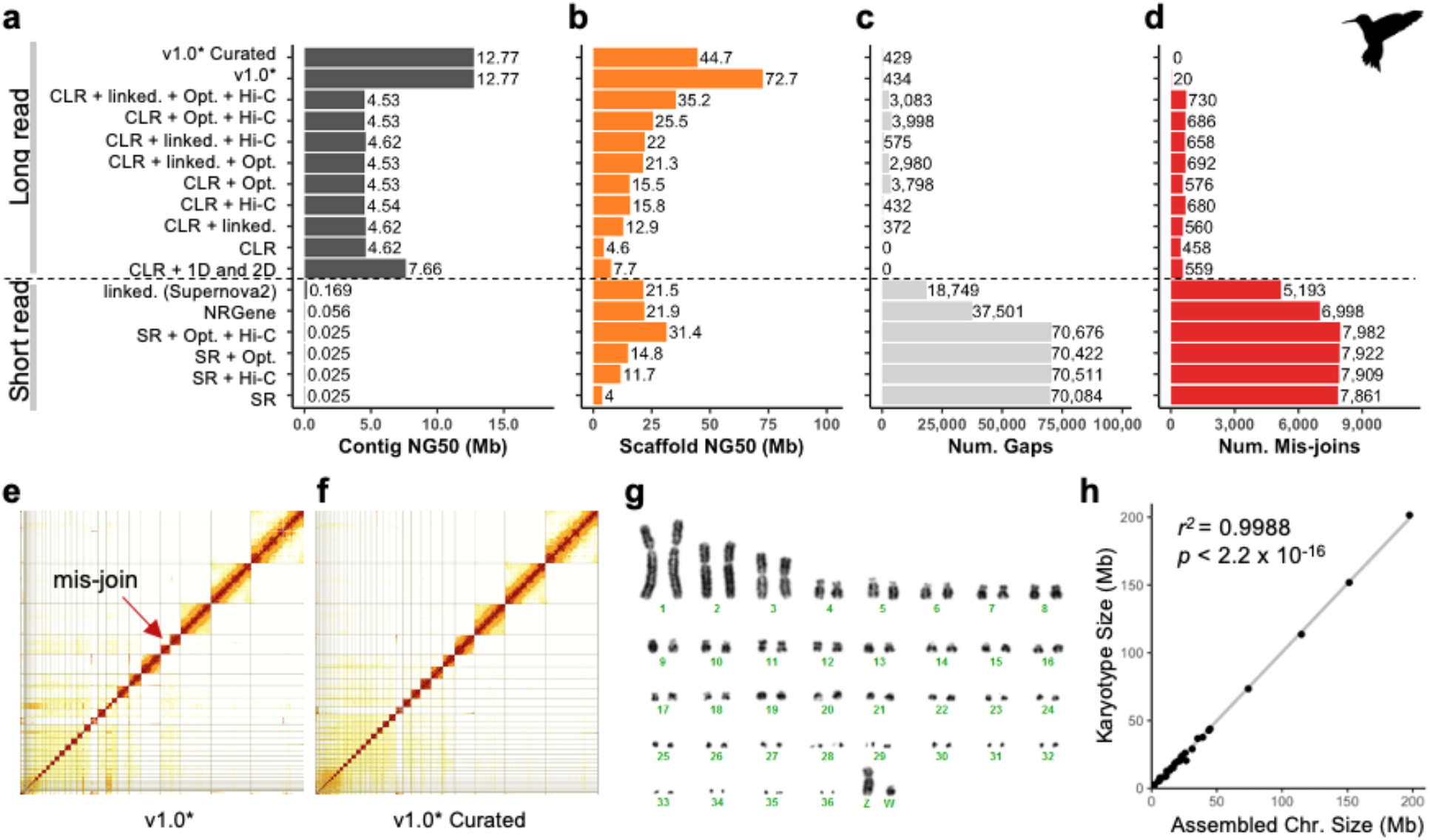
Comparative analyses of genome assemblies with various data types from Anna’s hummingbird. **a**, Contig NG50 values. **b**, Scaffold NG50 values. Only the primary pseudo-haplotype is shown for each assembly. **c**, Number of joins (gaps). **d**, Number of mis-join errors found in each assembly compared with the curated v1.0* assembly. *Assembly generated with software updates to FALCON^29^, following CLR + Linked + Opt. + Hi-C combination. **e-f**, Hi-C interaction heatmaps before and after manual curation, which identified 32 macro/micro-autosomes and ZW sex chromosomes. Grid lines indicate scaffold boundaries. Red arrow, example of a mis-joined scaffold that was corrected during curation. **g**, Karyotype of the identified chromosomes (n=36+ZW), consistent with Becak et al. (Anna’s Hummingbird)^30^. **h**, Correlation between estimated sizes (in Mbp) of chromosomes based on karyotype images in (**g**) and assembled scaffolds in **Supplementary Table 4** (bCalAnal). Abbreviations: VGP assembly v1.0 pipeline (v1.0), PacBio Continuous Long Reads (CLR), Bionano optical maps (Opt.), 10X Genomics linked reads (linked), Hi-C proximity ligation (Hi-C), Oxford Nanopore long reads (1D and 2D), NRGene paired-end Illumina reads (NRGene), and paired-end Illumina short reads (SR).

### Iterative assembly pipeline

We found that scaffolds generated with all three scaffolding technologies (i.e. linked reads, optical maps, and Hi-C) were ~50 to ~150% longer than those using one or two of the technologies, regardless of starting with short- or long-read based contigs (**Fig. 1b** and **Supplementary Table 3**). The highest scaffold NG50 obtained before curation and corrections to assembly algorithms was 35.2 Mbp, which was generated with long-read contigs followed by linked reads, optical maps, and Hi-C scaffolding. Despite similar scaffold continuity, the short-read only assemblies had from ~18,000 to ~70,000 gaps, whereas the long-read assemblies had substantially fewer, from ~400 to ~4,000 gaps (**Fig. 1c** and **Supplementary Table 3**).

During assembly curation, we discovered that the original version of FALCON^29^ software was producing inaccurate breaks in contigs at junctions between stretches of highly homozygous and highly heterozygous sequences of each haplotype (**Supplementary Note 1**). When we fixed this function in an updated FALCON release (**Supplementary Table 2**), the NG50 contig continuity nearly tripled to 12.77 Mbp with a concomitant increase in scaffold NG50 (* in **Fig. 1a,b**). Considering the final curated version of this assembly as correct (**Extended Data Fig. 1a**), we identified many mis-joins introduced by all automated contig and scaffolding methods (**Fig. 1d**). However, mis-joins in short-read based assemblies numbered from ~5,000 to 8,000, whereas those in long-read based assemblies numbered from only 20 to ~700. These mis-joins included chimeric joins and inversions not supported by the curated assembly, caused by large haplotypic differences between the assemblies or mis-assembly errors. After conducting the final curation step, which included contamination screening, correction of assembly errors, and making Hi-C based chromosome assignments (**Fig. 1e-f** and **Methods**), the final Anna’s hummingbird assembly had 33 scaffolds that closely matched the chromosome karyotype in number (33 of 36 autosomes plus sex chromosomes) and estimated sizes (~2 to ~200 Mbp; **Fig. 1g,h**). This assembly had from 1 to 30 gaps per autosome, and 73 and 92 gaps in the more heterogenous Z and W sex chromosomes, respectively (bCalAnn1 in **Supplementary Table 4**). From coverage analysis using Asset software^31^, we defined reliable blocks as two or more technologies that supported the chromosome assembly structure (**Methods**). Of the five autosomes with only 1 gap each, three (Chr 14, 15, and 19) had complete reliable block spanning support across the gap and entire scaffold, with no detectable issues of collapsed repeats (**Extended Data Fig. 1c**; bCalAnn1 in **Supplementary Table 4**), indicating that these scaffolds were near complete chromosome contigs. However, they were missing long arrays of the conserved vertebrate telomere repeats (AATCCC or TTAGGG) within 1 kb of their ends (**Extended Data Fig. 1c**; bCalAnn1 in **Supplementary Tables 4-5**), indicating that these chromosomes were missing the tail ends. All other autosomes had 2–10 regions of low coverage support or collapsed repeats, although some had defined telomeres, indicating that these chromosomes were also mostly complete.

During our studies on the Anna’s hummingbird genome, we also evaluated and improved Hi-C and other scaffolding, gap filling (**Supplementary Table 6**), and base-call polishing methods (**Supplementary Table 7**), as discussed further in **Supplementary Note 1**. Despite their increased continuity and structural accuracy compared to short-read-based assemblies, the CLR-based assemblies required at least two rounds of short-read consensus polishing to reach a base-level accuracy of 99.99% (1 error per 10 kbp, Phred^32^ Q40, **Supplementary Table 7**).

Based on the formula that gave the highest quality hummingbird genome (**Extended Data Fig. 1a**), we built an iterative assembly pipeline (v1.0): 1) generate primary pseudo-haplotype and alternate haplotype contigs with CLR using FALCON-Unzip^33^; 2) generate scaffolds with linked reads using Scaff10x^34^; 3) break mis-joins and further scaffold with optical maps using Solve^35^; 4) generate chromosome-scale scaffolds with Hi-C reads using Salsa2^36^; 5) fill in gaps and polish base-errors with CLR using Arrow (Pacific BioSciences); 6) perform two or more rounds of shortread polishing with linked reads using FreeBayes^37^; and 7) perform expert manual curation to correct potential assembly errors using gEVAL^38^ and through comparison with previous genome assemblies from the same and/or closely related species (**Fig. 2a, Extended Data Fig. 2a**). ONT data were not considered for further benchmarking due to practical issues concerning systematic base call errors, consistency, and scalability at the time (early 2017)^39^; however the technology has since improved in these areas^40^ and will be reconsidered in future phases of the VGP, as will PacBio’s recently released HiFi circular consensus sequencing (CCS)^41^.

**Fig. 2.**
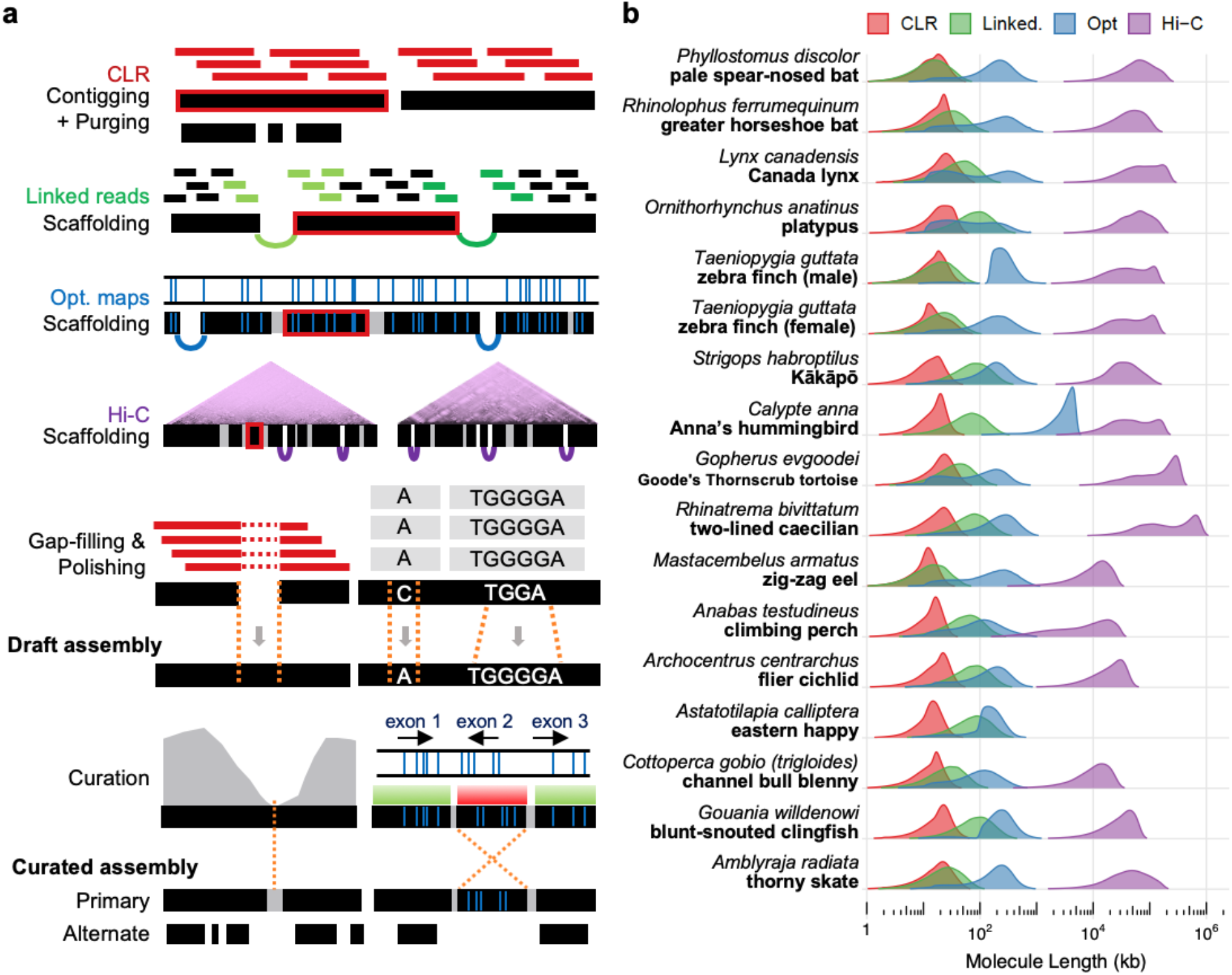
VGP assembly pipeline applied across multiple species. **a**, Iterative assembly pipeline of sequence data types (colored as in **b**) with increasing chromosomal distance. Thin bars, sequence reads; thick black bars, assembled contigs; black bars with space and arcing links, scaffolds; grey bars, gaps placed by previous steps; thick red border, tracking of an example contig in the pipeline. The curation step shows an example of a misassembly break identified by sequence coverage (grey, left) and an example of an inversion error (right) detected by the optical map. **b,** Intramolecule length distribution of the four data types used to generate the assemblies of 16 vertebrate species, weighted by the fraction of bases in each length bin (log scaled). Molecule length above 1 kbp was measured from read length for CLR, estimated molecule coverage for linked reads, raw molecule length for optical maps, and interaction distance for Hi-C reads, respectively (**Supplementary Methods**).

### VGP pipeline across vertebrate diversity

Based on our initial findings (**Fig. 1** and Korlach et al^22^), we set the goal to achieve minimum assembly metrics of: NG50 contig of 1 Mbp or greater; NG50 scaffold of 10 Mbp or greater; 90% or greater of the sequence assigned to chromosomal scaffolds, structurally validated by at least two independent lines of evidence; average base quality of Q40; and individual haplotypes assembled as complete and as correctly phased as possible.

We tested the above pipeline (**Extended Data Fig 1b**) for achieving these minimum metrics on vertebrate species spanning all major classes: four mammals; two additional birds; one reptile; one amphibian; six teleost fishes; and one cartilaginous fish (**Supplementary Tables 8-9**). These species were chosen: 1) to compare assemblies of simpler (birds) to more complex and repetitive genomes (amphibians and fishes); 2) to include those threatened (platypus) or critically endangered (kākāpō) of becoming extinct and having low heterozygosity due to small effective population sizes (Dussex, submitted); 3) to answer specific biological questions (e.g. in vocal learning birds and bats)^42,43^; 4) to compare with previous assemblies with available genetic linkage or FISH karyotype maps (zebra finch, platypus)^44,45^; 5) to contribute to collaborative projects with the VGP (e.g. Bat1K^46^); and 6) to take advantage of high quality samples and available funding within the VGP collaboration. For zebra finch, we used DNA from the male that was assembled previously^45^ and included a female zebra finch trio for benchmarking haplotype completeness and phasing accuracy of single-versus trio-based assembly methods, the latter for which sequenced reads from the offspring are binned by parental haplotypes before assembly^47^ (**Extended Data Fig. 2b**). For each species, the fragment length distribution of each data type was similar to those of the Anna’s hummingbird (**Fig. 2b**), with differences primarily influenced not by species but by the tissue type, the preservation method and collection/storage conditions (Dahn et al, in preparation).

While the VGP pipeline was in development, we assembled and submitted six of these species (four teleost fishes, the skate, the caecilian) to the public databases using a slightly different pipeline (**Fig. 3a, Supplementary Table 10** and **Supplementary Note 2**). Of all our submitted assemblies, 16 of 17 achieved the aspired continuity metrics (**Fig. 3a** and **Supplementary Table 11**). Among these 16 species, scaffold NG50 significantly correlated with genome size (**Fig. 3b**), suggesting that larger genomes tend to have larger chromosomes. On average, 98.3% of the assembled bases were supported by at least two platforms, with NG50 ranging from 2.3 to 40.2 Mbp (**Fig. 3a and Supplementary Table 11**). Completeness of the genome assembled measured from *k-mers* indicated 87.2 to 98.1%, with less than 4.9% of the *k-mers* duplicated consistent with BUSCO results (**Extended Data Fig. 3 and Supplementary Tables 11-12**). For consistency, when evaluating factors affecting continuity and assembly errors across species in our analyses below, we used data from applying the same VGP pipeline.

**Fig. 3.**
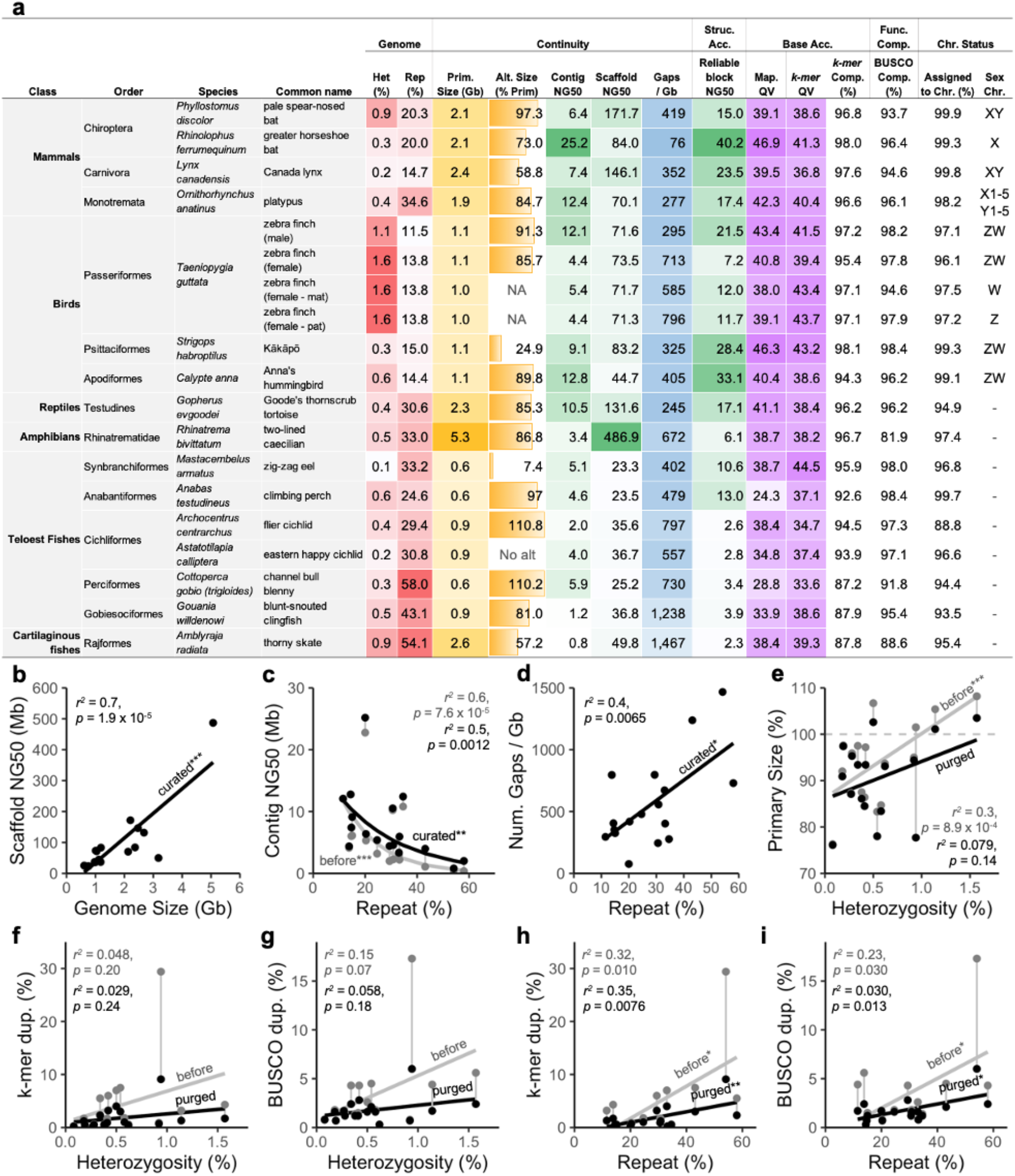
Impact of repeats and heterozygosity on assembly quality. **a**, Summary metrics of the curated and submitted vertebrate species assemblies. Color shading indicates degree of heterozygosity or repeats (red), primary assembly sizes and relative size of alternate haplotypes (orange), continuity measures (green), gaps (blue), and base call accuracy (purple). indicates that the sex chromosomes were not found or are not known. Accessions are available in **Supplementary Table 10**. **b**, Correlation between scaffold NG50 and genome size of the curated assemblies. **c**, Nonlinear correlation between contig NG50 and repeat content, before and after curation. **d**, Correlation between number of gaps per Gbp assembled and repeat content. **e**, Correlations between primary assembly size relative to estimated genome size (y-axis) and genome heterozygosity (x-axis), before and after purging false duplications. Assembly sizes above 100% may indicate false heterotype duplications and those below may indicate collapsed repeats. **f-g**, Correlations between genome duplication rate and specific conserved genes (BUSCO vertebrate gene set) with genome heterozygosity before and after purging false duplications. **h-i**, Similar analyses as f-g, but with whole genome repeat content before and after purging false duplications. Genome size, heterozygosity, and repeat contents were estimated with GenomeScope^47^ using 31-mers. Whole genome duplication rates were estimated with Merqury^48^ using 21-mers. Asterisks in **b-i** indicate degrees of statistical significance of the correlation coefficient.

### Repeats affect continuity

When applying the same standard VGP assembly pipeline (**Fig. 2a**) across species, all but two of the 17 assemblies exceeded the aspired continuity metrics before curation (**Supplementary Table 13**). However, contig NG50 showed a significant exponential decrease with increasing repeat content (**Fig. 3c**). The two genomes that did not initially meet the minimum NG50 of 1 Mbp contig size, thorny skate and channel bull blenny, contained the highest repeat contents (54.1% and 58.0%, respectively), substantially higher than those of the other fishes (>24%), mammals (15– 35%), and birds (10–15%) (**Fig. 3c** and **Supplementary Table 13**). Consequently, after scaffolding and gap filling, we observed a significant positive correlation between repeat content and the number of gaps per Gbp assembled, such that genome assemblies of species with fewer repeats had fewer gaps (**Fig. 3d**). For example, the kākāpō parrot assembly with 15% repeat content had ~325 gaps/Gbp, including zero gaps in two of its 26 chromosome-level scaffolds (Chrs 16 and 18), with no evidence of collapsed repeats or low support, suggesting that they might represent complete chromosome contigs (bStrHab1 in **Supplemental Table 4**); the greater horseshoe bat assembly with 20% repeat content also had no gaps in two of its 28 chromosome-level scaffolds (Chrs 8 and 26), with full support (mRhiFer1 in **Supplementary Table 4**). By contrast, the thorny skate with 54% repeat content had ~1,400 gaps/Gbp, with none of its 49 chromosomal-level scaffolds containing fewer than eight gaps, and all with some regions containing collapsed repeats or low support (**Fig. 3a**; sAmbRad1 in **Supplementary Table 4**). These findings quantitatively validate the hypothesis that repeat content is a major factor that impacts the ability to produce highly continuous assemblies.

### False duplications are common assembly errors

During curation, we discovered that one of the most common assembly errors was the introduction of false duplications. In biological investigations, these false duplications can be misinterpreted as exon, whole gene, or large segmental duplications that occur either in tandem on the same chromosome or as translocated to a different chromosome. We observed two types of false duplications: 1) false ‘*heterotype duplications*’; and 2) false ‘*homotype duplications*’ (**Box 1**). Heterotype duplications occur in regions of increased sequence divergence between paternal and maternal haplotypes, where separate haplotype contigs are incorrectly placed in the primary assembly. During scaffolding, these contigs are either brought together with a gap in between or remain separate with one haplotype residing as a smaller contig (**Box 1**). These duplications occurred in our assemblies despite setting FALCON^29^ parameters to resolve up to 10% haplotype divergence. Heterotype duplication sequence-length varied depending on heterozygosity level. For example, in the female zebra finch, we found a falsely duplicated heterozygous sequence over 1 Mbp in length during curation (**Extended Data Fig. 4a**). This zebra finch individual in particular had the highest heterozygosity (1.6%) relative to all other genomes (0.1-1.1%). Homotype duplications occur near contig boundaries or under-collapsed sequences caused by sequencing errors (**Box 1**). Similar to heterotype duplications, the duplicated sequences are either placed next to each other in a scaffold with a gap or remain as separate contigs (**Box 1**). In our assemblies, homotype duplication length at contig boundaries was approximately the length of sequence read lengths (**Extended Data Fig. 4b**).

#### Box 1. False duplication mechanisms in genome assembly

Here, we describe in more detail the mechanisms we noted behind false duplications in genome assembly. A **heterotype duplication** occurs when more divergent sequence reads from each haplotype A (blue) and B (red) (i.e. maternal and paternal) form greater divergent paths in the assembly graph (bubbles), while nearly identical homozygous sequences (black) become collapsed (**Fig. B1a**). When the assembly graph is properly formed and correctly resolved (green arrow), one of the haplotype specific paths (red or blue) is chosen for building a ‘primary’ pseudo-haplotype assembly and the other is set apart as an ‘alternate’ assembly. When the graph is not correctly resolved (purple arrow), one of four types of patterns are formed in the contigs and subsequent scaffolds (numbered 1-4 in **Fig. B1a**):

1. Separate contigs: Both contigs are retained in the primary contig set, an error often observed when haplotype-specific sequences are highly diverged.
2. Flanking contigs: The assembly graph is partially formed, connecting the homozygous sequence of the 5’ side to one haplotype (blue) and the 3’ side to the other haplotype (red).
3. Partial flanking contigs: Only one haplotype (blue) flanks one side of the homozygous sequence.
4. Failed connecting of contigs: All haplotype sequences fail to properly connect to flanking homozygous sequences.

Depending on the supporting evidence, the scaffolder either keeps these haplotype contigs on separate scaffolds or brings them together on the same scaffold, often separated by gaps.

A **homotype duplication** occurs where a sequence from the same genomic locus is duplicated in extra copies. There are at least two types of homotype duplications (**Fig. B1b**):

1. Overlapping sequences at contig boundaries: In current overlap-layout-consensus assemblers, branching sequences in assembly graphs that are not selected as the primary path have a small overlapping sequence (purple), dovetailing to the primary path where it originated a branch. The size of the duplicated sequence is often the length of a corrected read. Subsequent scaffolding results in tandemly duplicated sequences with a gap between.
2. Under-collapsed sequences: Sequencing errors in reads (red x) randomly or systematically pile up, similar to diverged haplotype sequences, forming under-collapsed sequences. Subsequent duplication errors in the scaffolding are similar to the heterotype duplications.

Purge_haplotigs^27^ align sequences to themselves to find a smaller sequence aligning fully to a larger contig or scaffold, and removes heterotype duplication types 1, 3, 4. Purge_dups additionally uses coverage information to detect heterotype duplication type 2 and homotype duplications. Shorter contig size and coverage information are often not enough to distinguish false duplications from allele-specific duplications or segmental duplications, and thus are at risk of being removed.

**Fig. B1.**
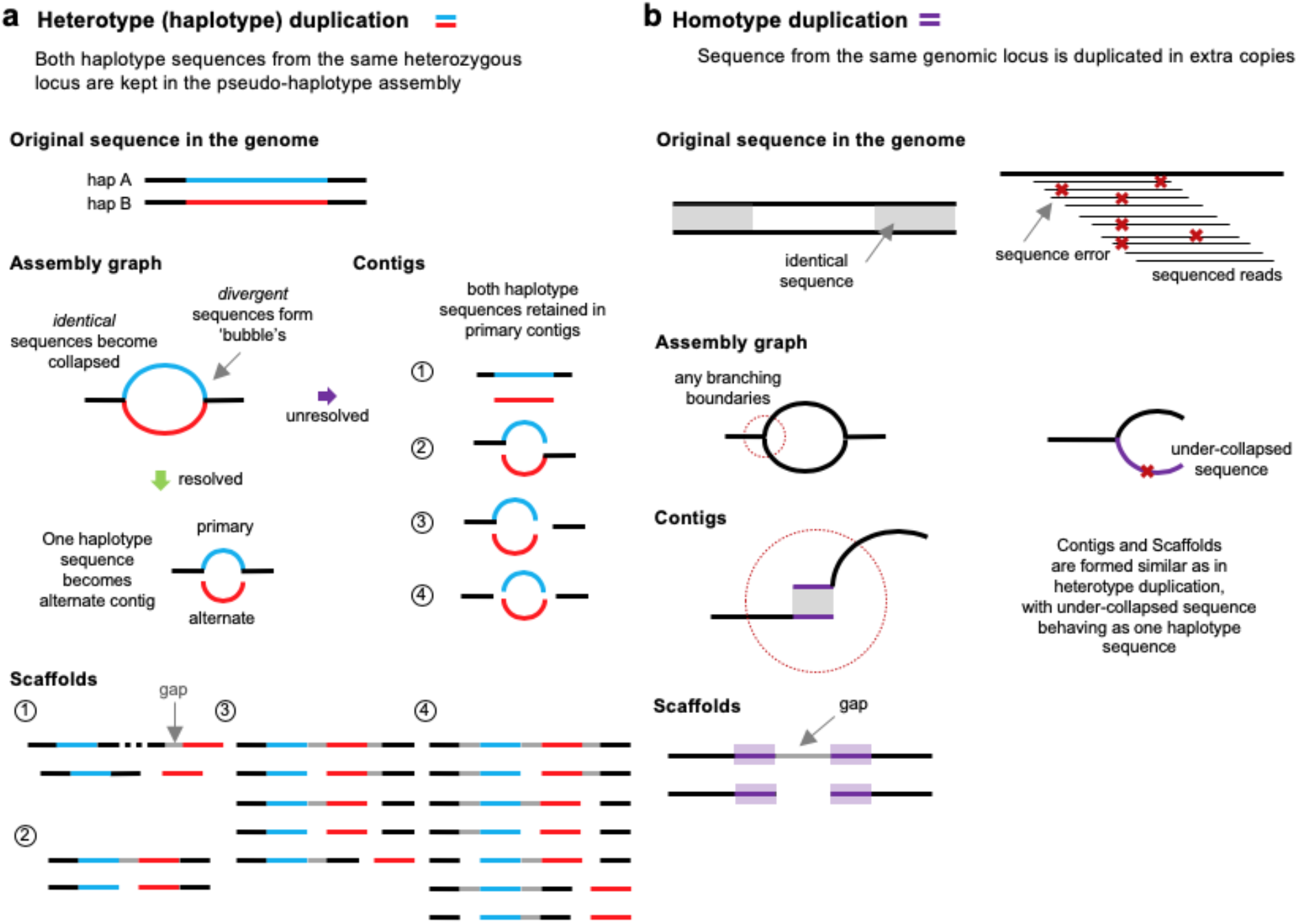
Types of artificial false duplications. **a**, False heterotype (haplotype) duplications. Haplotype specific sequences marked as red and blue are retained in the primary assembly as a false tandem duplication or as separate contigs. **b**, False homotype duplications. Sequences from the same genomic locus are erroneously duplicated as extra copies (purple). Left, duplicated sequences that occur at contig boundaries. Right, unresolved sequences forming duplications similar as heterotype duplications.

We identified and removed false duplications during curation using read coverage, self-alignments, transcript alignments, optical map alignments, Hi-C maps, and *k-mer* profiles (**Extended Data Fig. 4**). We distinguished the two types of duplications by: 1) haplotype specific variants in reads aligning at half coverage to each heterotype duplication; 2) differing consensus quality that resulted from read coverage fluctuations when aligning reads to homotype duplications; and 3) *k-mer* copy number anomalies in which homotype duplications were observed in the assembly with more than the expected number of copies.

Consistent with these observations, prior to purging these duplications, the primary assembly genome size was correlated positively with the estimated percent heterozygosity, such that more heterozygous genomes tended to have assembly sizes bigger than the estimated haploid genome size (**Fig. 3e**). Similarly, the extra duplication rate in the primary assembly as measured using *k-mers^48^* or conserved vertebrate BUSCO orthologous genes^49,50^ varied from 0.3% to 30% with a correlation trend to heterozygosity (**Fig. 3f-g** and **Supplementary Table 13**). There was also a stronger correlation of apparent false gene duplication rates with the overall repeat rate in the assemblies (**Fig. 3h-i**).

To remove these false duplications, we initially used purge_haplotigs^27^, which removed retained false duplicated contigs that were not scaffolded (**Box 1**, incorporated in VGP v1.0–1.5). Later, we developed purge_dups^51^ to remove both these fully duplicated contigs and end-to-end duplicated contigs within scaffolds (**Box 1**, incorporated in VGP v1.6), resulting in less manual curation. After applying these tools, the primary assembly sizes, the *k-mer* duplication rate, and the BUSCO gene duplication rate were all reduced, and their correlations with heterozygosity and repeat content were also reduced or eliminated (**Fig. 3e-i**). These findings indicate that properly phasing haplotypes and obtaining high consensus sequence accuracy are essential for preventing false duplications and associated biologically false conclusions.

### Curation is important for generating a high-quality reference

Each of the automated scaffolding methods introduced 10s to 1,000s of unique joins and breaks in contigs and scaffolds (**Supplementary Table 14**). Depending on species, the first scaffolding step with linked reads introduced ~50–900 joins between CLR generated contigs. Optical maps introduced a further ~30–3,500 joins, followed by Hi-C with ~30–700 more joins, and each identified up to several dozen joins that were inconsistent with the previous scaffolding step. The subsequent manual curation, considering all data types, resulted in an additional 7,262 total interventions for the 19 genome assemblies (2,496 breaks, 2,492 joins, 2,274 false duplication removals) or 236 interventions per Gbp of sequence (81 breaks, 81 joins, 74 false duplication removals/Gbp; **Supplementary Table 15**). Synteny analyses, when the same or a close relative species’ genome assembly was available, identified putative chromosomal breakpoints between them (**Supplementary Table 15**). The necessity of these interventions indicate that even with current state-of-the-art assembly algorithms, curation is essential for completing a high quality reference assembly and, perhaps even more importantly, for providing iterative feedback to improve assembly algorithms; for example, the FALCON improvements noted above and a new version of Salsa resulted directly from our curator’s feedback (**Supplementary Note 1**).

### Hi-C is comparable to karyotype mapping

Similar to the curated Anna’s hummingbird assembly, most large assembled scaffolds of all species spanned entire chromosomes, as shown by the relatively clean Hi-C heatmap plots across each scaffold after curation (**Extended Data Fig. 5**) and the presence of telomeric repeat motifs on some scaffold ends (**Supplementary Table 5**). To test the validity of these results, we performed chromosome homology alignments between the new VGP assemblies and the previous reference assemblies of the same species that used karyotype mapping approaches to scaffold and assign chromosomes. In our VGP zebra finch assembly, all inferred chromosomes were consistent with the genetic linkage map, except for chromosomes 1 and 1B, which were separated in the Sanger-based reference genome (**Extended Data Fig. 6a**). Their join in the VGP assembly was supported by both single-molecule CLR reads and optical maps through the junction. We also corrected nine inversions in smaller chromosomes of the prior zebra finch assembly and filled several large gaps at the ends of the larger chromosomes. In the platypus, we identified 18 differences in 13 scaffolds between the VGP assembly using Hi-C and the previous Sanger-based assembly anchored to chromosomes using FISH physical mapping (**Extended Data Fig. 6b; Supplementary Table 16**). Of these 18, seven were supported by both CLR and optical maps in the VGP assembly, so their chromosomal organization was reassigned. The remaining 11 were only supported by Hi-C. This comparison also revealed many very large (~1–30Mb) gaps of missing sequence in the previous platypus chromosome assignments that were filled by the VGP assembly, as well as many corrected inversions (**Extended Data Fig. 6b**). Our analyses also identified chromosomes that were not seen in previous assemblies, including two additional chromosomes in the zebra finch (Chrs 29 and 30) and six in the platypus (Chrs 8, 9, 16, 19, 21, and X4) (**Extended Data 6a,b**)(Zhou, submitted). Prior genome assemblies not assisted by genetic linkage or FISH karyotype mapping had more issues. For example, the prior short-read assembly of the Anna’s hummingbird was highly fragmented, despite being scaffolded with seven different Illumina libraries spanning a wide range of insert sizes (0.2-20 kbp). In this prior assembly, even the longest 20 scaffolds were small compared to the VGP Hi-C-based chromosome scaffolds (**Extended Data Fig 6c**). The climbing perch chromosomes were even more fragmented in the prior assembly and also had large gaps of missing sequence (**Extended Data Fig. 6d**). On average, after Hi-C scaffolding 97% (±3 s.d.) of all VGP assembled bases were assigned to chromosomes (**Fig. 3a**) compared with 76% in the prior zebra finch and 32% in the prior platypus references.

From these analyses, we conclude that our long-read assemblies scaffolded with Hi-C and optical mapping provide chromosome counts and assignments that are comparable or better in accuracy than genetic linkage or FISH physical mapping. We believe this comparable or better accuracy is due to the high sampling rate of Hi-C pairs across the genome, which provides higher resolution for identifying errors in the initial assembly and determining chromosome organization compared with linkage or FISH probes. Nonetheless, visual karyotyping remains valuable for complementary validation of chromosome count and structure^52^.

### Trios help resolve haplotypes

Building on the previously described trio-binning approach using offspring and parental data^53^, we assembled trio-based zebra finch contigs into separate maternal and paternal chromosome-level scaffolds (**Fig. 4a**) using our VGP trio pipeline (**Extended Data Fig. 2b**). Compared with the non-trio assembly, the trio version had fewer (7- to 8-fold) false duplications (*k-mer* and BUSCO dups in **Supplementary Table 11-12**), well-preserved haplotype-specific variants (*k-mer* precision/recall 99.99/97.08%), and higher base call accuracy, exceeding Q43 for both haplotypes (**Fig. 3a**). The trio-based zebra finch assembly was the only assembly with nearly perfect (99.99%) separation of maternal and paternal haplotypes, determined based on *k-mers* specific to each^48^. From this trio-based assembly, we identified structural variants between the two haplotypes, including inversions of 4.5 to 12.5 Mbp on chromosomes 5, 11, and 13 that we originally misinterpreted as inversion assembly errors in the non-trio version (**Extended Data Fig. 7**). Moving forward, the VGP is prioritising collection of mother-father-offspring trios or singleparent-offspring duos where possible to assist with diploid assembly and phasing, as well as development of improved methods for assembly of diploid genomes in the absence of parental genomic data, as described in a companion study^54^.

**Fig. 4.**
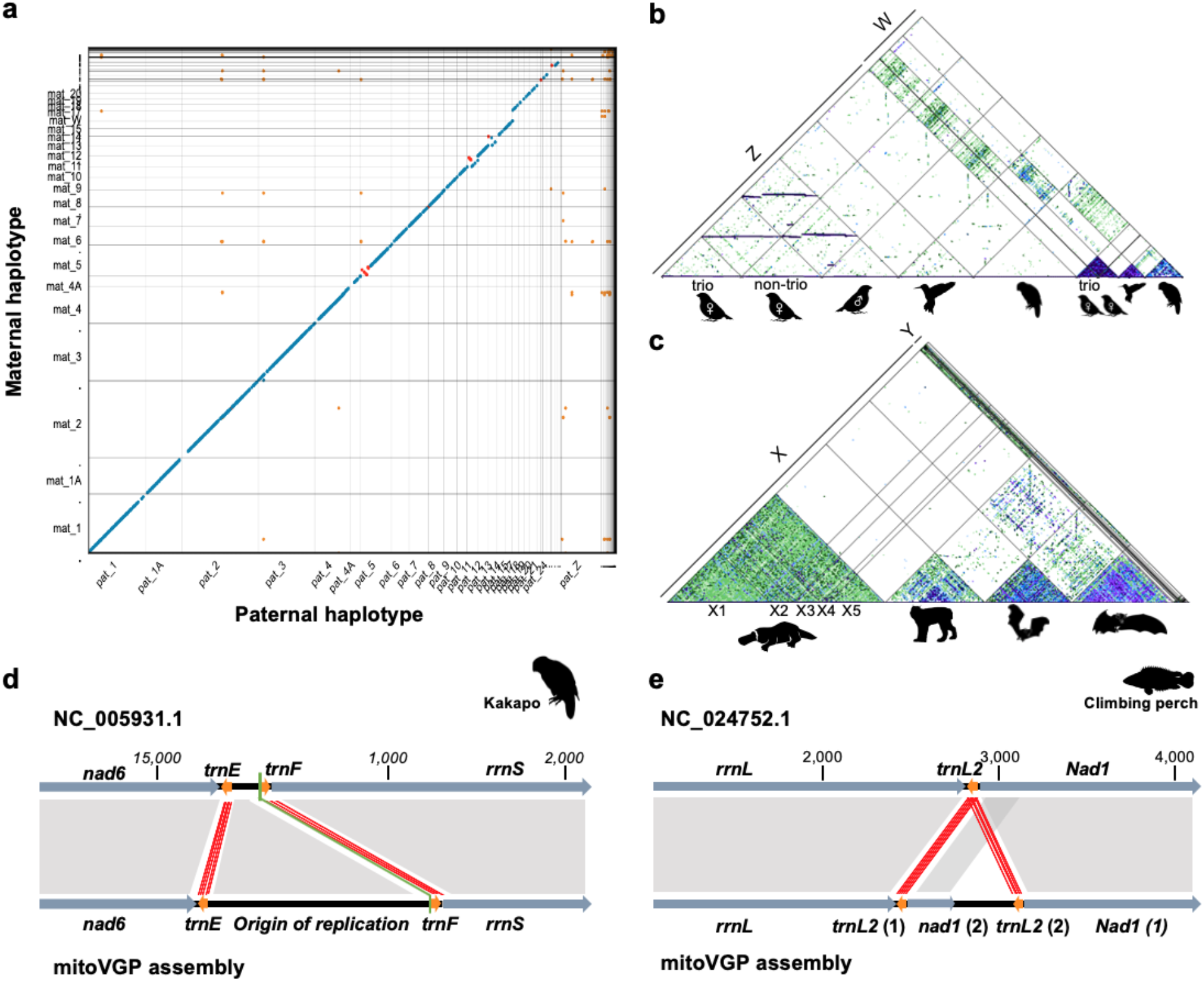
Haplotype resolved sex chromosomes and mitochondrial genomes. **a**, Alignment scatterplot, generated with MUMmer NUCmer^56^, of maternal and paternal chromosomes from the female zebra finch trio-based assembly. Colors denote the same orientation (blue), inversion (red), or repeats (orange) between haplotypes. Note, the paternal Z sex chromosome is highly divergent from the maternal W and so remained mostly unaligned. **b**, Alignment scatterplot of Z and W sex chromosomes across the three bird species assembled in this study, approximated with MashMap2^57^. Segments of 300 kb (green), 500 kb (blue), and 1 Mb (purple) are shaded darker with higher sequence identity, with a minimum of 85%. Relative to Z, the smaller size and higher repeat content of the W is clearly visible. **c**, X and Y chromosome segments of the monotreme (platypus) and placental mammals (Canada lynx, pale spearnosed bat, and greater horseshoe bat), using the same approach as in (b) and showing a higher density of repeats within the mammalian X than the avian Z. **d,** The VGP mitochondrial genome assembly of the kākāpō, reveals a missing repetitive sequences (adding 2,232 bp) in the origin of replication region of the current so-called complete reference, containing a 83 bp repeat unit. **e**, The climbing perch VGP mitochondrial genome assembly showing a duplication of *trnL2* as well as a partial duplication of *Nad1*, which was absent from the prior reference. Orange arrows and red lines, tRNA genes and their alignments; dark grey arrows and grey shading, all other genes and their alignments; black, noncoding regions; green line, conventional starting point of the circular sequence.

### Polishing helps but also introduces errors

Before polishing, the per-base assembly accuracy of our assemblies was in the Q30–35 range, short of our Q40 goal. The most common errors were short indels from inaccurate consensus calling during CLR contig formation, which resulted in amino acid frameshift errors in gene annotation. Using our combined approach of Arrow polishing with long reads and FreeBayes^37^ polishing with short reads of both primary and alternate sequences together, we polished between 82–99.7% of the primary and ~91.3% of the alternate assembly bases across the 17 assemblies (**Supplementary Table 17**). Of the remaining unpolished sequence, one haplotype was sometimes reconstructed at substantially lower quality than the other haplotype due to most reads being attracted to the higher quality haplotype during mapping before polishing. This resulted in one haplotype, mostly in the primary assembly, being transformed into a mosaic during polishing while the second haplotype remained unpolished (**Extended Data Fig. 8a**). We observed these mosaic haplotypes in the more homozygous regions that tended to be collapsed by FALCON-Unzip^33^. False duplications had similar effects, because the duplicated sequence acted as an attractor during the read mapping phase of polishing, resulting in uneven sequence depth and quality. From the zebra finch trio Z chromosome assembly, we confirmed that haplotype switches were erroneously introduced when polishing with unbinned short reads (**Extended Data Fig. 8b**). By using parental markers, we excluded read pairs from the wrong haplotype and performed one more round of polishing, which fixed the switch errors. These findings indicate that both sequence read accuracy and careful haplotype separation are important for producing accurate assemblies.

### Sex chromosome assemblies

Vertebrate sex chromosomes are notoriously difficult to assemble due to their greater divergence in the heterogametic sex relative to autosomes and their high repeat content^55^. When sequencing the heterogametic sex, we identified sex chromosomes in the avian and mammalian genomes based on half coverage, homology alignments with known sex chromosomes in other species, and presence of sex chromosome-specific genes (**Methods**). We successfully assembled both sex chromosomes (Z, W) for all three avian species (**Extended Data Fig. 5**), generating the first assembly of the W chromosome in vocal learning birds, a sexually dimorphic trait in these species (**Fig. 4b**). We assembled the X and/or Y in placental mammals (Canada lynx and two bat species), and for the first time all 10 sex chromosomes in the platypus (5X/5Y, **Fig. 4c**)(Zhou, submitted). Completeness and continuity of the zebra finch Z and W chromosomes were further improved by the trio-based assembly (**Fig. 4b**). However, while curating several trios, we found that in regions of low divergence between shared parental homogametic sex chromosomes (i.e. the X or Z), a small fraction of offspring CLR data from this chromosome was mis-assigned to the wrong haplotype. This mis-alignment resulted in a duplicate, low-coverage offspring X or Z assembly in the paternal (for mammals) or maternal (for birds) haplotype respectively, which required removal during curation. We are working on methods to improve the binning accuracy for an automatic resolution on this issue going forward.

### Mitochondrial genomes and gene duplications

Mitochondrial (MT) genomes, expected to be 11–28 kbp in size^58^, were initially found only in six assemblies (**Supplementary Table 18**). We found MT-derived raw reads for all other genomes, but they failed to assemble due in part to minimum read length cutoffs of the starting contig assembly. Further, if an assembled MT genome was not present during nuclear genome polishing, the raw MT reads were attracted to nuclear MT sequences (NuMTs) causing them to incorrectly convert to the organelle MT sequence (**Extended Data Fig. 8c**). To address these issues, we developed a reference-guided pipeline for selecting and assembling the MT-derived reads separately (**Extended Data Fig. 2c** and **Methods**). To prevent NuMTs from being erroneously replaced by MT sequence, the assembled MT genome was added back to the nuclear scaffolds during assembly polishing to act as an attractor for the MT reads (VGP v1.6). With these improvements, we reliably assembled 16 of 17 MT genomes (**Supplementary Table 18**). The high quality of these assemblies allowed us to discover previously unknown variation, including a 2 kbp repeat (repeat unit 83 bp) within the control region in the kākāpō (**Fig. 4d**) and a duplication of *Nad1* and *trnL2* genes in the climbing perch (**Fig. 4e**). These duplications were verified with single-molecule CLR reads spanning the duplication junctions and single reads comprising the entire MT genome. Their absence in the previous MT references^59,60^ are likely due to the inability of Sanger or short reads to correctly resolve the duplications. More details on the MT-VGP pipeline and new biological discoveries are reported in a companion study(Formenti et al, in preparation).

### Improvements to mapping and annotation

In comparison to previous intermediate-read Sanger (zebra finch and platypus) and short-read Illumina (Anna’s hummingbird and climbing perch) assemblies, we added ~42–176 Mbp of assembled sequence, placed 57.6 Mbp (zebra finch) to 1.8 Gbp (platypus) of previously unplaced sequence within chromosome-assigned scaffolds, corrected from ~7,800 to 64,000 mis-joins, and closed from 55,177 to 193,137 gaps per genome in the new VGP assemblies (**Supplementary Table 19**). Consistent with these differences, both transcriptome RNA-Seq data and genome ATAC-Seq data aligned with ~5 to 10% greater map-ability to our new VGP assemblies (**Fig. 5a**). Using the same input RNA-Seq, transcripts, and protein evidence to perform the EBI Ensembl and NCBI RefSeq annotation pipeline, we annotated ~15,000 to ~27,000 protein coding genes in both annotations (**Supplementary Table 20** and **Methods**). The new VGP assemblies had 5,434 to 14,073 more protein coding transcripts identified, with 94.1 to 97.8% fully supported (**Fig 5b left**). The VGP assemblies also had 1,527 to 5,373 fewer partial coding genes (**Fig. 5b right**), supporting the structural accuracy of the assemblies. We found more ortholog coding genes to human and/or the SwissProt protein database, and fewer transcripts required corrections to compensate for a premature stop codon or frameshifting indel error (**Extended Data Table 1**). The total number of genes annotated went down in the VGP assemblies (**Extended Data Table 1**), which we believe is because the VGP assemblies have fewer false duplications (0.5 to 1.7%) compared with their previous counterparts (5.1 to 12.3%; except for the climbing perch at 0.4 vs 1.2%; **Supplemental Table 19**). Further supporting these results, the VGP assemblies had 6 to 13% higher *k-mer* completeness (95±3.5% average; **Fig. 3a**) compared with the prior assemblies (88±4.3% average; **Supplementary Table 19**). These findings clearly demonstrate that assembly quality has critical implications on subsequent annotations and functional genomics.

**Fig. 5.**
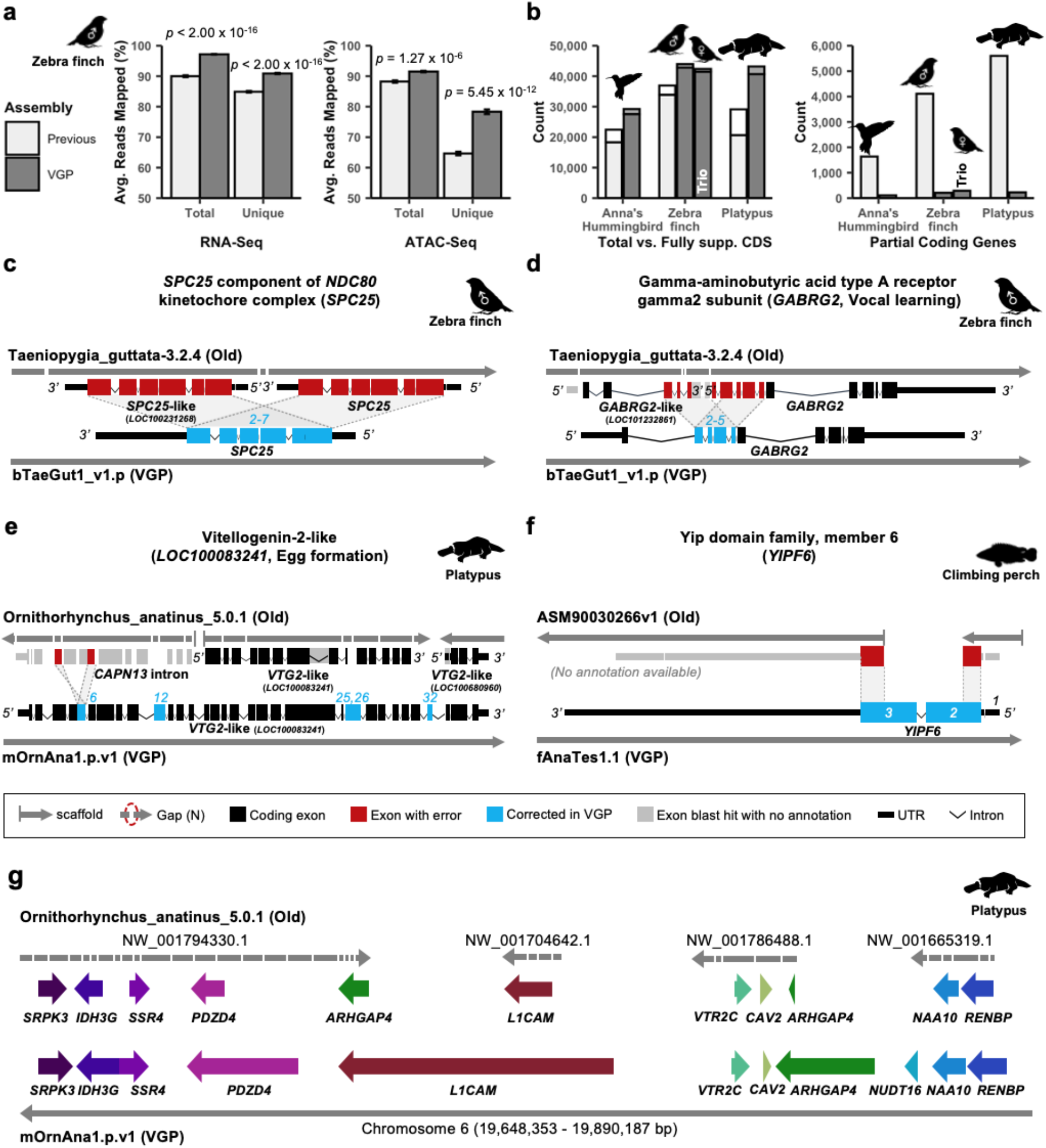
Improvements to alignments and annotations in VGP assemblies relative to prior references. **a**, Average percentage of RNA-Seq transcriptome (n=44) and ATAC-Seq genome (n=12) reads (± SEM) that align to the previous (Taeniopygia_guttata-3.2.4 2008) and new VGP (bTaeGut1_v1.p 2019) zebra finch assemblies. Unique reads mapped to only one location in the assembly. Total is the sum of unique- and multi-mapped reads. **b**, Total number of coding sequence (CDS) transcripts and portion fully supported (left) and partial RefSeq coding genes (right) annotated in the previous and new VGP assemblies of the Anna’s hummingbird, zebra finch, and platypus using the same input data (accessions in **Supplementary Table 19**). The VGP assemblies had higher numbers of transcripts and lower numbers of partial coding genes. **c–f**, Example assembly errors and associated annotation errors in previous (old) reference assemblies corrected in the new VGP assemblies. Both haplotypes of *SPC25* (**c**) were erroneously duplicated on two different contigs, annotating one as *SPC25-like*. The 5’ end part of *GABRG2* (**d**) was erroneously annotated as a separate *GABRG2-like* protein coding gene, due to false duplication of exons 2–5. The *VTG2* gene (**e**) was annotated on 3 scaffolds as part of 3 separate genes, two *VTG2-like* and an intron of CANP13. *YIPF6* (**f**) was partially missing in the previous assembly due to truncated exon sequences at the scaffold ends. No gene annotation was available for the previous climbing perch assembly. **g**, Gene synteny around the *VTR2C* receptor in the platypus shows completely missing genes (*NUDT16*), truncated and duplicated *ARHGAP4*, and many gaps in the prior Sanger-based assembly compared with the filled in and expanded gene lengths in the new VGP assembly. All examples shown here showed support from at least two technologies across these regions, while the prior assemblies showed hallmarks of misassembly.

Example cases include whole false gene duplications, as detected by comparing the latest RefSeq annotation of the prior reference and the VGP assemblies, such as the BUSCO gene *SPC25* of the NDC80 kinetochore complex^61^ of the prior zebra finch Sanger-based assembly^45^, of which only one copy is present in the VGP assembly (**Fig. 5c** and **Supplementary Table 21**). The GABA receptor *GABRG2* with specialized gene expression in vocal learning circuits^62^ had a partial tandem duplication of four of its 10 exons, resulting in an annotated partial false gene duplication as two adjacent genes (*GABRG2 and GABRG2-like*) in the Sanger-based zebra finch assembly (**Fig. 5d**). The vitellogenin-2 (*VTG2*) gene, an important component of egg-yolk in all egg-laying species^63^, was distributed across 14 contigs in three different scaffolds, two that received two corresponding *VTG2-like* gene locus (LOC) annotations and the third that was mistakenly included as part of the intron of another gene (*Calpain-13*) and that had an inverted non-tandem false exon duplication (red), all together causing false amino acid sequences in five exons (blue), in the prior Sanger-based platypus assembly^43^ (**Fig. 5e**). The BUSCO *YIPF6* gene, associated with inflammatory bowel disease^64^, was split between two different scaffolds and, thus, not annotated and presumed to be a gene loss in the prior Illumina-based climbing perch assembly^65^ (**Fig. 5f**). Each of these genes is now present on one long contig, with no gaps and no false gene-region gains or losses in the VGP assemblies, validated in reliable blocks with support from two or more sequencing platforms (**Supplementary Table 21**). Going beyond individual genes to gene synteny, a 10-gene window surrounding the vasotocin receptor 2C (*VTR2C; aka AVPR2*), a gene involved in blood pressure homeostasis and brain function^66,67^, was split into 34 contigs on four scaffolds, one containing a false haplotype duplication of *ARHGAP4*, in the prior Sanger-based platypus assembly (**Fig. 5g**). In our VGP assembly, all 11 genes were now in one long, gapless contig within a 220 Kbp region on chromosome 6. Further, the gene models for 8 of 11 genes in this region were remarkably increased in size due to corrections and additions of previously unknown missing sequences in the earlier assembly. The chromosomal region containing the 11 genes and many more was more GC-rich (54%) compared with the whole of chromosome 6 (46%). Additional details and mechanisms behind hundreds to thousands of such false gains and losses in previous reference assemblies that are now correct in the VGP assemblies are detailed in three companion studies(Ko et al, in preparation; Kim et al, in preparation; Theofanopoulou et al, 2020, in press).

### New biological discoveries

Our companion and other studies reveal that the higher quality VGP assemblies of the 16 species enabled novel, or more accurate, biological discoveries that were not possible with lower quality drafts. These include: 1) more accurate gene synteny across species, leading to a better understanding of the evolution and thus universal nomenclature for the vasotocin (a.k.a. vasopressin) and oxytocin ligand and receptor gene families across vertebrates (**Fig. 5g**)(Theofanopoulou et al, 2020, in press,); 2) greater understanding of the evolution of the carbohydrate 6-O sulfotransferase gene family, whose members encode enzymes that modify secreted carbohydrates^68^; 3) a genome-scale phylogeny that better resolves the relationships between bats and other mammals, and for discovering novel changes in genes involved in immunity, life span, and microRNAs, including reveleance for the current COVID-19 pandemic, in the first Bat1K study^46,69^; 4) discovery that deleterious mutations have been purged from the last surviving isolated and inbred population of this critically endangered kākāpō (Dussex et al, submitted); and 5) more complete resolution of the complex sex chromosome organization and evolution in monotreme mammals (Zhou et al, submitted). All of these novel discoveries were not possible with the previous reference assemblies of these or closely related species, and we expect many future discoveries to follow from these new assemblies.

### The Vertebrate Genomes Project (VGP)

Building on this initial set of assembled genomes and the lessons learned, we propose to expand the VGP to deeper taxonomic phases, beginning with Phase 1: representatives of approximately 260 vertebrate taxonomic orders, defined here as lineages separated by 50 or more million years of divergence from each other. Phase 2 will encompass species representing all ~1,000 vertebrate families; Phase 3 will encompass all ~10,000 genera; and Phase 4, nearly all of the 71,657 extant named vertebrate species (https://vertebrategenomesproject.org/). This strategy will take advantage of continuing improvements in genome sequencing technology, assembly and annotation to support scaling to each subsequent phase, while addressing specific scientific questions at increasing levels of phylogenetic refinement. Related large-scale reference genome efforts are adopting lessons learned with the VGP, including the Bat1K^70,46^ (https://bat1k.com), Bird B10K^71,72^ (https://b10k.genomics.cn/), Global Ant Genomics Alliance^73^ (GAGA; http://antgenomics.dk/), Earth Biogenome Project^74^ (EBP. https://www.earthbiogenome.org/), Global Invertebrate Genomics Alliance^75^ (GIGA; http://giga-cos.org/), Darwin Tree of Life (https://www.darwintreeoflife.org/), and human pangenome (https://humanpangenome.org/) projects, which target all species for particular clades or geographic regions of interest, or multiple individuals within a species representing diversity of the extant population. Additional details of the VGP strategy are given in **Box 2**.

#### Box 2. The Vertebrate Genomes Project

The goal of the Vertebrate Genomes Project (VGP) is to generate at least one high-quality, error-free, near gapless, chromosome-level, haplotype phased, and annotated reference genome assembly for all extant vertebrate species and to use those genomes to address fundamental questions in evolution, disease, and biodiversity conservation. We plan to conduct this international project in phases according to phylogenetic scale, from orders (Phase 1) to families (Phase 2), genera (Phase 3), and finally all species (Phase 4; **Fig. B2**). Phase 1 serves as a proof of principle project. At the family level, we would complete vertebrates in the Phase 1 goal of the Earth Biogenome Project (EBP) for high-quality reference genome assemblies for all eukaryotic families^74^. At the genera level, we would complete the original G10K mission of approximately 10,000 vertebrate species^5^. At the final species level, we would complete the data generation mission of the VGP (BioProject ID PRJNA489243) and specific vertebrate taxonomic groups, such as all birds (B10K^71,72^ BioProject PRJNA489244) and all bats (Bat1K^46,70^ BioProject PRJNA489245).

For Phase 1, although there are approximately 150 named orders of vertebrates, the criteria for taxonomic divisions are not consistently applied among vertebrate classes. Therefore we sought to use a more uniform definition. Based on findings from the Avian Phylogenomics Project^20^ and mammalian phylogenomic studies^100^, we noted that who taxonomists have often delimited orders encompass species that shared a most recent common ancestor 50-70 million years ago (MYA), following the last mass extinction event at the Cretaceous-Paleogene transition. Thus, for VGP Phase 1, we aimed to partition lineages that have an inferred common ancestor not substantially older than 50 MYA. This definition resulted in our current target list containing approximately 260 “order level” lineages (http://vgpdb.snu.ac.kr/details/).

When we first began working on the hummingbird assemblies in 2015 and initiated the VGP and sequencing of ordinal representative genomes in 2017, there were 66,178 named species gathered from various databases, estimated based on the IUCN Red List of Threatened Species and reported in the Vertebrate Wikipedia page from 2014 to date (https://en.wikipedia.org/wiki/Vertebrate). This is a number that we had initially used in public announcements of the project^101^. However, we collated available lists of vertebrate species, and we obtained 71,657 named species as of January 2019. We believe the increased number of species is due to additional species discoveries in the last 10 years, revisions of previously defined species (i.e. Northern vs. Southern ostrich), and analyses of genomic relationships^72^. With this list, we have created, for the first time that we are aware of, an all-vertebrate species list (http://vgpdb.snu.ac.kr/splist/). We are populating this list with accessions to the high-quality reference genomes, including the 17 of this study, as well as draft and medium quality genome assemblies. We hope that this list will be useful to the scientific community to track genome assemblies for all vertebrate species.

To conduct the VGP in an efficient and democratic manner, we built a governance and committee structure that consists of an executive council and task-specific committees focused on specific issues, including permits, sample preparation, genome assembly, genome annotation, comparative genomics, and conservation genomics. We developed an assembly pipeline scaleable on the cloud environment, where working data is hosted on an Amazon s3 bucket (s3://genomeark). The production of the VGP assemblies is performed on DNAnexus, which is available for VGP members. The entire source code of the pipeline to run locally or on the DNAnexus platform is publicly available on github (https://github.com/VGP/vgp-assembly). The scaffolding pipeline is also available to run on generic architecture and Docker containers (**Supplementary Note 4**). Intermediate assemblies and raw data are available to download on Genome Ark (https://vgp.github.io) until archived. We also built a public website for the VGP (https://vertebrategenomesproject.org/), and its parent G10K website, with links to associated projects (B10K, Bat1K, and EBP). Our assemblies and raw sequences are deposited in international public databases with NCBI and EBI under a VGP BioProject ID PRJNA489243 (https://www.ncbi.nlm.nih.gov/bioproject/489243). We currently produce about three genome assemblies per week, but will need to scale up to 125 genomes per week to complete all ~70,000 species within a 10-year period, assuming the availability of future funding and the development of more advanced computational infrastructure. In addition to the 17 genomes released publicly with this publication, there are 100 more assemblies in progress (https://docs.google.com/spreadsheets/d/1s5J-s3Tat3U_wQcik_xhVHwH6AAXr5D9AMRu-e22XDw/edit?usp=sharing), that are supported by individual institutions and crowd-funding amongst scientists (https://genome10k.soe.ucsc.edu/data-use-policies/).

**Fig. B2.**
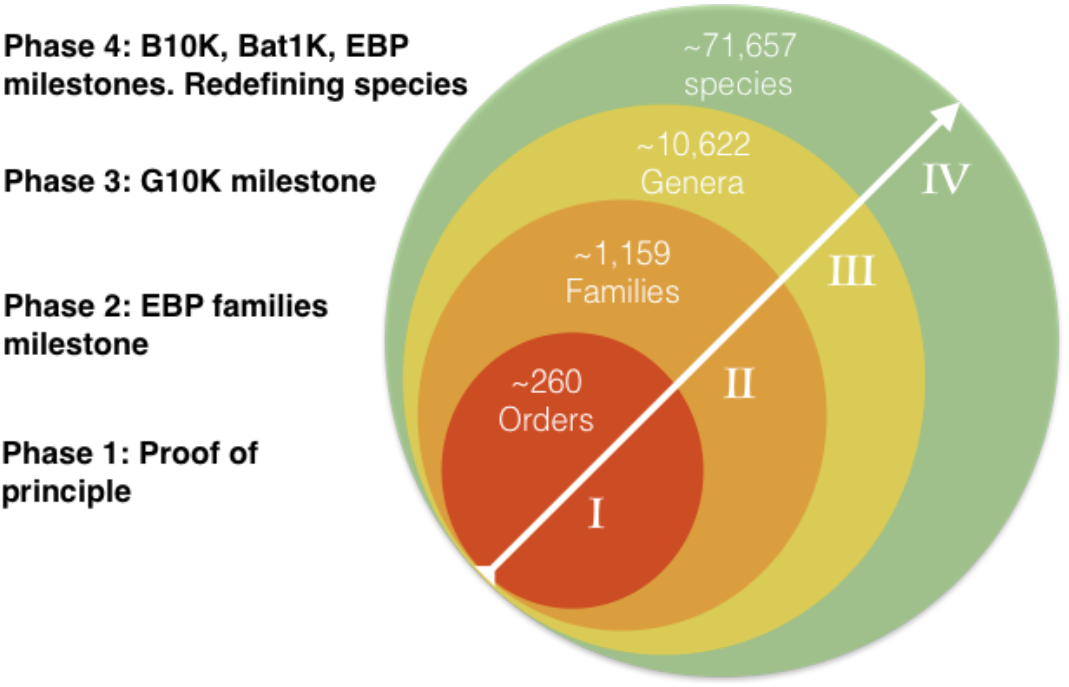
Schematic of proposed phases to conduct the VGP. Circles represent phylogenetic classification scales, going from smaller to larger numbers of species (arrow). A sequenced species represents an order, family, and genera in Phases 1-3. To the left are listed goals and related projects for whose milestones will be completed at the completion of specific VGP phases. Redefining species means that within Phase 4 it might become possible to use the genome sequence differences to determine when individuals should be considered belonging to the same species or their own distinct species^72^.

### Proposed assembly quality metrics

As described above, during this study we developed an initial quality goal in terms of continuity (contig N50 > 1 Mbp, scaffold N50 > 10 Mbp), completeness (>90% assigned to chromosomes), and base accuracy (>99.99%). We propose summarizing these minimum standards using the notation “6.7.Q40”, with the first digit representing the log-scaled contig NG50 size, the second digit the log-scaled scaffold NG50 size, and the QV as Phred-scaled base accuracy. We also propose using a “C” character to denote “complete” contigs or scaffolds that reach telomere-to-telomere continuity. This is revised from a prior notation we developed and noted elsewhere^46,70,74^, which reported log-scaled continuity measured in kilobases rather than bases. However, based on lessons learned here and knowing that these three metrics do not capture all important variables, we now propose to characterize genome assembly quality more comprehensively using 14 metrics under six categories: continuity, structural accuracy, base accuracy, haplotype phasing, functional completeness, and chromosomal status (**Table 1**).

**Table 1.** Proposed standards and metrics for defining genome assembly quality. The six broad quality categories in the 1st column are split into sub-metrics in the 2nd column. The recommendations for draft to finished qualities (columns 3–6) are based on those achieved in past studies, this study, and what we aspire to. X.Y.Q = contig NG50 log10 value; scaffold NG50 log10 value; and QV value. The range of metrics in the VGP assemblies of this study (last column) satisfy a combination of the 6.7.Q40 and 7.C.Q50 metrics; specific values are in Fig. 3a and Supplemental Tables 11-12,17. Transcript mappability was only determined for those species for which there was transcriptome data. *Phase blocks were calculated only for the zebra finch non-trio assembly using haplotype specific k-mers from parental data^48^; the trio assemblies of the same species had NG50 phase blocks of 17.3 Mb (maternal) and 56.6 Mb (paternal).

| Quality Category | Quality Metric | Finished | 7.C.Q50 | 6.7.Q40 | 4.5.Q30 | VGP |
| --- | --- | --- | --- | --- | --- | --- |
| Continuity | Contig (NG50) | = Chr. NG50 | >10 Mbp | >1 Mbp | >10 kbp | 1-25 Mbp |
|  | Scaffolds (NG50) | = Chr. NG50 | = Chr. NG50 | >10 Mbp | >100 kbp | 23-480 Mbp |
|  | Gaps / Gbp | No gaps | <200 | <1,000 | <10,000 | 75-1500 |
| Structural accuracy | False duplications | 0% | <1% | <5% | <10% | 0.2-5.0% |
|  | Reliable blocks | = Chr. NG50 | >90% of Scaffold NG50 | >75% of Scaffold NG50 | >50% of Scaffold NG50 | 2-75% |
|  | Curation improvements | All conflicts resolved | Automated + Manual | Automated | No requirement | Automated + Manual |
| Base accuracy | Base pair QV | >60 | >50 | >40 | >30 | 39-43 |
|  | k-mer completeness | 100% complete | >95% | >90% | >80% | 87-98% |
| Haplotype phasing | Phased block (NG50) | = Chr. NG50 | >1 Mbp | >100 kbp | No requirement | 1.6 Mbp* |
| Functional completeness | Genes | >98% complete | >95% complete | >90% | >80% | 82-98% |
|  | Transcript mappability | 98% | >90% | >80% | >70% | 96% |
| Chromosome status | Assigned % | 98% | >90% | >80% | No requirement | 94.4-99.9% |
|  | Sex chromosomes | Right order, no gaps | Localized homo pairs | At least 1 shared (e.g. X or Z) | Fragmented | At least 1 shared |
|  | Organelles (e.g. MT) | 1 Complete allele | 1 Complete allele | Fragmented | No requirement | 1 Complete allele |

These measures have been described throughout this study, and we provide additional details in **Box 3**. We further note here, that base error rate inferred directly from *k-mers* was more comprehensive and accurate than the widely used mapping and variant calling protocols, which artificially inflated QV values (**Supplementary Table 17**) because they exclude regions that are difficult to map. VGP assemblies exceeding Q40 contained fewer frameshift errors, as predicted^76^, and therefore we recommend targeting a minimum QV of 40 whenever possible. Haplotype phasing and false duplications are the most underdeveloped measures, presumably because this is an under-appreciated area of need, but recent tools, developed here and elsewhere^27,48,51,53,77^, are helping to address this.

#### Box 3. Assessing overall genome assembly quality

Below are examples of measures for the six categories of genome assembly quality proposed in this study (**Table 1**).

##### Continuity

The current most popular measure of genome assembly continuity is the scaffold N50, and secondarily the contig N50, defined as the largest *s* where scaffolds (or contigs) of length *s* or greater is half or greater the total assembly size. However, the assembly size can be larger or smaller than the true genome size depending on the assembly tools and data quality used. Thus, we recommend using NG50 (G for genome), which uses the estimated genome size instead of the assembly size for normalization^102^. We prefer to estimate the genome size from actual sequence data, using *k-mers* (sequence fragment length of *k*) such as done in GenomeScope^47^. Note that all high copy *k-mers* should be included when counting *k-mers* to properly include repeat contents, which is not a default behavior in most k-mer counters. For gaps, we recommend using a measure of the rate of gaps per unit of Gbp assembled, as larger genomes would be unfairly penalized for more gaps.

##### Structural accuracy

To assess reliable blocks of structural accuracy without a known truth, we propose mapping the raw data types to the final assembly and measure concordance. We define concordance here as how many of the data types support the assembled structure at each base. In this study, we defined reliable blocks with support from at least two of the four sequencing platforms (CLR, linked reads, Opt, and Hi-C). This can be extended with additional data types, such as genetic maps or FISH karyotypes, when available.

##### Base accuracy

###### Base pair QV

There are multiple ways of measuring base-level accuracy (QV). One approach is to align (i.e map) highly accurate reads to the assembled genome and call base errors similarly to variant calling. We define “mappable” as all reads that align, excluding low-coverage and excessively high-coverage regions (see **Supplementary Methods** for exact parameters used), where we can rely on base error calls. The lower mappability to the alternate contigs in our nontrio assembly reflects the biased quality that affected mapping for polishing, suggesting that the primary assembly set should be used for most downstream analyses. The other, more reliable, way to measure base accuracy is using *k-mers* found both in the assembly and highly accurate unassembled reads. All *k-mers* found only in an assembly are likely produced from a base pair error. By counting these *k-mers* and comparing the fraction to all *k-mers* found in an assembly, we can estimate the error rate and calculate the quality value using the *k-mer* survival rate^48^. We found *k-mer* based methods include unmappable regions and thus avoid over-estimated QVs from the mapping-based approach.

###### *K-mer* completeness

To assess if all bases in a genome are properly assembled, we propose using *k-mers* as the truth set to get an estimate. Reliable *k-mers* obtained from highly accurate reads are obtained by excluding erroneous *k-mers* from sequencing errors. The fraction of the *k-mers* found in the assembly of these reliable *k-mers* are indicative for genome completeness. This measure is dependent on base level accuracy as well, because *k-mers* from assembly errors will affect the completeness measure. We use the implementation in Merqury^48^ to obtain the fraction of reliable *k-mers*, reported in **Fig. 3a**.

###### False duplications

For a *k*-mer size that is sufficiently long to be unique in the genome, and a genome sequenced with high-fidelity reads to a depth of coverage *c*, a complete *de novo* assembly should recover *k*-mers from the homozygous (two-copy) regions of the genome with roughly *c* times and *k*-mers from the heterozygous (single-copy) regions with *c/2* times. All *k-mers* in the heterozygous and homozygous regions are expected to be found once in a (pseudo) haplotype assembly. Any additional *k-mer* copy found in the assembly compared with the high-fidelity reads are considered to be falsely duplicated^77^. We use the implementation in Merqury^48^ to count the number of distinct *k-mers* with additional copies in the heterozygous and homozygous regions and report the relative portion to the expected *k-mers* with no additional copies.

###### Haplotype phasing

We propose to use phase block NG50s as a measure for haplotype consistency. A phase block is expected to match one of the parental haplotype sequences, with no haplotype switches. Haplotype consistency is important for gene annotation, because haplotype switches could mix the true gene structure, creating an artificially mixed gene that does not exist in nature. Currently, the most reliable way to measure phase consistency is by using parental sequences. In this study, we use Merqury^48^ to infer haplotype blocks from haplotype specific *k-mers*. Accounting for sequencing errors accidentally corrupting a true haplotype specific *k-mer*, we allow short-range switches to occur up to 100 times within 20kbp, which is the typical length one CLR read can phase. We expect block sizes to be more dependent on genome heterozygosity levels, where less heterozygous genomes will have longer runs of homozygosity (ROH) that prevent linking of heterozygous sites when no parental information is used. Heterozygosity will also vary across segments of a genome, and thus, one value may not be equally applicable across the genomes. Therefore, we set smaller block NG50 requirements in the quality metric (**Table 1**), independent of chromosome sizes except for the “finished” quality. Measures of the fraction of the genome that is haplotype-phased are not as well developed with no parental data, presumably because the importance of phasing to prevent errors has been unappreciated. This measure pertains to not only diploid genomes, but also polyploid genomes, which is found in amphibians and fishes.

###### Functional completeness

Gene-based metrics could be used as an indicator for genome completeness and is one of the most important factors when conducting functional studies. However, it is almost impossible to have a truth set of all genes, especially for genomes with no reference available. One indirect way to measure functional completeness is by using BUSCO genes sets, which are sets of highly conserved orthologous genes present in single copy across vertebrates or other groups of species^49,50^. To work properly, the sequences of the gene set needs to be complete and error free, but this is not the case for many BUSCO genes^22^. Without it being complete and with natural gene losses in some species, there will be an upper limit of less than 100% mapping. Overall however, the absence or apparent duplication of these genes in an assembly may be evidence of assembly error. Transcript mappability is another indirect way to measure gene completeness, because a more complete genome is expected to map more transcriptome sequences unambiguously (uniquely) to the assembly.

###### Chromosome status

For defining scaffolds as chromosomes, we believe the current best tool besides genetic linkage or FISH karyotype mapping is Hi-C mapping. We consider a scaffold as a complete chromosome (albeit with gaps) when there is a clear diagonal in the Hi-C mapping plot for that scaffold and there are no other large scaffolds that can be placed in that same scaffold. The Hi-C maps prove useful for identifying large-scale structural aberrations in the assemblies, including false chromosome fusions. The more uniform the Hi-C signal across the main diagonal, the more likely the assembly structure is correct. High-frequency, off-diagonal Hi-C interactions are a strong sign of mis-assembly, which can be corrected with manual curation. Based on these criteria, one can then estimate the percent of the genome that is assigned to chromosomes. See Lewin et al.^103^ for an alternate view of naming scaffolds.

###### Sex chromosomes

Sex chromosomes are typically a challenge as they are often highly diverged between the partners. The sex-specific chromosome (e.g. Y in XY mammals or W in ZW birds and snakes) are often rich in highly repetitive heterochromatin. Sex determination mechanisms are highly variable in amphibians, reptiles, and fishes, with different sex genes (mostly unknown) defining non-homologous sex pairs. In many species, it is unclear whether there is male heterogamety (XY as in mammal) or female heterogamety (ZW as in birds). Many reptiles and some fish have no sex chromosomes and determine sex by an environmental signal (commonly temperature). Thus, we only require sex chromosomes to be assembled and identified in lineages known to have sex chromosomes, and make an effort to sequence the heterogametic sex to assemble both sex chromosomes, or one of each sex to have greater confidence. Once a pseudohaplotype assembly is assembled, sex-specific chromosomes can be further determined by comparing read depth in males and females when available, identification of known of sex-specific genes for the relevant clade, synteny with sex chromosomes in closely related species, and the coverage pattern of PAR and haploid regions.

These metrics serve as a guideline for presenting the state of an assembly and to better understand the applications to which it is suited. We are aware that it is difficult to define a single standard across all metrics, because different methods can yield high quality on different metrics and the required quality of an assembly depends on the biological questions it is meant to answer. Thus, we propose measuring each facet of assembly quality separately. For example, the 10XG linked read assembly of the Anna’s hummingbird would be categorized as “4.5.Q40”, with low continuity and high base accuracy. Such a genome would be suitable for use as a reference for population-scale SNP surveys. If instead, a genome is to be used for studies of chromosomal rearrangements and gene synteny, then “6.7.Q40”, with high structural and base accuracies and > 90% assigned to chromosomes, would be necessary, as was achieved for the VGP reference kākāpō assembly presented here. For gene-based studies, a pseudo-haplotype reference with local phasing to preserve accurate gene structures is sufficient and high-quality phasing across chromosomes may be unnecessary.

“Finished” quality is obviously the ideal assembly result, but this level of quality is currently routine for only bacterial and some non-vertebrate model organisms with smaller genome sizes that lack large centromeric satellite arrays^78–80^. The possibility of achieving complete, telomere-to-telomere assemblies of vertebrate and other eukaryotic genomes is foreseeable, given our identification of some assembled avian and bat chromosomes with zero gaps in this study and the recent ultra-long ONT-based assembly of the first complete human X chromosome^81^. Although the latter project utilized a fully homozygous cell line (CHM13) and extensive manual curation, we are optimistic that, given continuing advances in diploid sequencing and assembling technology, finished-quality reference genomes will be achievable at reasonable cost for most species of interest within the next decade.

### Future efforts

Going forward, the VGP has identified areas that still need improvements, including more accurate and complete haplotype phasing, improved base-call accuracy, better resolution of long repetitive regions (e.g. telomeres and centromeres), more complete sex chromosomes, and fewer genes or organelle genomes left unassembled (**Supplementary Note 3**). Improved scalability and throughput is also needed throughout the process, from sample permitting, collection, and preparation, to sequencing, assembly, annotation, data transfer, and presentation. Each of these are crucial to achieving high-quality reference genome assemblies for tens of thousands of vertebrate species and millions of eukaryotic species. Ultimately, we aim to enable the complete and accurate assembly of any vertebrate genome by anyone with sufficient resources to collect a fresh sample and pay the sequencing costs. The VGP is working towards these goals and making all data, protocols, and pipelines openly available. We encourage the scientific community to use and evaluate the assemblies, with the associated raw data, and to provide feedback towards improving all processes for complete and error-free assembled genomes of all species.

Despite remaining imperfections, our reference genomes are the highest quality to date for each of the species sequenced. When we began generating genomes beyond the Anna’s hummingbird for the VGP in 2017, only eight vertebrate species were represented in GenBank with genomes that met our target continuity quality metrics (Supplementary Table 22). Among these were the human and mouse genomes with initial assemblies that required billions of dollars to generate, mostly using the Sanger sequencing approach. Of the remaining six (consisting only of mammals and fish), four were generated by members of the VGP using lessons learned by the group, and none were haplotype phased. We now provide another 16 species, all with some haplotype separation. Many others generated following the VGP model are in progress as part of the Phase 1 VGP (100 species as of April 2020; BioProject PRJNA489243). We hope our efforts will continue to serve as a model not only for vertebrate genomes, but for all species, and we look forward to learning more from the scientific community on how to achieve these objectives.

## Submitted papers as part of VGP package

Dussex N, Walk T, Wheat C, Díez del Molino D, von Seth J, Foster Y, Kutschera V, Guschanski K, Rhie A, Phillippy A, Korlach J, Howe K, Chow W, Pelan S, Damas J, Lewin H, Hastie A, Fedrigo O, Guhlin J, Harrop T, Le Lec MF, Dearden P, Haggerty L, Martin F, Kodali V, Thibaud-Nissen F, Iorns D, Knapp M, Gemmell N, Robertson F, Moorhouse R, Digby A, Eason D, Vercoe D, Howard J, Jarvis ED, Robertson B, Dalen L. Purging of genetic load in the highly inbred and critically endangered kākāpō. *Submitted*.

Zhou Y, Shearwin L, Li J, Song Z, Hayakawa T, Stevens D, Fenelon J, Peel E, Cheng Y, Pajpach F, Bradley N, Suzuki H, Nikaido M, Damas J, Daish T, Perry T, Zhu Z, Geng Y, Rhie A, Sims Y, Wood J, Fedrigo O, Li Q, Yang H, Wang J, Johnston S, Phillippy AM, Howe K, Jarvis ED, Ryder O, Kaessmann H, Donnelly P, Korlach J, Lewin H, Graves J, Belov K, Renfree MB, Grutzner F, Zhou Q, Zhang G. Genomes of the egg-laying monotremes reveal the biology and evolution of mammals. *Under revision*.

Theofanopoulou C, Gedman G, Cahill JS, Boeckx C, and Jarvis ED. Universal nomenclature for oxytocin-vasotocin ligand and receptor families. *In press*. (2020)

**<u>Extended Data Figures and Table</u>**

**<u>Extended Data References</u>**

**<u>Methods</u>**

**<u>Methods References</u>**

**<u>Data availability</u>**

**<u>Code availability</u>**

**<u>Acknowledgements</u>**

**<u>Author contributions</u>**

**<u>Competing interests</u>**

**<u>Author affiliations</u>**

**<u>Additional Information (Box 1–3)</u>**

<u>**Supplementary Tables:** Separate Worksheet Document</u>

<u>**Supplementary Information (Notes and Methods):** Separate Document</u>

## Extended Data Figures and Table

**Extended Data Figure 1.**
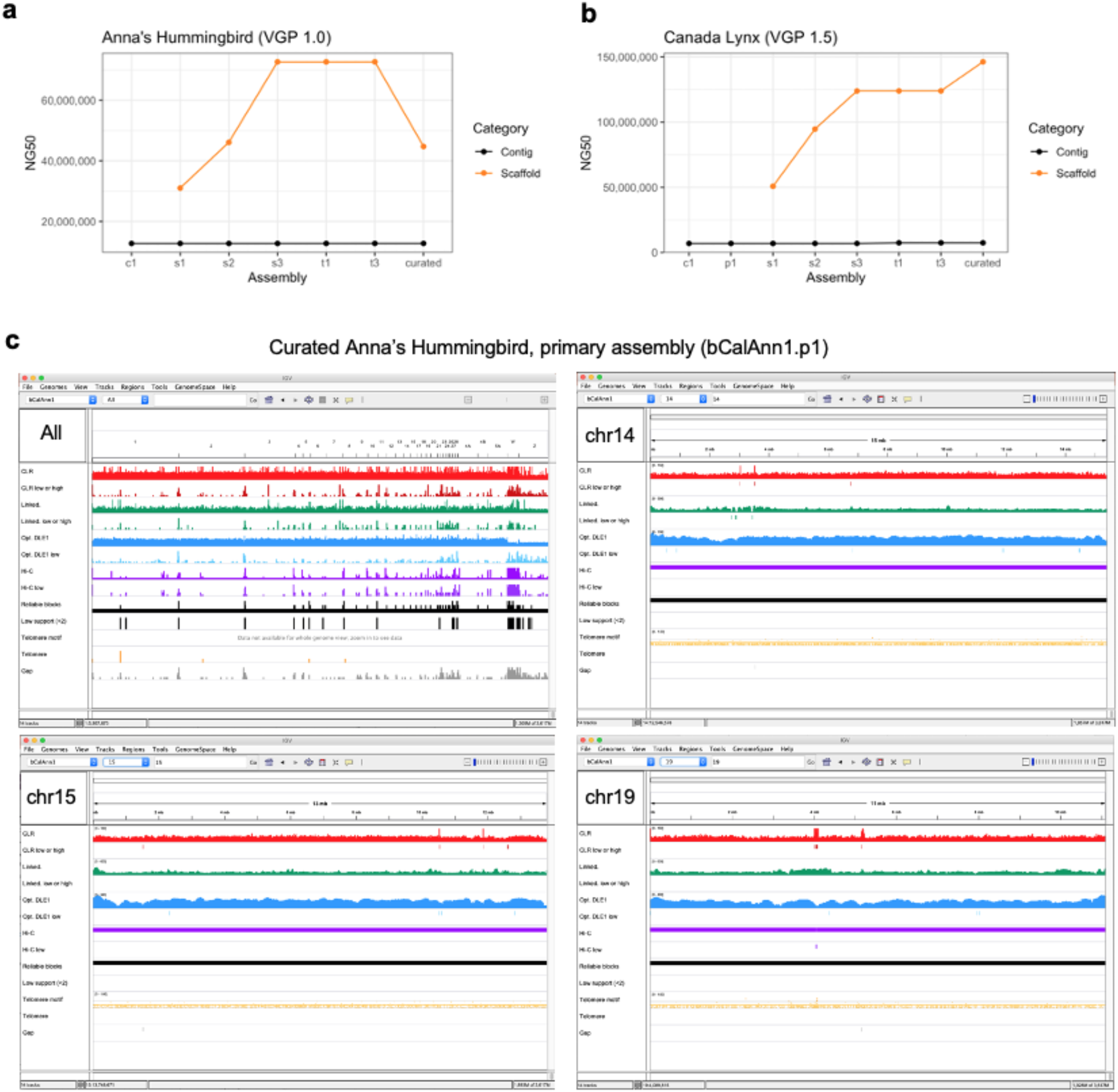
Assessment of completeness of the Anna’s hummingbird assembly. **a-b,** Steps and NG50 continuity values of the VGP assembly pipeline that gave the highest quality assembly for Anna’s hummingbird and Canada lynx in this study. The specific steps are outlined further in Extended Data Fig. 2a, and Methods. **c**, Whole genome alignment of CLR (red), linked reads (green), optical maps (blue), and Hi-C reads (purple) of the Anna’s hummingbird, along with telomere motif (TTAGGG and its reverse complement, yellow) and gaps (gray). For each data type, the first band shows the mapped coverage, while the second shows the number of counts of low coverage or signs of collapsed repeats. Larger chromosomal scaffolds (1-19) have fewer gaps and low coverage or collapsed regions compared with the micro chromosomes (20~33). Chromosome 14, 15, 19 of the Anna’s hummingbird were the most structurally reliable scaffolds, having only one gap each with no low supportive regions.

**Extended Data Fig. 2.**
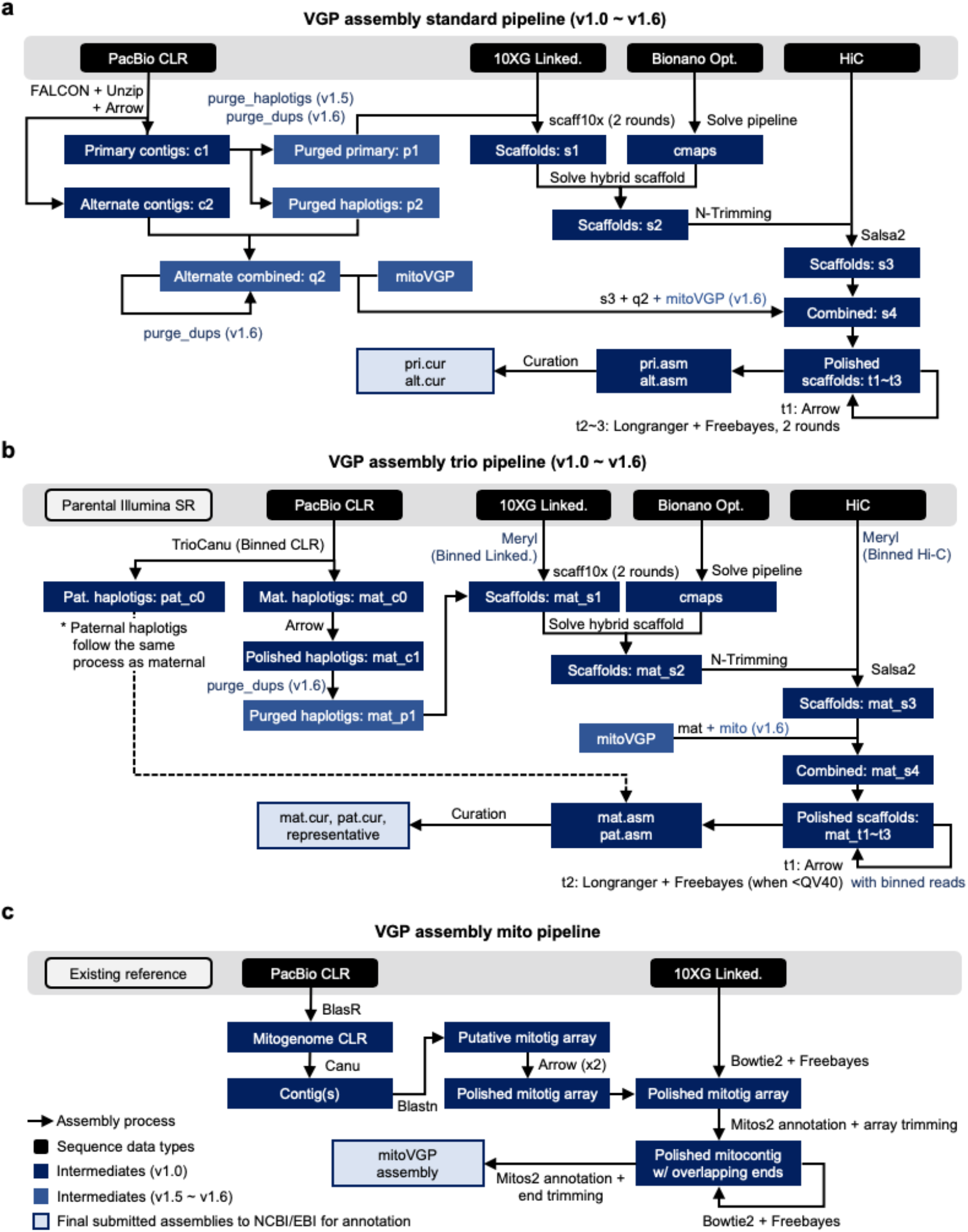
Flow charts of assembly pipelines used to generate high quality assemblies in this study. **a,** Standard VGP assembly pipeline when sequencing data of one individual. **b**, Standard VGP trio assembly pipeline when DNA is available of a child and parents. Dashed line indicates that the other haplotype went through the same steps prior to curation. In addition to the curated assemblies per haplotypes, a representative haplotype with both sex chromosomes is submitted. **c**, Mitochondrial assembly pipeline. Figure legend applies to **a-c**. Steps newly introduced in v1.5 ~ v1.6 are highlighted in light blue. Abbreviations: c - contigs; p - purged false duplications from primary contigs; q - purged alternate contigs; s - scaffolds; t - polished scaffolds. Further details and instructions are available at https://github.com/VGP/vgp-assembly.

**Extended Data Fig. 3.**
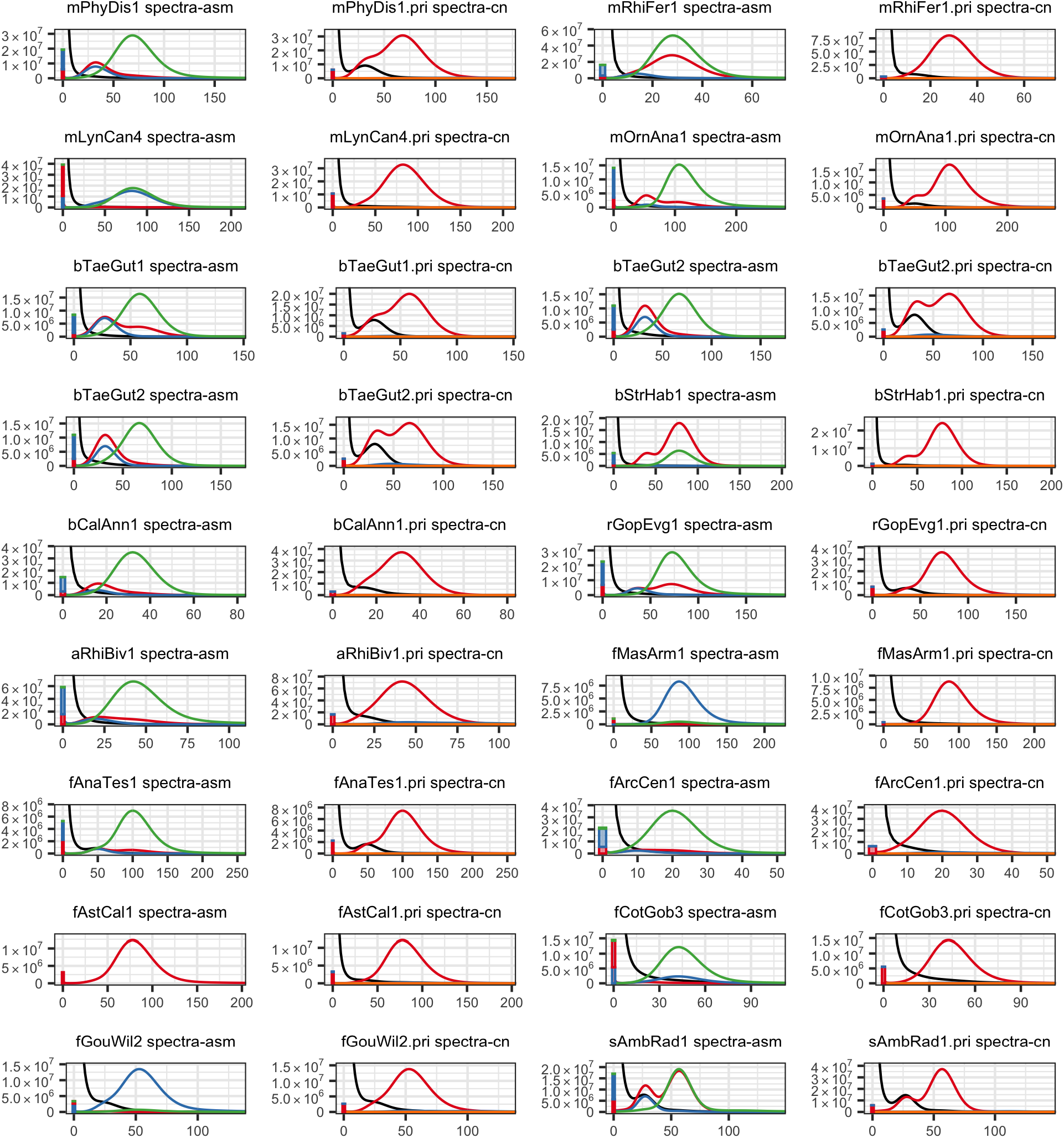
*k-mer* spectrum of each submitted assembly, plotted on the 21-mer multiplicity found in the illumina sequencing set from the linked reads. For each species, the left two columns of graphs show the overall *k-mers* found (spectra-asm) only in the primary set (red), alternate set (blue), shared in both assemblies (green), and missing *k-mers* in any assembly set (black). Fewer missing *k-mers* observed (black) indicates the assembly more completely represents the genome. The right two columns show the copy number spectrum (spectra-cn) of the primary assembly set, colored by the copy numbers found in the assembly: once (red), twice (blue), 3 (green), 4 (purple), >4 (orange), and missing (black). *k-mers* in spectra-cn are expected to be found once (red) in a pseudo haplotype assembly; thus, *k-mers* found more than once (blue, green, purple, and orange) originate from false duplicated sequences (homotype duplication) assuming no allele-specific duplications exist in the genome.

**Extended Data Fig. 4.**
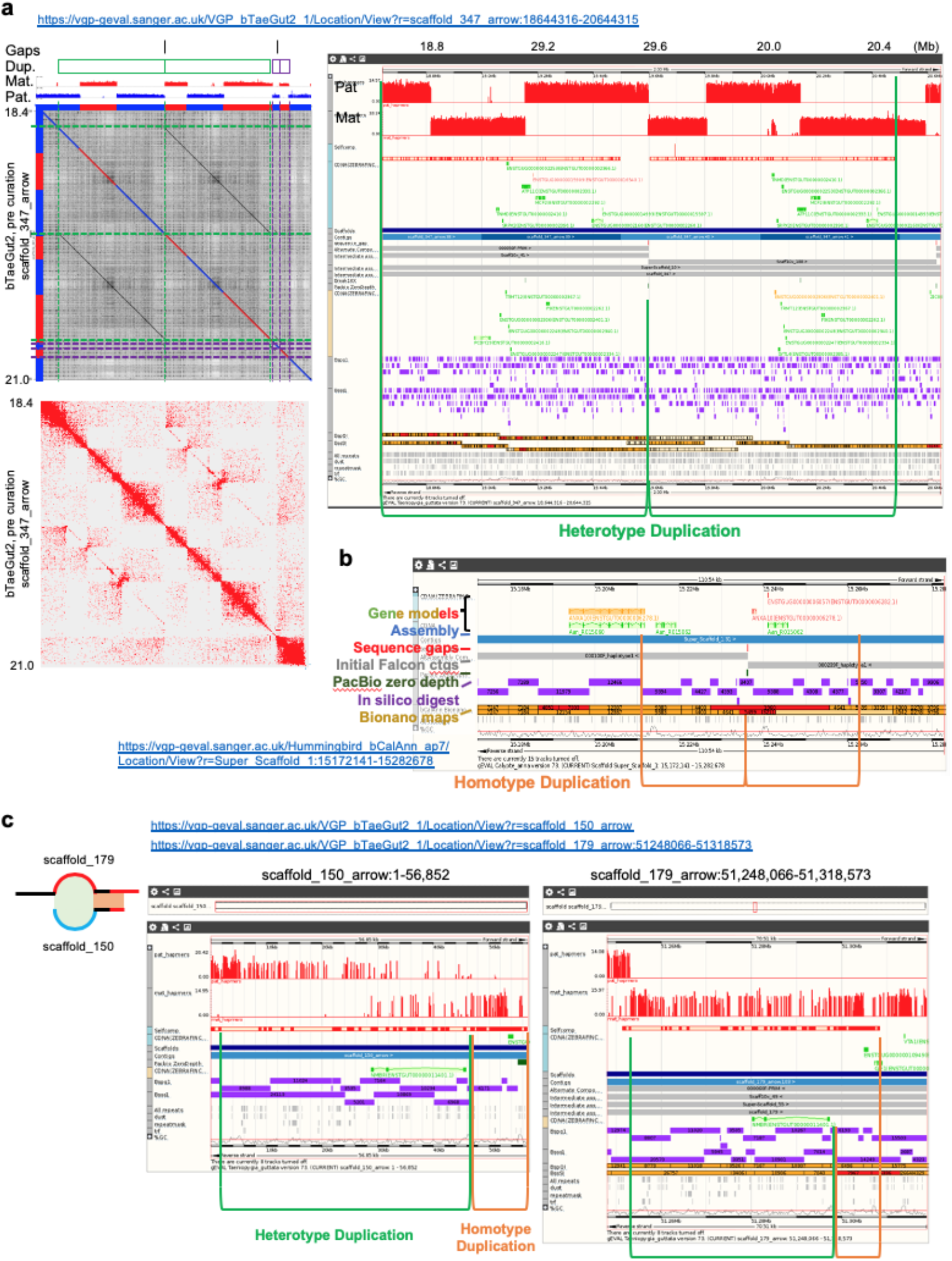
False duplication examples fixed during manual curation. **a**, An example of a heterotype duplication in the female zebra finch, non-trio assembly. Left, a self-dot plot of this region generated with Gepard^1^, sequences colored by haplotypes. Gaps, duplicated sequences (green and purple), and haplotype specific marker densities are indicated on the top. Right, a detailed alignment view of the green haplotype duplication with paternal, maternal markers, self-alignment components, transcripts annotated, contigs, bionano maps, and repeat components displayed on gEVAL^2^. **b**, Example of a homotype duplication found in the hummingbird assembly. These were caused by an algorithm failure in FALCON, which was later fixed. **c**, Example of a combined duplication involving both heterotype (green) and homotype (orange) duplications. Assembly graph structure is shown on the left for clarity, highlighting the overlapping sites at the contig boundary shaded following the duplication type, respectively. Assembly errors including above false duplications were detected and fixed during the curation process.

**Extended Data Fig. 5.**
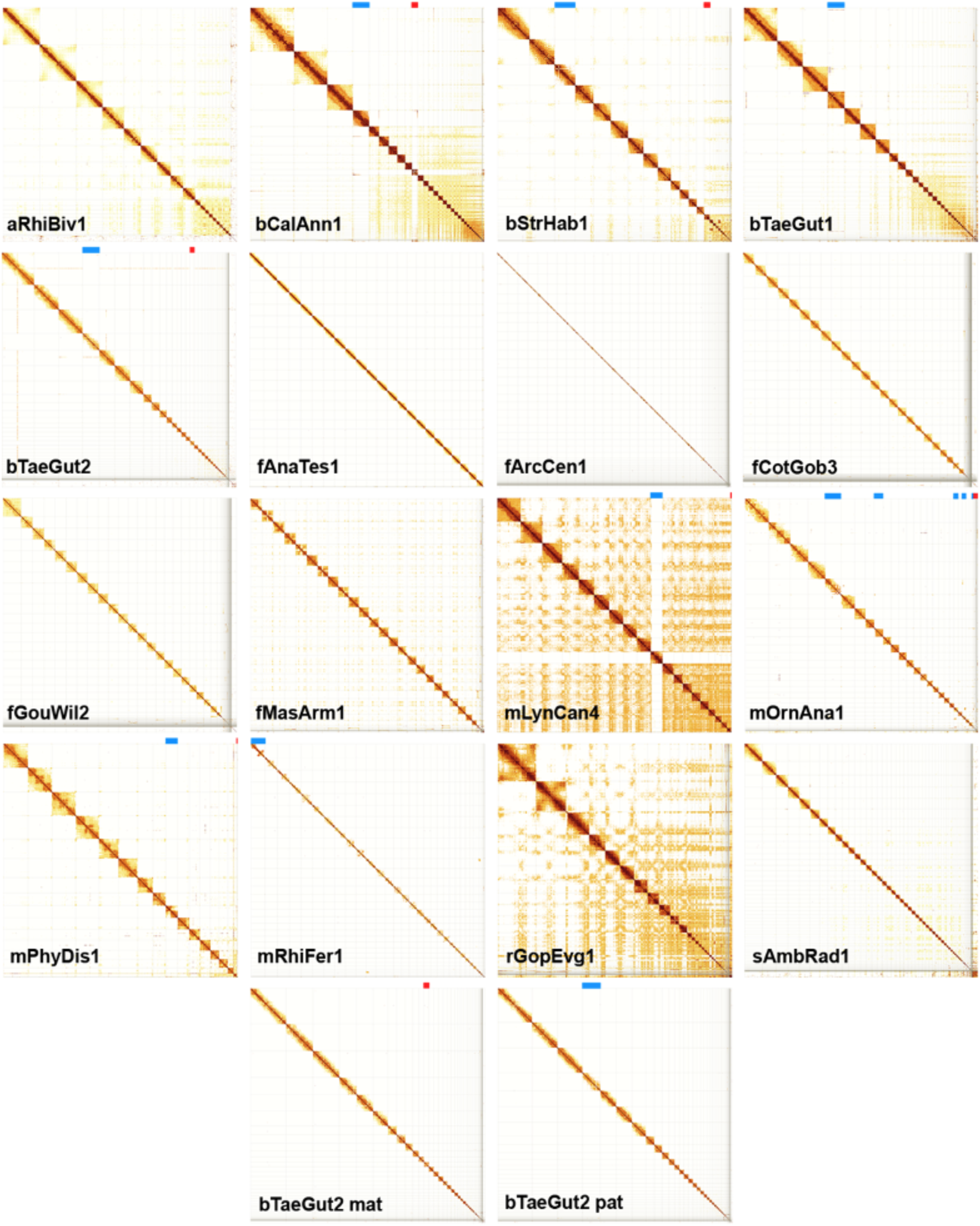
Evidence of near complete chromosome scaffolds in the VGP assemblies. Shown are Hi-C interaction heatmaps for each species after curation, visualized with PretextView^3^. A scaffold is considered a putative arm-to-arm chromosome when all Hi-C read pairs in a row and column map to a square (i.e. assembled chromosome) on the diagonal without any other interactions off the diagonal. Those with remaining off-diagonal matches to smaller scaffolds are not linked because of ambiguous order or orientation, are instead submitted as “unlocalised” belonging to the relevant chromosome. Bands at the top of each heatmap shows scaffolds identified as XZ (blue) or YW (red) sex chromosomes. The Hi-C map of fAstCal1 is not included as we had no remaining tissue left of the animal used to generate Hi-C reads.

**Extended Data Fig. 6.**
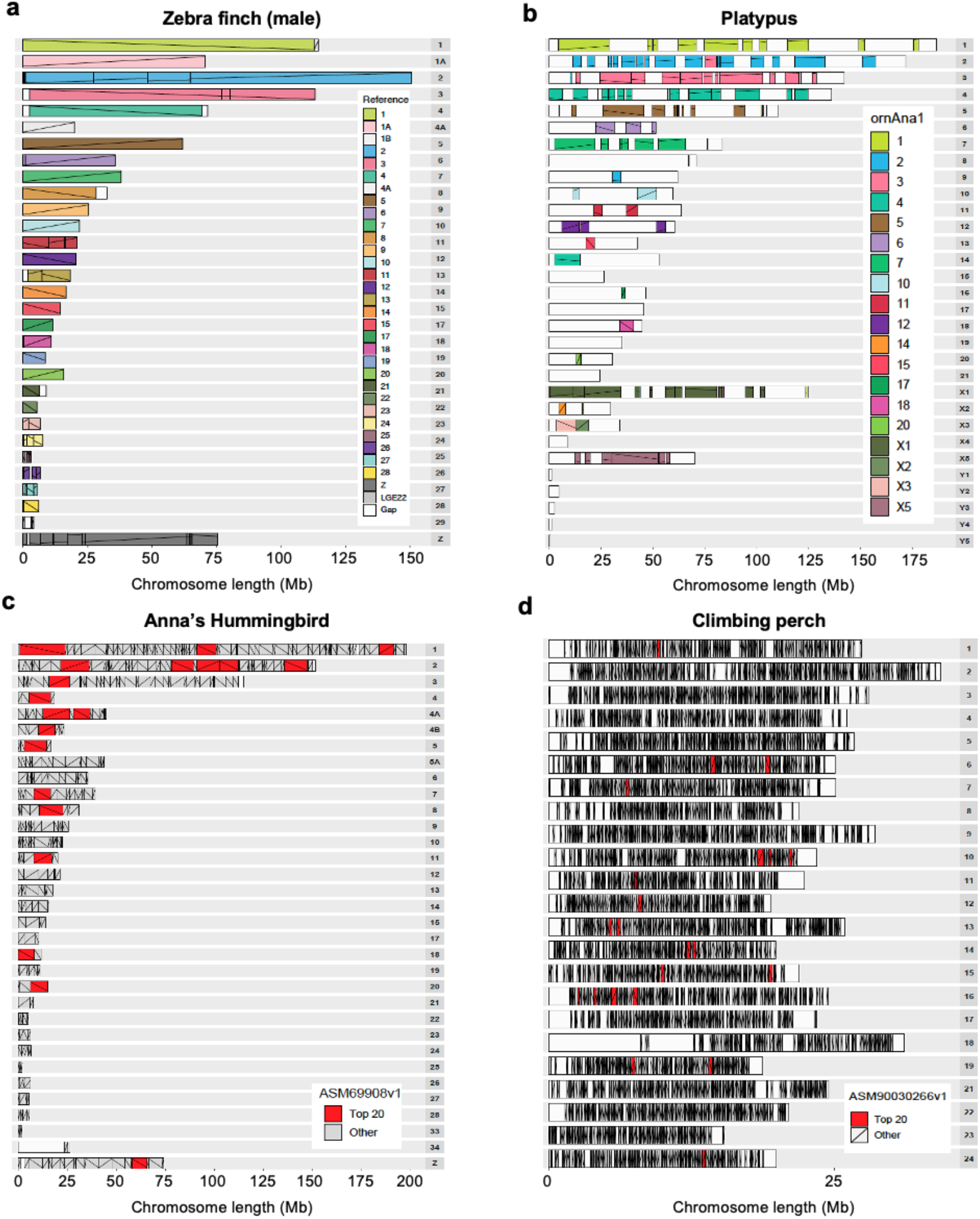
Comparison of chromosomal organization between previous and new VGP assemblies. **a**, Zebra finch male to a reference of the same animal. **b**, Platypus male compared with the previous assembly of a reference female, and thus the Y chromosomes are not compared. **c**, Hummingbird female to a reference of the same animal. **d**, Climbing perch to a previous reference. Each row represents a VGP generated chromosome for the target species. Colors depict identity with the reference (legend on the right); more than one color indicates reorganization in the VGP assembly relative to the reference. The lines within each block depict orientation relative to the reference; a positive slope is the same orientation as the reference, whereas a negative slope is the inverse orientation. Gaps are white boxes with no lines, in the reference relative to the VGP assembly. The accession numbers of the assemblies compared are listed in **Supplementary Table 19**.

**Extended Data Fig. 7.**
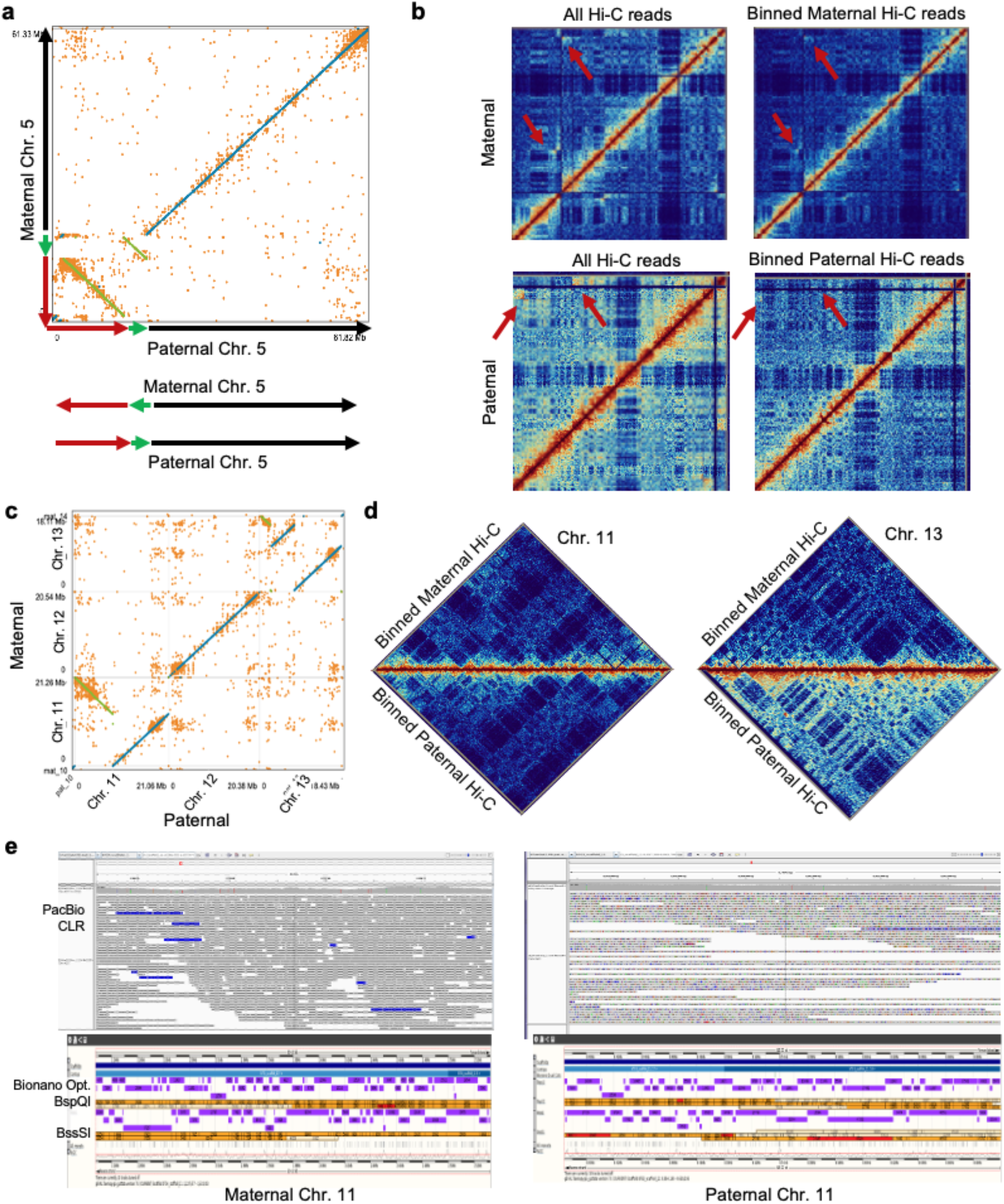
Large haplotype inversions with direct evidence in the zebra finch trio assembly. **a,** Two inversions (green and red) in chromosome 5 found from NUCmer^4^ alignment of the maternal and paternal haplotype assemblies, visualized with dot^5^. **b**, Hi-C interaction plot showing that the trio-binned Hi-C data removes most of the interactions from the other haplotype (red arrows), which could be erroneously classified as a mis-assembly if one haplotype was used as a reference. **c**, A 8.5 Mb inversion found on chromosome 11 and a complicated 8.1 Mb rearrangement on chromosome 13 between maternal and paternal haplotypes. **d**, No mis-assembly signals were detected from the binned Hi-C interactions plots, indicating the haplotype specific inversions are real. **e**, Half the PacBio CLR span and Bionano optical maps agrees with the inversion breakpoints in chromosome 11, supporting the haplotype specific inversion.

**Extended Data Fig. 8.**
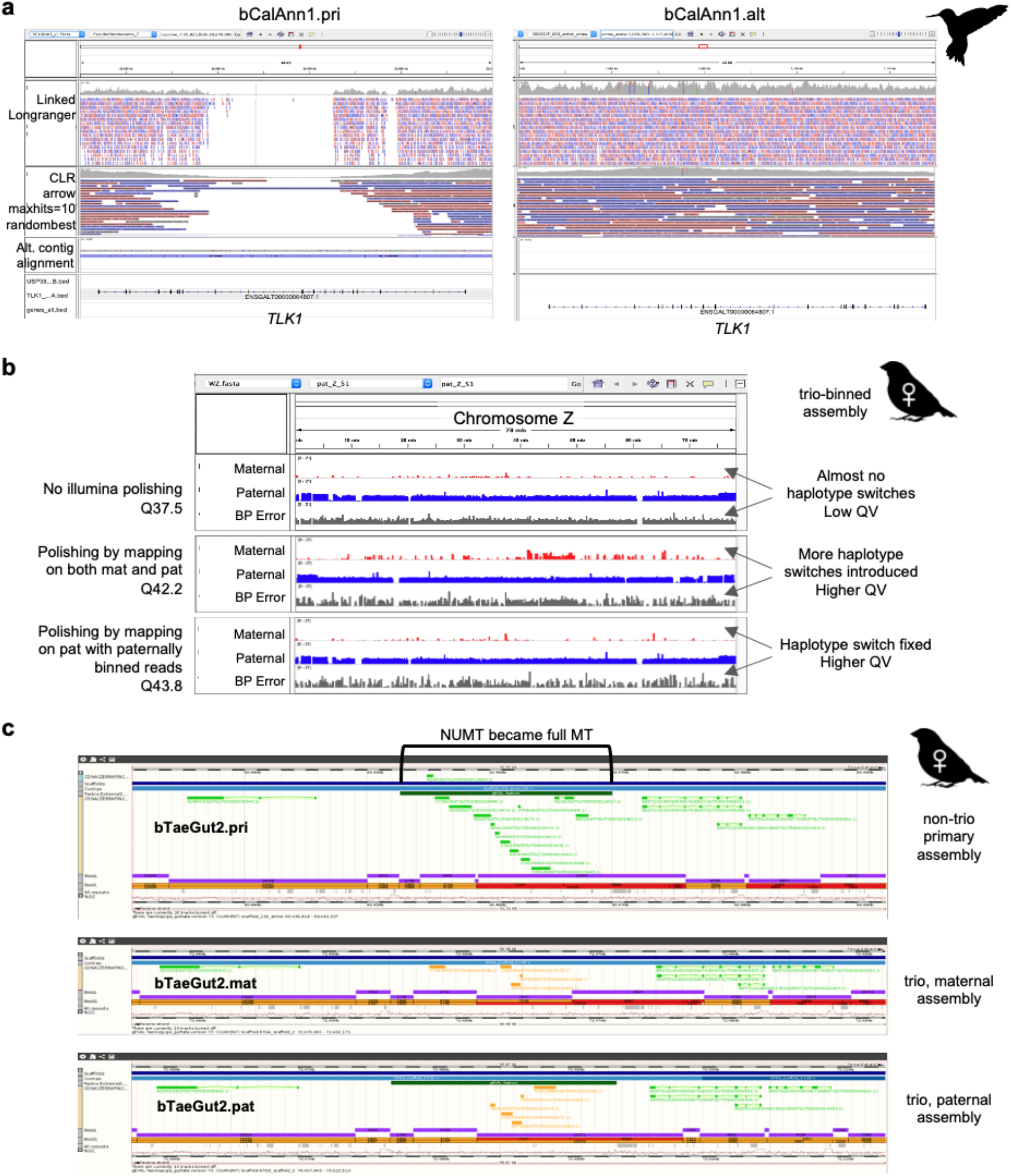
Polishing artifacts. **a,** An example of uneven mapping coverage in the primary and alternate sequence pair of the Anna’s hummingbird assembly. In this example, the alternate (alt) sequence was built in a higher quality, attracting all linked-reads for polishing. The matching locus in the primary (pri) assembly was left unpolished, resulting in frameshift errors in *TLK1* gene. **b**, Haplotype specific markers (red for maternal, blue for paternal) and error markers found in the assembly on the Z chromosome (inherited from the paternal side) of the trio-binned female zebra finch assembly. Each row shows markers before short-read polishing, short-read polishing by mapping all reads to both haplotype assemblies, and polishing by mapping paternally binned reads to the paternal assembly. Polishing improves QV, but introduces haplotype switches when using reads from both haplotypes as shown in row 2. This can be corrected and avoided when using haplotype binned reads, even in short-read polishing. **c**, Example of over-polishing. The nuclear mitochondria (NUMT) sequence was transformed as a full mitochondria (MT) sequence during long-read polishing due to the absence of the MT contig, and the NUMT attracted all long-reads from the MT. In comparison, the trio-binned assembly had the MT sequence assembled in place, preventing mis-placing of MT reads during read mapping.

**Extended Data Table 1.**
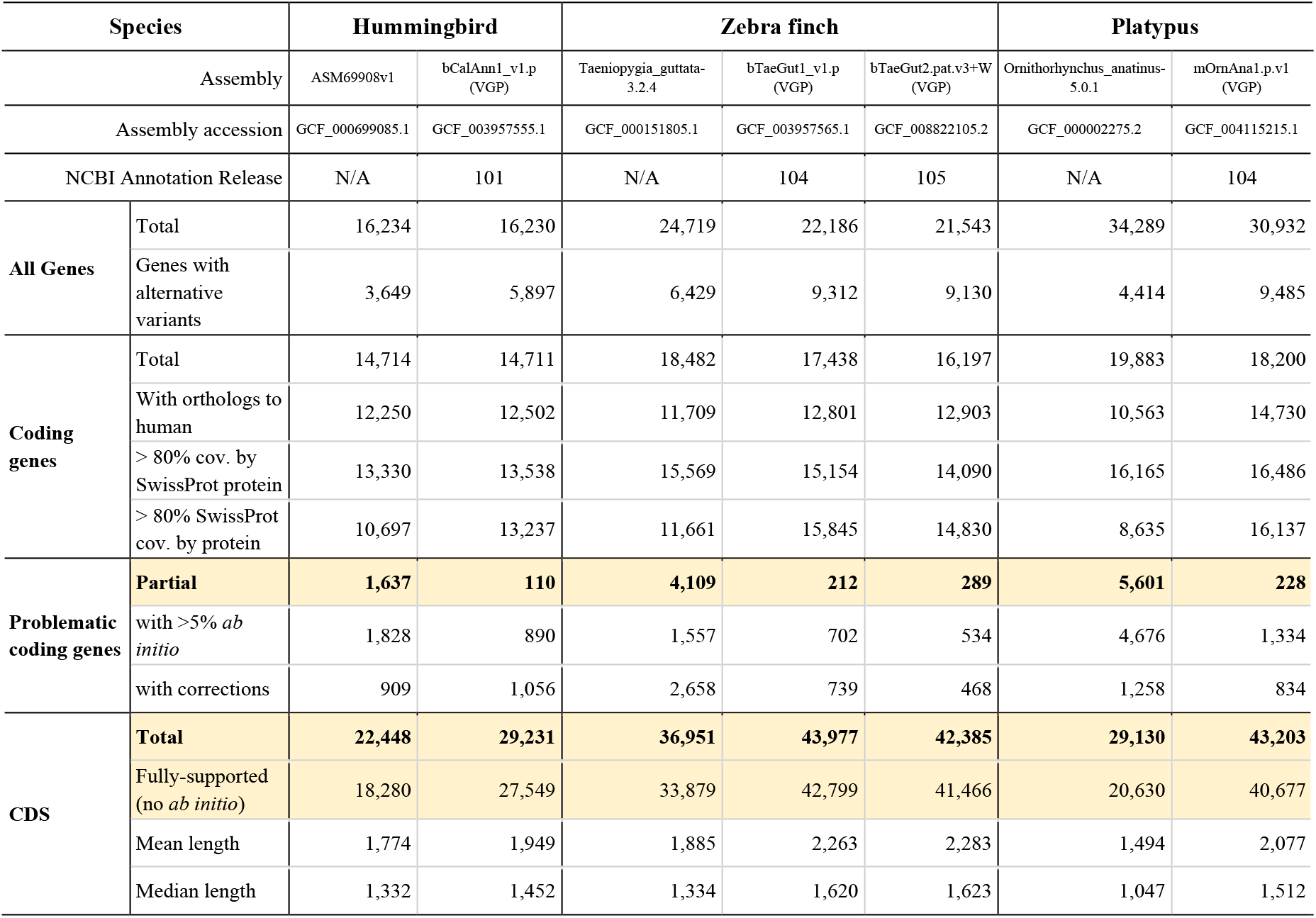
Annotation summary statistics in previous and newly assembled VGP reference genomes. Annotation results of VGP assemblies and previous reference assemblies with the NCBI Eukaryotic Genome Annotation Pipeline, using the same RNA-Seq data and nearly identical sets of transcripts and proteins on input. Highlighted rows are plotted in Fig. 5b.

## Methods

### Genome assembly naming

For each completed assembly of an individual, we gave that assembly an abbreviated name with the following rules: Lineage/GenusSpecies/Individual#.Assembly#. The first letter, in lowercase, identifies the particular lineage: m, mammals; b, birds; r, reptiles; a, amphibians; f, teleost fish; and s, for sharks and other cartilaginous fishes. The next three letters (first in caps) identify the species scientific genus name; the next three letters (first in caps) identifies the specific species name. In the last position is the genome identifier, where integers (1,2,3, …) represent different individuals of the same species, and decimals (1.1, 1.2, 1.3, …) represent different assemblies of the same individual. For example, the first submission of the curated Anna’s hummingbird (*Calypte anna*) assembly is bCalAnn1.1, and an updated assembly of the same individual is bCalAnn1.2. When the abbreviated Lineage/GenusSpecies name for two or more species were identical, we replaced the 4th, 5th, etc, letter of the genus or species name until they could be differentiated. We have created abbreviated names for all 71,657 vertebrate species (http://vgpdb.snu.ac.kr/splist/).

### Sample collection

Producing high-quality genome assemblies required obtaining high quality cells or tissue that would yield high molecular weight DNA for long-read sequencing technologies (CLR and ONT), and optical mapping (Bionano). Thus, we obtained fresh-frozen samples of various tissues (**Supplementary Table 8**). All tissue types tested yielded sufficient quantity and quality DNA for sequencing and assembly, but we found blood worked best for species that have nucleated cells (i.e. bird and reptiles), and spleen or cultured cells worked best for mammals, as of to date. Analysis of different tissue types is presented in our companion study (Dahn et al in preparation).

### Isolation of high molecular weight DNA

#### Agarose plug DNA isolation

For tissue, high molecular weight DNA (HMW DNA) was extracted using the Bionano animal tissue DNA isolation fibrous tissue protocol (cat# RE-013-10; document number 30071), according to manufacturer guidelines. A total of 25-30 mg was fixed in 2% formaldehyde and homogenized using the Qiagen TissueRuptor or manual tissue disruption. For nucleated blood, 27-54 μl was used with an adapted protocol (Bionano personal communication) of the Bionano Prep Blood and Cell Culture DNA Isolation Kit (cat# RE-130-10). Lysates were embedded into agarose plugs and treated with Proteinase K and RNase A. Plugs were then purified by drop dialysis with 1x TE. DNA quality was assessed using pulse field gel electrophoresis (PFGE) (Pippin Pulse, SAGE Science, Beverly, MA) or the Femto Pulse instrument (Agilent). PFGE revealed that we isolated ultra-high molecular weight DNA between ~100 and ~500 kbp long.

#### Phenol-Chloroform gDNA extraction

For some samples, we performed phenol-chloroform extractions for HMW gDNA. Snap-frozen tissue has been pulverized into a fine powder with a mortar and pestel in liquid nitrogen. The powdered tissue was lysed overnight at 55°C in high-salt tissue lysis buffer (400 mM NaCl, 20 mM Tris base (pH 8.0), 30 mM EDTA (pH 8.0), 0.5% SDS, 100 μg/ml Proteinase K), powdered lung tissue was lysed overnight in Qiagen G2 lysis buffer (Cat. No. 1014636, Qiagen, Hilden, Germany) containing 100 μg/ml Proteinase K at 55°C. RNA was removed by incubating in 50 ug/ml RNase A for 1 hour at 37°C. HMW gDNA was purified with two washes of Phenol-Chloroform-IAA equilibrated to pH 8.0, followed by two washes of Chloroform-IAA, and precipitated in ice-cold 100% Ethanol. Filamentous HMW gDNA was either spooled with shepherds hooks or collected by centrifugation. HMW gDNA was washed twice with 70% ethanol, dried for 20 minutes at room temperature and eluted in TE. For the flier cichlid muscle gDNA sample used for PacBio CLR and 10XG libraries, glycogen precipitated was performed by adding 1/10 (v/v) 0.3 M Sodium Acetate, pH 6.0 to the extracted genomic DNA, mixed carefully and spun at room temperature at 10.000 xg. PFGE revealed DNA molecule length was between 50 and 300 kbp, often lower in size than that obtained with the agarose plug but sufficient long range sequencing of CLR and linked read data types.

#### Others

We also used the Qiagen MagAttract HMW DNA kit (cat# #67563) and the KingFisher Cell and Tissue DNA kitr (Thermo Scientific; cat# 97030196), following the manufacturer’s guidelines. These protocols yielded HMW DNA ranging from 30 to 50 kbp. The Genomic Tip (Qiagen) kit was also used for tissue based extraction of HMW DNA.

### Libraries and sequencing

#### PacBio libraries and sequencing

DNA obtained from agarose plugs was sheared down to ~ 40 kbp fragment size with a MegaRuptor^™^ device (Diagenode, Belgium), fragmented using Covaris g-tubes (#520079) or by Needleshearing. PacBio large insert libraries were prepared with either the SMRTbell Template Prep Kit 1.0-SPv3 (#100-991-900) or the SMRTbell Express Template Prep Kit v1 (#101-357-000). Libraries were size-selected between 12 and 25 kbp using Sage BluePippin (Sage Science, USA), depending on the DNA quality and extraction method. These libraries were sequenced on either RSII, or Sequel I instruments at least 60X coverage per species using Sequel Binding Kit and Sequencing Plate versions 2.0 and 2.1 with 10 hours movie time (**Supplementary Table 9**).

#### 10X Chromium libraries and sequencing

Unfragmented HMW DNA from the agarose plugs were used to generate linked read libraries on the 10X Genomics Chromium platform (Genome Library Kit & Gel Bead Kit v2 PN-120258, Genome Chip Kit v2 PN-120257, i7 Multiplex Kit PN-120262) following manufacturer guidelines. We sequenced the 10X libraries at ~60X coverage per species on an Illumina NovaSeq S4 150bp PE lane.

#### Bionano libraries and optical map imaging

Unfragmented uHMW DNA from the agarose plugs were labeled using either two different nicking enzymes (BspQI and BssSI) or a direct labelling enzyme (DLE1) following the Bionano Prep Labeling NLRS (document Number 30024) and DLS protocols respectively (document Number 30206). Labelled samples were then imaged on a Bionano Irys or on a Bionano Saphyr instrument. For all species, we aimed for at least 100X coverage per label (**Supplementary Table 9**).

#### Hi-C libraries and sequencing

Chromatin interaction (Hi-C) libraries were generated using either Arima Genomics, Dovetail Genomics, or Phase libraries on muscle, blood, or other tissue with *in-vivo* cross-linking (**Supplementary Table 9**) and sequenced on Illumina instruments. Arima-HiC preparations were performed by Arima Genomics (https://arimagenomics.com/) using the Arima-HiC kit that uses two enzymes (P/N : A510008). The resulting Arima-HiC proximally-ligated DNA was then sheared, size-selected around ~200-600bp using SPRI beads, and enriched for biotin-labeled proximity-ligated DNA using streptavidin beads. From these fragments, Illumina-compatible libraries were generated using the KAPA Hyper Prep kit (P/N: KK8504). The resulting libraries were PCR amplified and purified with SPRI beads. The quality of the final libraries was checked with qPCR and Bioanalyzer, and then sequenced on Illumina HiSeq X at ~60X coverage following the manufacturer’s protocols. Dovetail-HiC preparations were performed by Dovetail using a single enzyme (DpnII) proximity ligation approach. Phase-HiC libraries were made by Phase Genomics using a Proximo Hi-C Library single enzyme reaction.

### Quality control

Before performing any assembly, all genomic data of all data types from each sample were used to screen potential outlier libraries, outlier sequencing runs, or accidental species contamination with Mash^82^ by measuring sequence similarity (**Supplementary Fig. 2**). When running Mash, 21-mers were used to generate sketches with sketch size of 10000 and compared among each sequencing run, and then differences assessed between sequencing sets.

### Genome size, repeat content, and heterozygosity estimations

These estimations were made with *k*-mer-based methods applied to the Illumina short reads obtained from 10XG linked sequencing libraries. After trimming off barcodes during scaff10x preprocessing, canonical 31-mer counts were collected using Meryl^48^. With the resulting 31-mer histogram, GenomeScope^47^ was used to estimate the haploid genome length, repeat content, and heterozygosity. The thorny skate linked read data failed QC, which we suspect was due to low complexity sequences from the high repeat content (54.1%) of the genome; so *k*-mers were collected later from Illumina whole genome sequencing reads instead. The genome size of the channel bull blenny was estimated from an alternative method that looks at the mode of long read overlap coverage, as the estimated genome size from GenomeScope was almost doubling the known haploid genome size (1.29 Gbp vs. 0.6 Gbp).

### Benchmarking assembly steps with the Anna’s hummingbird

For developing the VGP Standard Pipeline, various scaffolding, gap filling, and polishing tools were compared. Default options were used unless otherwise noted. Detailed software versions are listed in **Supplementary Table 2**.

#### Contigging and Scaffolding

FALCON^29^ and FALCON-Unzip^33^ (smrtanalysis 3.0.0) was used to generate contigs that used CLR. Canu^83^ 1.5+67 was used to generate the combined CLR and ONT assembly. To benchmark scaffolding with linked reads, we used scaff10x 2.0. For the linked read only assembly, Supernova 2^84^ was used. For the optical maps, two enzyme hybrid scaffolding was used in the Bionano Solve v3.2.1 software, using BspQI and BssSI initially, as well as DLE1 later when the technology was developed. For benchmarking Hi-C in scaffolding, Salsa 2.2^36^ was used for scaffolding results in **Fig. 1a**, with Hi-C reads generated from Arima Genomics. Additional comparisons for the Hi-C libraries were performed using assemblies provided by Dovetail Genomics and Phase Genomics (**Supplementary Table 3**). We used Hi-C from Arima Genomics as it had the least number of PCR duplicates and better coverage for short and long interactions at the time of comparison (**Supplementary Fig. 1**). Assembly statistics from HiRise, Proximo HiC, 3D-DNA^85^ with Arima Hi-C are available in **Supplementary Table 3**. We concluded all Hi-C scaffolding algorithms had similar performance. We decided to use Salsa as HiRise and Proximo HiC were not open access, and 3D-DNA was not computationally scalable on the DNAnexus platform. For short read assemblies, other than Supernova and the NRGene assembly, the assembly GCA_000699085.1^19^ was used for benchmarking, which was generated with Illumina paired-end, multiple mate-pair libraries and the SoapDeNovo assembler. The NRGene assembly was provided by the company with DeNovo Magic.

#### Gap filling

We ran PBJelly with `support --capturedOnly --spanOnly` parameters, to avoid gap greedy closures with no spanning read support. For conservatively filling sequences, we compared different parameters `output` stage with `--minreads 1` and `--minreads 4` in addition to no restrictions. We found the number of gaps closed was similar to the gaps filled with Arrow (**Supplementary Table 6**) and chose not to run PBJelly for future assemblies.

#### Short-read polishing

Illumina polishing benchmarking was performed using Longranger 2.1.3 and Pilon^86^ 1.21 with `--fix bases, local option` (**Supplementary Table 7**). Later, for the VGP Pipeline, we used FreeBayes^37^ as Pilon^86^ was not computationally scalable for large genomes with the updated Longranger 2.2.2.

#### Base-level accuracy estimate

Base-level accuracy was measured using mapping based approach and later using the k-mer based approach. For mapping based QV estimation to determine number of rounds to polish, Illumina paired-end reads from Zhang et al^19^ were used.

#### Mis-joins and missed-joins

The curated hummingbird assembly was mapped to the target assemblies with MashMap2^57^ with `--filter_mode one-to-one --pi 95` using 5 kbp segments (`-s 5000`) for CLR assemblies and 1 kbp (`-s 1000`) for SR assemblies to compensate for the shorter contig sizes, as contigs smaller than a segment size will be excluded from the alignment. The number of mis-joins and missed-joins were identified using the assembly_comparison.pl used in curation (**Supplementary Methods**).

### VGP standard genome assembly pipeline 1.0 to 1.6

All 17 genomes were assembled with the VGP pipeline (**Extended Data Fig. 2a**) for benchmark purposes, with some uncurated. The pale spear-nosed bat, greater horseshoe bat, Canada lynx, platypus, male and female zebra finch, kākāpō, Anna’s hummingbird, Goode’s thornscrub tortoise, flier cichlid, and the blunt-snouted clingfish assemblies were generated using the **VGP Pipeline 1.0** to **1.6** and curated for submission to NCBI and EBI public archives. The curated and submitted two-lined caecilian, zig-zag eel, climbing perch, channel bull blenny, eastern happy, and the thorny skate assemblies were generated with a similar process developed in parallel (**Supplementary Note 2**). Two submitted curated versions of the female zebra finch were made, one using the standard VGP pipeline and the other using the VGP trio pipeline, so that comparative analyses can be performed by others.

#### Contigging

For PacBio data, contigs were generated from subreads using FALCON^29^ and FALCON-Unzip^33^, with one round of Arrow polishing (smrtanalysis 5.1.0.26412). A minimum read length of 2 kbp or a cutoff where reads longer than the cutoff includes 50x coverage was used, whichever was longer. Estimated genome size for calculating the read coverage was used from http://www.genomesize.com/ when available, or from the literature (**Supplementary Table 11**) while waiting for 10XG sequencing to estimate genome size using *k-mers*. FALCON and FALCON-Unzip were run with default parameters, except for computing the overlaps. Raw read overlaps were computed with DALIGNER parameters `-k14 -e0.75 -s100 -l2500 - h2 4 0 -w8` to better reflect higher error rate in early PacBio sequel I and II. Pread (preassembled read) overlaps were computed with DALIGNER parameters `-k2 4 -e.90 -s100 -11000 - h600` intending to collapse haplotypes for the FALCON step to better unzip genomes with high heterozygosity rate. FALCON-Unzip outputs both a pseudo-haplotype and a set of alternate haplotigs representing the secondary alleles. We refer to these outputs as the primary contig set (c1) and alternate contig set (c2).

#### Purging false duplications

We discovered that FALCON-Unzip incorrectly retains some secondary alleles in the primary contig set, which appear as false duplications. To reduce these false duplications, we ran purge_haplotigs^27^, first during curation (VGP v1.0 pipeline) and then later after contig formation (VGP v1.5 pipeline). To do the former, purge_haplotigs was run on the primary contigs (c1), and identified haplotigs were mapped to the scaffolded primary assembly with MashMap2^57^ for removal. In the latter, identified haplotigs were moved from the primary contigs (c1) to the alternate haplotig set (p2). The remaining primary contigs were referred to as p1; p2 combined with c2 was referred to as q2. Later in the VGP v1.6 pipeline we replaced purge_haplotigs with purge_dups^51^, a new program developed by several of the authors in response to purge_haplotigs not removing partial false duplication at contig boundaries. Purging also removes excessive low coverage (junk) and high coverage (repeats) contigs.

#### Scaffolding with 10XG linked reads

The 10X Genomics linked reads were aligned to the primary contigs (p1), and an adjacency matrix was computed from the barcodes using scaff10x v2.0-2.1 (https://github.com/wtsi-hpag/Scaff10X). Two rounds of scaffolding were performed. The first round was run with parameters `-matrix 2000 -reads 12 -link 10`, and the second round with parameters `-matrix 2000 -reads 8 -link 10`. A gap of 100 bp (represented with ‘N’s) was inserted between joined contigs. The resulting primary scaffold set was named s1.

#### Scaffolding with Bionano optical maps

Bionano cmaps were generated using the Bionano Pipeline in non-haplotype assembly mode and used to further scaffold the s1 assembly with Bionano Solve v3.2.1. We began with a one enzyme nick map (BspQI), followed by two enzyme nick map (BspQI and BssSI), and then with a DLE-1 one enzyme non-nicking approach when the later data type became available (**Supplementary Table 9**). Scaffold gaps were sized according to the software estimate. The resulting scaffold set was named s2.

#### Scaffolding with Hi-C reads

Hi-C reads were aligned to the s2 scaffolds using the Arima Genomics mapping pipeline (https://github.com/ArimaGenomics/mapping_pipeline). In brief, both ends of a read pair were mapped independently using BWA-MEM^87^ with the parameter -B8, and filtered when mapping quality was <10. Chimeric reads containing a restriction enzyme site were trimmed from the restriction site onward, leaving only the 5’ end. The filtered single read alignments were then rejoined as paired read alignments. The processed alignments were then used for scaffolding with Salsa2^36^, which analyzes the normalized frequency of Hi-C interactions between all pairs of contig ends to determine a likely ordering and orientation of each. We used parameters `-m yes -i 5 -p yes` to allow Salsa2 to break potentially mis-assembled contigs and perform 5 iterations of scaffolding. After feedback from curation, a later versions of Salsa were developed, which more conservatively determines number of iterations (v2.1) and actively breaks at mis-assemblies (v2.2), and run for the Canada lynx, Goode’s thornscrub tortoise, and two-lined caecilian. The restriction enzyme(s) used to generate each library were specified using parameters ‘-e GATC,GANTC’ for Arima and ‘-e GATC’ for Dovetail and Phase Genomics Hi-C data. The resulting Hi-C scaffolded assembly was named s3.

#### Consensus polishing

To polish bases in both haplotypes with minimal alignment bias, we concatenated the alternate haplotig set (c2 in v1.0 or q2 in v1.5~1.6) to the scaffolded primary set (s3) and the assembled mitochondrial genome (mitoVGP in v1.6). We then performed another round of polishing with Arrow (smrtanalysis 5.1.0.26412) using PacBio CLR reads, aligning with pbalign `--minAccuracy=0.75 --minLength=50 --minAnchorSize=12 -- maxDivergence=30 --concordant --algorithm=blasr --algorithmOptions=--useQuality --maxHits=1 --hitPolicy=random -- seed=1` and consensus polishing with variantCaller `--skipUnrecognizedContigs haploid -x 5 -q 20 -X120 -v --algorithm=arrow`. While this round of polishing resulted in higher QV for all genomes herein considered, we noticed that it is particularly sensitive to the coverage cutoff parameter (-x). This is due to the fact that Arrow generates a *de novo* consensus from the mapped reads without explicitly considering the reference sequence. Later, we found that the second round of Arrow polishing sometimes reduced the QV accuracy for some species. Upon investigation, this issue was traced back to option `-x 5`, which requires at least 5 reads to call consensus. Such low minimum requirements can lead to uneven polishing in low coverage regions. To avoid this behavior, we suggest to increase the -x close to the half sequence coverage (e.g. 30x when 60x was used for assembly) and check QV before moving forward.

For genomes with a combined assembly size larger than 4 Gbp, we used Minimap2 with parameters `-ax map-pb` instead of Blasr to overcome reference index size limitations.

Two more rounds of base-pair polishing were performed with linked reads. The reads were aligned with Longranger align 2.2.2, which incorporates the Lauriat for barcode-aware alignment^88^. From the alignments, homozygous mismatches (variants) were called with FreeBayes^37^ v1.2.0 using default options. Consensus was called with bcftools consensus^89^ with – `i’QUAL>1 && (GT=“AA” || GT=“Aa”)’ -Hla`.

#### VGP Trio Pipeline v1.0 ~ v1.6

The Trio Pipeline is similarly designed to the Standard Pipeline, except the use of parental data (**Extended Data Fig. 2b**). When parental genomes are available, the child’s CLR reads are binned to maternal and paternal haplotypes, and assembled separately as haplotype-specific contigs (haplotigs) using TrioCanu^53^. In brief, parental specific marker k-mers are collected using Meryl from the parental Illumina WGS reads of the parents. These markers were filtered and used to bin the child’s CLR read. A haplotype was assigned given the markers observed, normalized by the total markers in each haplotype. The subsequent purging, scaffolding, and polishing steps were similarly updated with the use of purge_dups^51^ (v1.6). We extended binning to linked reads and Hi-C reads, by excluding read pairs having any parental specific marker. The binned Hi-C reads were used to scaffold its haplotype assembly, and polished with the binned linked reads from the observation of haplotype switching using the standard polishing approach. During curation, one of the haplotype assemblies with the higher QV and/or contiguity was chosen as the representative haplotype. The heterogametic sex chromosome from the unchosen haplotype was added to the representative assembly.

For the female zebra finch in particular, contigs were generated before the binning was automated in the Canu assembler as TrioCanu1.7, and thus a manual binning process was applied as described in the original Trio-binning paper^53^ (**Supplementary Methods**). Contigs were assembled for each haplotype using the binned reads, excluding unclassified reads. The contigs were polished with two rounds of Arrow polishing using the binned reads, and scaffolded following the v1.0 pipeline with no purging. Additional scaffolding rounds with Bionano (s4) and Hi-C were applied. Scaffolds were renamed according to the primary scaffold assembly of the same individual (s5), with sex chromosomes grouped as Z in the paternal assembly and W in the maternal assembly following synteny to the Z chromosome from the curated male zebra finch VGP assembly. Two rounds of SR polishing were applied using linked reads, by mapping on both haplotypes. After haplotype switches were discovered, additional rounds of polishing were applied using binned linked reads (**Supplementary Methods**).

#### Mitochondrial genome assembly

Similar to other recent methods^90,91^, we developed a reference-guided MT assembly pipeline. MT reads in the raw CLR data were identified by mapping the whole read set to an existing reference sequence of the specific species or of closely related species using Blasr. Filtered mtDNA CLR were assembled into a single contig using Canu v1.8, polished with Arrow using CLR and then FreeBayes^37^ v1.0.2 together with bcftools v1.9 using short reads from the 10XG data (**Extended Data Fig. 2**). The overlapping sequences at the ends of the contig were trimmed, and the remaining contig sequence circularized. The mitoVGP pipeline is made available at https://github.com/VGP/vgp-assembly/tree/master/mitoVGP. A more detailed protocol description of the assembly pipeline and new discoveries from the MT assemblies are presented in our companion study (Formenti et al, in preparation).

### Curation

The VGP genome assembly pipeline produces high quality assemblies, yet no automated method to date is free from producing errors, especially during the scaffolding stages. To minimize the impact of the remaining algorithmic shortcomings, we subjected all assemblies to rigorous manual curation. All data generated for a species in this study and other publicly available data (e.g. genetic maps, gene sets and genome assemblies of the same or closely related species) were collated, aligned to the primary assembly and analyzed in gEVAL^38^ (https://vgp-geval.sanger.ac.uk/index.html), visualizing discordances in a feature browser and issue lists. In parallel, Hi-C data was mapped to the primary assembly and visualized using Juicebox^92^ and/or HiGlass^93^. With this data, genome curators identified mis-joins, missed joins and other anomalies, and corrected the primary assembly accordingly. No change was made without unambiguous evidence from available data types; for example, a Hi-C suggested join would not be made unless supported by BioNano maps, long-read data, or gene alignments.

#### Contamination removal

A succession of searches was used to identify potential contaminants in the generated assemblies.

1. A megaBLAST^94^ search against a database of common contaminants (ftp://ftp.ncbi.nlm.nih.gov/pub/kitts/contam_in_euks.fa.gz) requiring e-value ≤ 1e-4, reporting matches ≥ 98% sequence identity and match length 50-99 bp, ≥ 94 % and match length 100-199 bp, or ≥ 90% and match length 200 bp or above.
2. A vecscreen (https://www.ncbi.nlm.nih.gov/tools/vecscreen/) search against a database of adaptor sequences (ftp://ftp.ncbi.nlm.nih.gov/pub/kitts/adaptors_for_screening_euks.fa)
3. After soft-masking repeats using Windowmasker^95^ a megaBLAST search against chromosome-level assemblies from RefSeq requiring e-value ≤ 1e-4, match score >= 100, and sequence identity ≥ 98%; regions matching highly conserved rDNAs were ignored.

Manual inspection of the results was necessary to differentiate contamination from conservation and/or horizontal gene transfer. Adaptor sequences were masked; other contaminant sequences were removed. Assemblies were also checked for runs of Ns at the ends of scaffolds, created as artefacts of the iterative scaffolding process, and when found they were trimmed.

#### Organelle genomes

These were detected by a megaBLAST search against a database of known organelle genomes requiring e-value ≤ 1e-4, sequence identity ≥ 90%, and match length ≥ 500; the databases are available at ftp://ftp.ncbi.nlm.nih.gov/blast/db/FASTA/mito.nt.gz and ftp://ftp.ncbi.nlm.nih.gov/refseq/release/plastid/*genomic.fna.gz. Only scaffolds consisting entirely of organelle sequences were assumed to be organelle genomes, and replaced by the genome from the separate organelle assembly pipeline. Organelle matches embedded in nuclear sequences found to be NuMTs were kept.

#### False duplication removal

Retained false duplications were identified using purge_haplotigs^27^ run on either post scaffolding and polishing (Anna’s hummingbird, kākāpō, male zebra finch, female zebra finch, platypus, pale spear-nose bat, and greater horseshoe bat) or on the c1 before scaffolding (two-lined caecilian, flier cichlid, Canada lynx, and Goode’s thornscrub tortoise). Subsequent manual curation identified additional haplotypic duplications for the listed assemblies and also those that were not treated with purge_haplotigs^27^ (Eastern happy, climbing perch, zig zag eel). The evidence used included read coverage, sequence self-comparison, transcript alignments, Bionano map alignments and Hi-C 2D maps, all confirming the superfluous nature of one allele. The identified haplotype duplications were moved from the primary to the alternate assembly.

#### Chromosome assignment

For a scaffold to be annotated as a chromosome, we used evidence from Hi-C as well as genetic linkage or FISH karyotype mapping when available. For Hi-C evidence, we considered a scaffold as a complete chromosome (albeit with gaps) when there was a clear unbroken diagonal in the Juicebox or HiGlass plots for that scaffold and no other large scaffolds that could be joined to that same scaffold; if present and no unambiguous join possible, we named it as an unlocalised scaffold for that chromosome. When we could not find evidence of a complete chromosome, we kept the scaffold number for its name. We named all evidence-validated scaffolds as chromosomes down to the smallest Hi-C box unit resolution allowed with these characteristics. When there was an established chromosome terminology for a given species or set of species, we use the established terminology except when our new assemblies revealed errors in the older assembly, such as scaffold/chromosome fusions, fissions, rearrangements, and non-chromosome names. For species without an established chromosome terminology, we name the scaffolds as chromosomes numbers 1, 2, 3 …, in descending order of scaffold size. For the sex chromosomes, we use the letters X and Y for mammals and Z and W for birds.

#### Using comparative genomics to assess assembly structure

In cases where a high-quality chromosome-level genome was available for a closely related species, comparative genome analysis was performed. The polished primary assembly (t3.p) was mapped to the related genome using MashMap2^57^ with `--pi 75 -s 300000`. The number of chromosomal differences were identified using a custom script available at https://github.com/jdamas13/assembly_comparison. This resulted in the identification of ~60 to ~450 regions for each genome assembly flanking putative misassemblies or lineage-specific genome rearrangements. To identify which were real misassemblies, the identified discrepancies were communicated to the curation team for manual verification (see above).

To identify any possible remaining mis-joins, each curated avian and mammalian assemblies were compared with the zebra finch (taeGut2) or human (hg38) genomes, respectively. Pairwise alignments between each of the VGP assemblies and the clade reference were generated with LastZ^96^ (version 1.04) using the following parameters: *C = 0 E = 30 H = 2000 K = 3000 L = 2200 O = 400*. The pairwise alignments were converted into the UCSC “chain” and “net” formats with axtChain (parameters: *-minScore = 1000 -verbose = 0 -linearGap = medium*) followed by chainAntiRepeat, chainSort, chainPreNet, chainNet and netSyntenic, all with default parameters^97^. Pairwise synteny blocks were defined using maf2synteny^98^ at 100, 300, and 500 kbp resolutions. Evolutionary breakpoint regions were detected and classified using an ad hoc statistical approach^99^. This analysis identified 2 to 90 genomic regions per assembly that could be flanking misassemblies, lineage-specific chromosome rearrangements, or reference-specific chromosome rearrangements (116 in the human and 26 in the zebra finch). Determining the underlying cause for each of the flagged regions will need further verification. All alignments are available for visualization at Evolution Highway comparative chromosome browser (http://eh-demo.ncsa.illinois.edu/vgp/).

### Annotation

NCBI and Ensembl annotation pipeline used in this study are described in **Supplementary Methods**.

### Evaluation

Detailed methods for other types of evaluations, including BUSCO runs, mis-join and missed-join identification, reliable blocks, collapsed repeats, telomeres, RNA-Seq and ATAC-Seq mapping, false gene duplications and annotations are in the **Supplementary Methods.**

## Data availability

All raw data, intermediate and final assemblies are publicly available via GenomeArk (https://vgp.github.io/genomeark) and archived on NCBI/EBI BioProject PRJNA489243 (https://www.ncbi.nlm.nih.gov/bioproject/489243) with annotations and browsable on the UCSC Genome Browser (https://hgdownload.soe.ucsc.edu/hubs/VGP/). The final primary assembly from the automated pipeline before curation is browsable on gEVAL (https://vgp-geval.sanger.ac.uk) with all four raw data mappings. The VGP assembly pipeline is available as a stand-alone pipeline (https://github.com/VGP/vgp-assembly) as well as a workflow on DNAnexus (https://platform.dnanexus.com/). A VGP-specific assembly hub portal in the U.C. Santa Cruz browser is available as a gateway to access all VGP genome assemblies and annotations (https://hgdownload.soe.ucsc.edu/hubs/VGP).

## Code availability

All codes used in the VGP Assembly Pipeline and the Trio Pipeline are publicly available at https://github.com/VGP/vgp-assembly/tree/master/pipeline.

## Acknowledgements

We thank the following persons for feedback and support: Rebbecca Johnson, Elinor Karlsson, Kerstin Lindblad Toh, Wang Jun, Ion Korf, Wilfried Haerty, Graham Etherington, Bernardo Clavijo, and Aleksey Komissarov for discussions in the early stages of the project; Rochelle Fuller for help with the G10K website maintenance, and Harry Segal with VGP website development; Memory Linh Pham for help with initial grant writing; Lauren Shalmiyev for administrative help; Deanna Church formerly of 10X Genomics, Guy Kol, Kobi Baruch, and Omer Barad of NRGene, Ivan Liachko of Phase Genomics, Ivo Gut of Nanopore, Evgeny Muzychenko, Shilpa Garg, and Mikhail Kolmogorov for their preliminary analyses performed on one or more genomes; Karen Oliver, Craig Corton and Jason Skelton for data generation; Edward Harry for technical support in scaff10x and Pretext; Camila Mazzoni for coordinating students and training at Leibniz Institute for Zoo and Wildlife Research and Berlin Center for Genomics in Biodiversity Research, Germany; Maximilian Driller, Calvinna Caswara, Majid Vafadar, Nicholas Hill, Diego De Panis, Annabel Whibley, Brigid Maloney, Cailey Mitchell, Guido Gallo, Jonathan Gaige, Kwame Amoako-Boadu, Maria Jose Gomez, Maripaz Montero, Danil Ratnikov, Samara Brown, Sarah Zylka, Stephanie Marcus, Tomás Carrasco for completing training and testing the VGP pipeline by producing ordinal representative genome assemblies not described in this manuscript. We further thank assistance from company partners (listed below), NCBI, EBI, and Amazon AWS, including AWS for sponsoring storage.

AR, SK, BPW and AMP were supported by the Intramural Research Program of the NHGRI, NIH. AR was also supported by the Korea Health Technology R&D Project through KHIDI, funded by the Ministry of Health & Welfare, Republic of Korea (HI17C2098). SAM, IB and RD were supported by Wellcome WT207492, WC, MSm, ZN, YS, JC, SPe, JT, AT, JW and KerH by WT206194, LH, FM, KevH and PFli by WT108749/Z/15/Z, WT218328/B/19/Z and the European Molecular Biology Laboratory. OF and EDJ were supported by Howard Hughes Medical Institute and Rockefeller University start-up funds for this project. JD and HL were supported by the Robert and Rosabel Osborne Endowment. MU-S received funding from the European Union’s Horizon 2020 research and innovation programme under the Marie Skłodowska-Curie grant agreement (750747). FT-N, JHof, PMa and KC were supported by the Intramural Research Program of the NLM, NIH. CL, BK, JKi and HK were supported by the Marine Biotechnology Program of KIMST, funded by the Ministry of Ocean and Fisheries, Republic of Korea (20180430). SCV was funded by a Max Planck Research Group awarded from the Max Planck Society, and a Human Frontiers Science Program (HFSP) Research grant (RGP0058/2016). TML, WEJ and the Canada lynx genome was funded by the Maine Department of Inland Fisheries & Wildlife (F11AF01099), including when WEJ held a National Research Council Research Associateship Award at the Walter Reed Army Institute of Research (WRAIR). CB was supported by NSF (1457541 and 1456612). DB was funded by The University of Queensland. DI was supported by Science Exchange Inc. (555 Bryant Street, #939, Palo Alto, CA 94301). HWD was supported by NSF (OPP-0132032; ICEFISH 2004 Cruise) and PLR-1444167. GN and the thorny skate genome were funded by Lenfest Ocean Program (30884). MP was funded by the German Federal Ministry of Education and Research (01IS18026C). MMal was supported by EMBO fellowship ALTF 456-2016. SSel was supported by NIH R44HG008118, CVM, SRF, PVL by R21 DC014432/DC/NIDCD, KDM by R01GM130691, HC by 5U41HG002371-19, MD by U41HG007234, BP by R01HG010485. DG was supported by National Key Research and Development Program of China. FOA was supported by Al-Gannas Qatari Society and The Cultural Village Foundation-Katara, Doha, State of Qatar and Monash University Malaysia. CT was supported by The Rockefeller University. HC was supported by NHGRI (5U41HG002371-19). RHK was funded by the Max Planck Society with computational resources at the bwUniCluster and BinAC funded by the Ministry of Science, Research and the Arts Baden-Württemberg and the Universities of the State of Baden-Württemberg, Germany (bwHPC-C5). MTPG was supported by ERC Consolidator Award 681396-Extinction Genomics, Danish National Research Foundation Center Grant (DNRF143). ECT was supported by European Research Council (ERC-2012-StG311000) and Irish Research Council Laureate Award. TW was supported by NSF (1458652). J-MG was supported by Australian Research Council. EWM was partially supported by the German Federal Ministry of Education and Research (01IS18026C). Complementary sequencing support for the Anna’s hummingbird and several genomes was provided by Pacific Biosciences, Bionano Genomics, Dovetail Genomics, Arima Genomics, Phase Genomics, 10X Genomics, NRGene, Oxford Nanopore Technologies, Illumina, and DNAnexus. All other sequencing and assembly was conducted at the Rockefeller University, Sanger Institute, and Max Planck Institute Dresden genome labs. Part of this work utilized the computational resources of the NIH HPC Biowulf cluster (https://hpc.nih.gov). We acknowledge funding from the Wellcome Trust (108749/Z/15/Z) and the European Molecular Biology Laboratory. We thank Le Comité Scientifique Régional du Patrimoine Naturel and Direction de l’Environnement, de l’Aménagement et du Logement, Guyanne for research approvals and export permits.

## Author contributions

Wrote the paper and co-coordinated the study: A.R., E.D.J, A.M.P, R.D., E.W.M, Ker.H., S.A.M., O.F.

Coordination with vendors: J.Ko., S.Sel., R.E.G., A.H., M.Mo.

Collected samples: M.T.P.G, W.E.J, R.W.M, G.Z., B.V., M.B., J.How., S.C.V., T.M.L., F.G., W.C.W., D.B., M.J.Ge., M.B., D.I., A.D., D.E., T.E., M.Wi., G.T., A.Me., A.F.K., P.Fr., H.W.D., H.S., M.Wa., G.J.P.N., R.D., E.D.J.

Generated genome data: O.F., I.B., M.Sm., B.H., J.Mo., S.W., C.B., A.Me., A.F.K., P.Fr., D.F.C., C.V.M

Generated genome assemblies: A.R., S.A.M., S.K., M.P., S.B.K., R.H., J.G., Z.N., J.L., B.P.W., M.Ma.

Generated/modified softwares: S.K., A.R., S.B.K., R.H., Z.K., J.Ko., I.S., C.D., Z.N., A.H., J.L., J.G., E.G., C.V.M., S.R.F.

Pipeline development: A.R., S.A.M., G.F., S.K., M.U-S., A.F., M.Si., B.T.H., T.P., M.P., E.W.M., R. D., A.M.P.

Generated MT assemblies: G.F., J.Ko.

Curation: Ker.H., W.C., Y.S., J.C., S.Pe., J.T., A.T., J.W., Y.Z., J.D., H.L.

Sex chromosomes: Y.Z., R.S.H., K.D.M., P.Me., J-M.G.

Hummingbird karyotype analyses: M.H., A.Mi., M.P., E.W.M., E.D.J. Annotation: F.T-N., L.H., J.Hof., P.Ma., K.C., F.M., Kev.H., P.Fl., D.B.

Evaluation analysis: A.R., J.D., M.U-S., G.L.G., L.J.C., F.T-N., L.H., C.L., BJ.K., J.Ki., J.M.Ge., J.G., D.G., S.E.L., D.F.C., C.V.M., S.R.F., P.V.L., E.O., F.O.A-A., S.Sec., C.T., H.K., Ker.H., E.W.M., R.D., A.M.P, E.D.J.

Data availability: A.R., S.A.M., W.C., A.F., S.Pa., M.Si., B.T.H., B.P.W., W.K., H.C., M.D., L.N., B.P., A.M.P., E.D.J.

G10K council, founders, and coordination of VGP: T.M-B., A.J.C., F.D-P., R.D., M.T.P.G., E.D.J., K-P. K., H.L., R.W.M., E.W.M., E.C.T., B.V., G.Z., A.M.P., S.Pa., J-M.G., O.A.R., D.H., S. J.O.

All authors reviewed the manuscript.

