## Supplementary Information for "Towards complete and error-free genome assemblies of all vertebrate species"

**Supplementary Note 1**
**Supplementary Note 2**
**Supplementary Note 3**
**Supplementary Note 4**
**Supplementary Methods**

### Supplementary Note 1

This supplementary note contains additional findings that we believe will be useful for the genome community. Some of the results were generated during evaluations with genome company technology engineers and bioinformaticians, including several co-authors, which we actively engaged with feedback on what we and they discovered using the genomic richness of the species in this study.

#### Haplotype phasing

Most of the contiging and scaffolding tools we used were originally designed to handle a haploid representation of the genome. As shown in this study, this design can result in errors in and around heterozygous alleles. We found that such errors introduced from haplotype differences early in the process propagates to later stages, and are not easily removable. The earlier the haplotypes are sorted in the assembly pipeline, the higher the assembly quality metrics especially for highly heterozygous genomes, as seen here with both haplotypes of the female zebra finch assembly in the following order of increasing metric quality: 1) Collapsing first and phasing afterwards (e.g. FALCON-Unzip[1]); 2) Collapsing first and phasing afterwards using Hi-C (e.g. FALCON-Phase[2]); and 3) Phasing reads before assembling into contigs (e.g. TrioCanu[3]). Below we present additional findings.

During curation, we found a pattern of excessive breaks flanking heterozygous sites, which contributed to false duplications in the primary assembly. We traced the source and found FALCON often unnecessarily broke the contigs at the branching point between runs of homozygosity and pairs of heterozygous alleles, reducing the continuity of the assembly. This case is illustrated in types 3 and 4 in Box 1a, leaving many homotype duplications at contig boundaries as described in Box 1b (left). In response, PacBio fixed the problem in an updated software release (smrtanalysis upgrade, April 2018), which doubled the contig N50 sizes on a number of genomes.

Haplotypes are essentially a chromosome-scale genomic repeat, and so new methods developed for repeat separation should also help the haplotype assembly problem. The fundamental challenge is distinguishing true genomic variants from errors in the sequencing data, and paralogs from orthologs, and then linking those solutions across the length of full chromosomes. This includes needed tools that can better model the diploid (or polyploid) architecture of the genome by integrating long-range evidence from multiple sources, across large repeats, while still preserving haplotype specific variation in the genome.

#### Optical mapping

Bionano Genomics used our Anna's hummingbird and Kakapo samples to help develop and test their 2-enzyme nicking (BspQI and BssSI) and 1-enzyme non-nicking (DLE-1) approaches for hybrid scaffolding. In early 2015, we found that using two sets of nicking enzyme maps together resulted in better scaffolding continuity compared to using only one (data not shown). This was because the two enzymes compensate each other and eliminate scaffold breaks. Later in 2017, in the development of DLE-1, we found the molecule sizes were superior with avoiding unintentional cutting of genomic DNA at label sites. When applying to the same FALCON-Unzip primary contigs, we confirmed the scaffold NG50s were better in the DLE-1 only approach compared to the 2-enzyme hybrid approach (**Supplementary Table 3**). Therefore, we decided to move forward with the latest DLE-1 technology whenever possible.

**Comparisons of Hi-C data types and scaffolding**

As mentioned in the main text, we tested Hi-C and Chicago libraries on the same Anna's hummingbird sample from three sources (Dovetail Genomics Hi-C v1 and Chicago v1, Phase Genomics v1, and Arima Genomics v1) with versions developed as of mid-2016. We mapped back the paired Hi-C reads using Juicer[4] to the previous reference hummingbird assembly generated with short reads[5], and evaluated interaction size distribution, duplication rate, and genome coverage. We caution that this mapping depends on the structural and base call accuracy of the prior assembly, but all Hi-C datasets were at least being compared to the same assembly. We found that insert size (linking distance) differed: Arima > Phase > Dovetail Hi-C > Dovetail Chicago. Dovetail Hi-C tended to have more paired end reads without an insert, which with our feedback they fixed in an upgraded chemistry at the time. Phase Genomics Hi-C had more PCR duplicates, which were possible to screen out. Arima had the highest per base coverage for phasing (**Supplementary Fig. 1**), presumably due using two enzymes instead of one at the time, but the overall genome assembly was not distinguishable between the data sets. Based on these analyses, we chose to use Arima Hi-C v1 to generate most of the assemblies on other species for this study. We note, though, that each company has continued to make improvements, and thus choice of Hi-C data type will need continued evaluation.

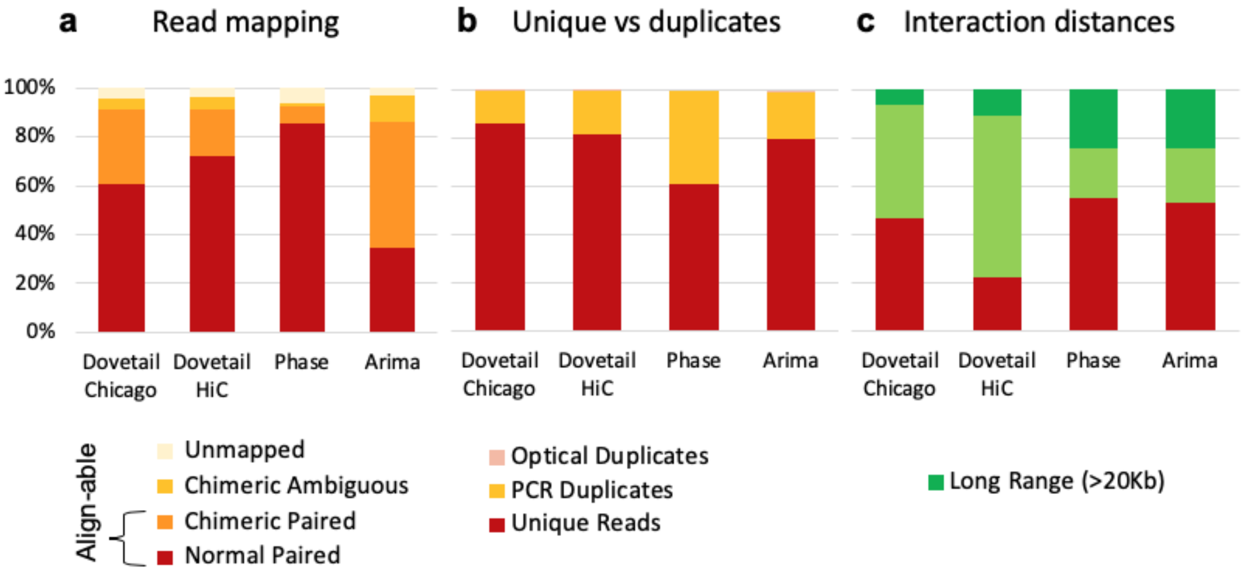


**Supplementary Fig. 1 | Read mapping statistics and interaction distance of three Hi-C platforms benchmarked on Anna's hummingbird assembly. a,** Read % mapped back to the reference; the higher the alignable portion of the reads mapped the more useful for assembly. **b**, The higher the proportion of unique mapped reads the more useful for assembly; duplicates do not add new information. **c**, The higher the proportion of long range interactions distances between Hi-C read pairs, greater than the long read lengths (~20Kb for CLR) and spanning contigs, the more useful for chromosomal scale scaffolding. These data sets are from v1 chemistries from each company (Dovetail Genomics, Phase Genomics, and Arima Genomics) used on the samples from the same Anna's hummingbird, mapped back to its original short read reference[5]. Each company has made improvements since, including based on the results of this figure and associated data that we provided.

We tested different Hi-C scaffolding algorithms (Phase Genomics Proximo Hi-C; Dovetail Chicago HiRise; 3D-DNA[6], and Salsa[7]). We found that with software versions used at the time, Salsa 2.0 resulted in the highest NG50 metrics with apparent over-scaffolding when tested on the hummingbird Hi-C data sets. With feedback from curation of the hummingbird and other assemblies, we also improved Salsa to Salsa2.2, with a feature that breaks mis-joins introduced in prior scaffolding rounds with support from Hi-C interactions. These changes simplified and streamlined downstream visualization of the assemblies. As with the Hi-C data types, developers of these algorithms continue to make improvements, including with our iterative feedback and may thus work differently than the versions we tested.

### Gap filling

We attempted to use PBJelly[8] on the hummingbird, a tool developed using a previous G10K supported assembly of the Assemblathon 2.0 budgerigar[8] and other genomes to fill in gaps between contigs in scaffolds with PacBio CLR reads. However, we found during evaluation that in addition to properly filling gaps, PBJelly introduced many erroneous base errors in the gap-filled consensus when using default options. The default scaffolding function also introduced mis-joins. Subsequently, we tested Arrow (smrtanalysis 5.1.0.26412 version) on these same assemblies, a consensus base caller that also has a gap filling function, developed by Pacific BioSciences (https://github.com/PacificBiosciences/GenomicConsensus)[9]. We found running PBJelly with conservative parameters (**Supplementary Table 6**) had similar results to Arrow. In fact, Arrow was more conservative and had better consensus quality of the filled gaps. With this result, we replaced PBJelly with Arrow in our initial pipeline.

### Polishing

We initially attempted to use Pilon[10] for Illumina SR polishing, and found the best combination of polishing to get the highest QV with minimal steps (**Supplementary Table 7**). However, Pilon was memory intensive to run on large genomes. Thus, we switched to FreeBayes and bcftools for variant calling and consensus generation. During our initial FreeBayes polishing, we encountered regions with excessive coverage (>4000x). We worked with the original author of FreeBayes (E.G. of this study)[11] and found it was hanging on low complexity regions or regions with excessive high coverage. The low complexity issue was fixed by changing the coding in the entropy calculation steps; the excessive coverage issue from loading reads was fixed by setting an upper limit of the coverage (release v1.3.1). These fixes also resulted in optimized memory usage and speed. In all cases of polishing, the reads were mapped to both the primary and alternate haplotype assemblies and the better of the two mappings selected as the primary to avoid reverting haplotype-specific variants.

### Comparative iterative scaffolding

When using Hi-C scaffolding alone, we found there were a higher number of inversion errors for the smaller contigs as reported in Ghurye et al[7] (**Supplementary Table 3**), which can be difficult to orient correctly using long range data alone. Optical mapping with the non-nicking DLE-1 chemistry yielded the most accurate scaffolding, but similar to Hi-C, optical mapping had a limited ability to scaffold short contigs due to the smaller chance of containing sufficient labeled enzyme sequence sites to confidently align the optical cmaps. These small contigs can be difficult to handle in later assembly stages, because most scaffolding tools were not designed to place a contig within an existing scaffold gap. Together, 10XG linked reads were better at localizing smaller contigs, complementing what optical maps missed. We found a few cases where scaff10x erroneously mis-joined contigs due to the ambiguity at the contig boundaries near repeats; however, because subsequent scaffolding and curation steps can break such errors and it is difficult to manually place short sequences, we kept the linked read scaffolding step before the optical map scaffolding step.

### Supplementary Note 2

The submitted reference versions of five of the fish genomes (eastern happy, climbing perch, channel bull blenny, zig-zag eel, and thorny skate) and the two-lined caecilian were assembled using modified approaches (**Supplementary Table 10**).

**The two-lined caecilian, aRhiBiv1.1 (GCA_901001135.1) assembly**: This assembly mostly follows the VGP 1.5 standard pipeline, however run at the Sanger Institute without having a formal functional equivalence evaluation between this setup and the centralised VGP setup. Versions used and differences in the pipeline are as follows: FALCON-Unzip[1] (v1.2.1), Purge haplotigs[12] (v1.0.1), scaff10x (v3.0) (we ran two rounds of scaff10x followed by one round of break10x, which differs from the main VGP 1.5 pipeline), Bionano Solve (v3.2.2), SALSA2[7] (v2.2), Arrow (GenomicConsensus 2.2.2), longranger (v2.2.2), freebayes (v1.1.0-3-g961e5f3) and bcftools consensus (v1.7). Manual curation was applied as described elsewhere and chromosome-scale scaffolds confirmed by the Hi-C data were named in order of size.

**The zig-zag eel, fMasArm1.2 (GCA_900634775.2) assembly**: FALCON-Unzip (v1.8.6) contigs were created using a length cutoff of 6500. The primary contigs were extended by merging with a miniasm (0.2-r168) assembly. The contigs were then scaffolded using the 10XG linked read Illumina data with scaff10x (v1.0). Further scaffolding was applied using synteny with Lates calcarifer (v3 from http://seabass.sanbi.ac.za/) and the cross_genome tool (https://sourceforge.net/projects/phusion2/files/cross_genome/). PBJelly (PBSuite_15.8.24) was used to fill gaps followed by long read polishing with Arrow (GenomicConsensus 2.2.1). The assembly was polished again using the linked reads by mapping with bwa mem (0.7.17-r1188), calling homozygous non-reference variants with freebayes (v1.1.0-3-g961e5f3) and editing the reference to correct these errors with bcftools consensus (v1.7). This assembly was manually curated with incorporating evidence from Bionano optical map and Arima Hi-C data. This initial fMasArm1.1 (GCA_900634775.1) was submitted as scaffolds. An additional run of SALSA (v2.0) was applied, followed by another round of manual curation to remove heterotypic duplications to produce chromosome-level scaffolds. These chromosome-level scaffolds were named based on synteny to a medaka genome assembly and submitted as fMasArm1.2 (GCA_900634775.2).

**The climbing perch, fAnaTes1.2 (GCA_900324465.2) assembly**: An initial PacBio contig assembly was made using FALCON-Unzip (v1.8.6) using a length cutoff of 10000. The primary contigs were then scaffolded using the 10XG linked read data with two rounds of scaff10x (v1.0) followed by a round of break10x to break at mis-joins identified by the 10XG data. PBJelly (PBSuite_15.8.24) was used to fill gaps followed by long read polishing with Arrow (GenomicConsensus 2.2.1). The assembly was polished again using the 10XG Illumina data by mapping with bwa mem (0.7.17-r1188), calling homozygous non-reference variants with freebayes (v1.1.0-3-g961e5f3) and editing the reference to correct these errors with bcftools consensus (v1.7). Manual curation incorporated evidence from Bionano optical map and Arima Hi-C data, and the initial fAnaTes1.1 (GCA_900324465.1) assembly was submitted as scaffolds. An additional run of SALSA (v2.0) was applied, followed by another round of manual curation to remove heterotype duplications using Purge Haplotigs (v1.0) to produce chromosome-level scaffolds. These chromosome-level scaffolds were named based on synteny to a medaka genome assembly and submitted as fAnaTes1.2 (GCA_900324465.2).

**The channel bull blenny, fCotGob3.1 (GCA_900634415.1) assembly**: An initial PacBio assembly was made using Falcon-unzip (falcon-2018.03.12-04.00) without Dazzler repeat-masking during overlap detection. A separate wtdbg (v1.1) assembly was made from the PacBio reads. Contigs from the wtdbg assembly were used to guide initial scaffolding of the Falcon contigs using cross_genome, then scaffolded further with the 10XG Illumina data and scaff10x (v1.0). The Bionano optical map data was used for two-enzyme hybrid scaffolding (Solve3.2.2_08222018). The PacBio CLR data was used to gap fill with PBJelly (PBSuite_15.8.24) and polish with Arrow (GenomicConsensus 2.2.2). The assembly was polished again using the 10XG Illumina data by mapping with bwa mem (0.7.17-r1188), calling homozygous non-reference variants with freebayes (v1.1.0-3-g961e5f3) and editing the reference to correct these errors with bcftools consensus (v1.7). The assembly failed to meet the VGP contig NG50 goals, so a new strategy was tried to improve the assembly. Canu v1.6 was used to correct the PacBio reads using kmer k=21. Contigs from a wtdbg (v1.1) assembly of the corrected reads were then used to conservatively fill gaps in the main assembly where contigs were unambiguously anchored on either side of a gap. Long-read and short-read polishing as performed above were applied again to ensure sequence that had been used to fill gaps was also polished. Retained haplotigs were identified and removed with Purge Haplotigs (v1.0). Finally, the assembly was scaffolded to chromosomes using Arima Hi-C data and SALSA (v2.0). Manual curation was applied using gEVAL as described elsewhere to correct mis-joins and improve concordance with the Bionano optical map data and Arima Hi-C data. This assembly met the VGP metrics, and chromosome-scale scaffolds were named based on synteny to a medaka genome assembly.

**The eastern happy, fAstCal1.2 (GCA_900246225.3) assembly**: First the PacBio raw reads were scrubbed to remove chimeric reads and other artifacts using the Dazzler framework (https://dazzlerblog.wordpress.com/2017/04/22/1344/). The scrubbed reads were then used to make an initial assembly with PacBio Falcon-unzip (https://github.com/millanek/FALCON-integrate). A separate assembly was created with miniasm (0.2-r159), then used to scaffold the Falcon primary contigs using cross_genome. The contigs were then scaffolded further using the 10XG Illumina data with scaff10x (v1.0). Some contigs in the scaffolds were gap filled with PBJelly (PBSuite_15.8.24) and polish with Quiver (GenomicConsensus 2.2.1). The assembly was manually curated using gEVAL to correct mis-joins and improve concordance with the Bionano optical map data. The assembly was then polished again using the 10XG Illumina data by mapping with bwa mem (0.7.17-r1188), calling homozygous non-reference variants with freebayes (v1.1.0-3-g961e5f3) and editing the reference to correct these errors with bcftools consensus (v1.7). This assembly was submitted as fAstCal1.1 (GCA_900246225.1). A further round of curation and verification with gEVAL along with integration with two genetic maps[13,14] allowed assignment of scaffolds to chromosomal linkage groups. This was submitted as revised assembly fAstCal1.2 (GCA_900246225.3).

**The thorny skate, sAmbRad1.pri (GCA_010909765.1) assembly:** Applying the VGP 1.0 pipeline to the thorny skate did not result in an assembly the met all the desired VGP metrics, due to the very high repeat content in this species, as described in the main text. Therefore, we developed a modified approach that can handle high repeat genomes better. We used Canu v1.7 contigs purged with purge_haplotigs instead of the purged FALCON-Unzip contigs, because the contig NG50 and overall BUSCO completeness score were higher in the Canu contigs (**Supplementary Table 13**, compare p1 stats of the vgp_standard_1.5 and vgp_nhgri_1.5). We believe these differences could be due to the low repeat masking in the Canu assembler. The rest of the scaffolding process followed the VGP Standard Pipeline 1.5. Two rounds of Arrow polishing was applied (t1), with 3 rounds of SR polishing (t2~t4) with linked reads using longranger align 2.2.2 and freebayes 1.3.1 `--skip-coverage` option to skip regions with excessive coverage. Too many false duplications were found during curation, and thus the assembly was sent back for further improvements. We applied purge_dups on t4 primary scaffold, by breaking scaffolds at any gaps. Purged primary contigs (u1) were re-scaffolded with optical maps (u2) and further scaffolded with 2 rounds of Salsa (u3-u4). We discovered linked read library failure, thus obtained additional Illumina WGS reads. Using the WGS reads, 3 rounds of polishing was performed with bwa and freebayes.

### Supplementary Note 3

**Missing genomic regions**

Although the VGP genomes have a great amount of more genomic content assembled relative to the most commonly used prior references when compared (e.g. **Extended Data Fig. 6**), we also noted that some of them had a small proportion of missing genomic content found in prior assemblies, which may need further improvements or explanation. An example was the initial mitochondrial genomes (MT), which we added by performing a separate MT assembly process. We investigated further the repeat masking process and tried various length cutoffs to see if these missing regions are recovered.

Almost all genome assemblers mask out reads with repeat structure before assembly, as it becomes computationally expensive or impossible to assemble with repeats present. FALCON masks portions of reads which coincide with repetitive regions, and does so in two stages: 1) tandem repeat masking; and 2) masking of general repeats/segmental duplications. The masked repetitive regions then do not contribute to the overall overlap computation. For large repeats (longer than the read length), this means that the genomic region will not be represented in the final assembled contigs. In addition, and not related to repeat masking, FALCON also applies a cutoff threshold to limit the minimum length of reads to find overlaps. If the limit is set too high, the assembly may miss some genomic regions. After the initial contig phase, these repeats as well as smaller reads are brought back into the assembly for Arrow polishing and later gap filling. However, we found that certain reads with repeats and a given read size were not being incorporated into the assembly if they did not have a region to anchor onto in the initial contigs. An example were some genes with GC-rich sequence and repeat regions of the Anna's hummingbird, that were present on reads shorter than 10,000 bp. We surmised that these non-repetitive genes were surrounded by GC-rich and repeat genomic regions, which are relatively fragile[15] to obtain long molecules compared to the rest. These shorter and/or repetitive reads were excluded from overlap detection, or ignored due to the shorter overlap length with no anchor to bring them into the assembly at later stages. When we reduced the pread cut off to 2,000 bp, the NG50 values decreased, but many of the genes on these shorter and repetitive reads were incorporated into the assembly. This highlights the need of further investigation and improvements in ways to rescue missing regions when applying general length cutoffs during the assembly process.

### Supplementary Note 4

An implementation of the pipeline has been developed to run on generic architecture using WDL workflows (https://github.com/openwdl/wdl/blob/master/versions/1.0/SPEC.md) and Docker containers (https://www.docker.com/). We intend for this to be a portable and modular implementation, which diverges from the main workflow as little as possible.

WDL (Workflow Description Language) is a standard which enables the description of workflows in both human- and machine-readable ways. Workflows are composed of tasks; tasks have defined inputs and outputs, a script to perform the work, hardware requirements, and an environment in which to be run. Docker is a virtualization tool that we use to provide the environment for tasks. A Docker container is a lightweight image of a filesystem that can contain specific tool versions. It uses a layered filesystem, where multiple snapshots can inherit from a single base image.

The design of the VGP's WDL workflow implementation aims to replicate current functionality while minimizing changes to the main codebase. The main codebase is designed to run in an HPC environment and uses CEA-HPC Modules (https://github.com/cea-hpc/modules) to manage use of specific tool versions. The WDL implementation replicates this environment in the Docker images, so as to reduce modification to the existing scripts. There is a base Docker image which includes the modules infrastructure, common libraries, and tools used in multiple tasks. Task-level Docker images extend from this and add task-specific code. Slurm submission scripts from the original pipeline were rewritten and translated into WDL tasks. Operative bash scripts (the entrypoints Slurm uses) are copied into the task images and are invoked directly where possible.

Scaffolding and QC tasks have been implemented in WDL/Docker:
Contigging+purging, linked read scaffolding, optical map scaffolding, hi-c scaffolding, BUSCO, and Merqury[16]. Each task can be run independently, and the whole scaffolding suite can be run via a single workflow. For information on running the workflow, see the manual here:
https://github.com/VGP/vgp-assembly/blob/master/wdl_pipeline/WDL_Manual.md

### Quality control and contamination screening

Before assembly of the sequence and scaffold data, a quality control screening for poor sequencing reactions or contamination with foreign genome data was performed using Mash. When running Mash[17], 21-mers were used to generate sketches with sketch size of 10,000 and compared among each sequencing runs. For example, using this approach, we detected two outlier libraries in the Canada lynx PacBio data that did not cluster with the other sequencing runs (**Supplementary Figure 2**). Further investigation determined that these files had been mis-tracked and the data originated from an unrelated sequencing project on rice, so the rice runs were removed prior to assembly.

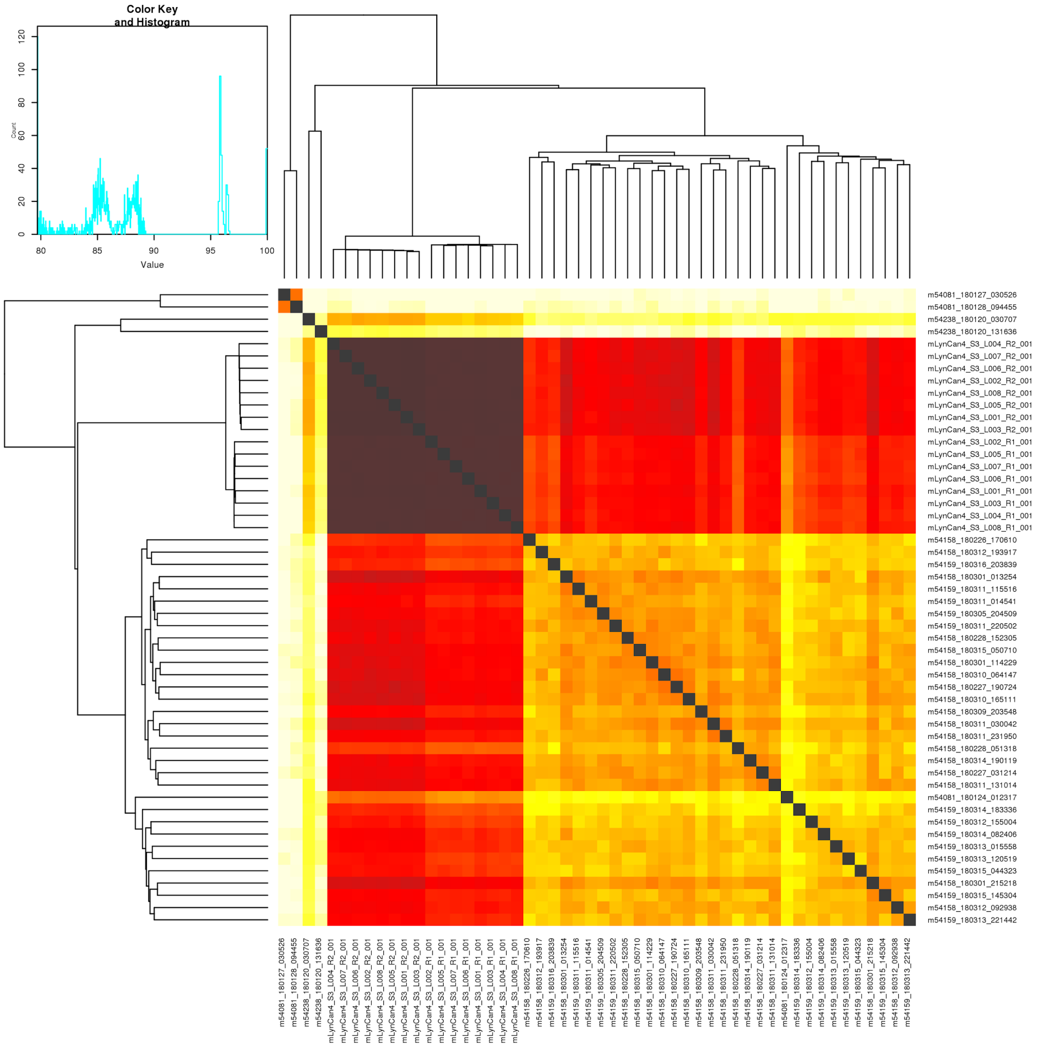


**Supplementary Fig. 2 | Mash maps to detect poor quality data and foreign species contamination.** Raw read data of PacBio CLR (top left 4 and the bottom right) and linked reads (darker brown area) are aligned to each other. Based on sequence similarity and coverage, the top two (top left corner) SMRT cells were identified as outliers. The higher the similarity and coverage, the darker the color.

### Supplementary Methods

#### Binning 10XG linked reads and Hi-C reads

Linked reads and Hi-C reads were binned using an "exclusion" criteria. Unlike trio-binning[3], here we exclude any read-pair having at least one parental specific k-mer when assigning to a bin. This allows short reads to remain in both maternal and paternal bins, which originate from the homozygous part of the genome. This process uses `meryl-lookup` Meryl[16] v1.0:

```
meryl-lookup -memory 2 -exclude -mers pat.meryl -sequence $read1 -
sequence2 $read2 -r2 mat.R2.fastq.gz | pigz -c > mat.R1.fastq.gz
meryl-lookup -memory 2 -exclude -mers mat.meryl -sequence $read1 -
sequence2 $read2 -r2 pat.R2.fastq.gz | pigz -c > pat.R1.fastq.gz
```

An implemented version is available on Merqury:
https://github.com/marbl/merqury/blob/master/trio/exclude_reads.sh

**ENSEMBL annotation pipeline**

The Ensembl gene annotation system[24] was used to generate annotation for the high-quality assemblies. Annotation was created primarily through alignment of transcriptomic data to the genome, with gap filling via protein to-genome alignments of a select set of vertebrate proteins from UniProt[25] and, for mammal species, coordinate mapping of GENCODE[26] human reference annotations via a pairwise whole genome alignment.

The transcriptomic data consisted primarily of short read RNASeq data sourced from the public archives. Where available, Nanopore and PacBio Iso-Seq were long read data were also used. Short read data were initially mapped via BWA[27] and then locally re-aligned in a splice-aware manner via Exonerate[28]. Transcripts were then inferred based on the strongest intron/exon signals for likely genic loci on a per tissue basis. Long-read data were mapped to the genome using Minimap2[21] with the recommended settings for Iso-Seq and Nanopore data. Due to the high error rate of the Nanopore data, post mapping error correction was employed to maximize the number of usable mappings. Intron/exon boundaries that were non-canonical or deemed low frequency (five or fewer observations across all mappings at a locus) were replaced with high frequency boundary coordinates (greater than five observations) within a 50bp edit distance. High frequency boundary observations were determined both from canonical boundary observations from the Nanopore mapping themselves and also from the alignments of the short-read data. A similar strategy was employed to remove likely artificial gaps of 200bp or less from exons described by the Nanopore data. In these cases, low frequency potential gaps between two adjoining exons were filled in based on high frequency observations of single exons with the same terminal boundary coordinates. For each transcript model generated from either short or long read data, the longest open reading frame was assessed via a BLAST[19] of UniProt vertebrate proteins that had experimental evidence of existence at either the protein level or the transcript level.

For gap filling, where the transcriptomic data were absent or fragmented, homology-based methods were employed. Splice-aware protein-to-genome alignments were carried out via GenBlastG[29]. Annotation mapping from human was carried out via a pairwise alignment using LastZ[30] (https://etda.libraries.psu.edu/catalog/7971) and subsequent exon coordinate mapping and transcript reconstruction via both in-house software and CESAR 2.0[31].

At each locus, low quality transcripts (in particular those with evidence of a fragmented ORF) models were removed, and the data collapsed and consolidated into a final gene model plus its associated non-redundant transcript set. ORF likelihood was determined by aligning the ORF translation against known vertebrate proteins. Priority was given to models derived from transcriptomic data. For loci where the transcriptomic data was not available or highly fragmented, homology data took precedence, with preference given to longer transcripts that had strong intron support from the short-read data. Summary statistics of the annotations are available in the annotation reports in **Supplementary Table 20**.

**NCBI annotation pipeline**

The NCBI Eukaryotic Genome Annotation Pipeline was used to annotate genes, transcripts, and proteins on the primary assembly of the 17 assemblies, submitted between September 2018 and April 2020, and the product of the annotations were added to the RefSeq collection. The genome sequences were masked using Windowmasker[18]. RNA-Seq reads were retrieved from SRA for the species, or for the family, in the case of lynx, kakapo, Anna's hummingbird, Goode's thornscrub, blunt snouted clingfish and thorny skate for which no or insufficient amount of RNA-Seq data was found at the species level. Depending on the species, between 385 million and 10 billion RNA-Seq reads, ESTs, and RefSeq and GenBank transcripts for the species or closely related species were aligned to the masked genome using BLAST[19] followed by Splign[20]. PacBio IsoSeq for lynx, kakapo, Anna's hummingbird, zebra finch and Oxford Nanopore Technologies transcriptomics reads for kakapo were aligned to these species' respective assemblies using minimap2[21]. In addition, human RefSeq proteins and GenBank and RefSeq proteins for related organisms were aligned to the genome using Blast and ProSplign. The gene models' structures and boundaries were obtained with Gnomon(NCBI eukaryotic gene prediction tool. Available at: https://www.ncbi.nlm.nih.gov/genome/annotation_euk/gnomon/) by "chaining" the alignments into preliminary models. Partial open reading frames on these chained alignments (missing either a start or a stop) were joined and filled if adjacent and in compatible frames, or extended by the ab initio module of Gnomon using a hidden Markov model trained on the species, if the coding propensity of the region was sufficiently high, or called non-coding. tRNAs were predicted with tRNAscan-SE:1.23[22] and small non-coding RNAs were predicted by searching the RFAM 12.0 HMMs for eukaryotes using cmsearch from the Infernal package[23].

Gene and transcript models were then evaluated at each locus, and one of overlapping gene models was chosen in this order of precedence: same-species curated RefSeq models, RFAM models, Gnomon models, and finally tRNA models. Among Gnomon models, if multiple fully-supported transcript variants were predicted for a gene, only the models supported in their entirety by a single long alignment (e.g., a full-length mRNA) or by RNA-Seq reads from a single BioSample were selected. Poorly supported Gnomon models conflicting with better-supported models annotated on the opposite strand were excluded from the final set of models. Further filtering of gene models, and assignment of function, name and type to the final accepted gene set was based on orthology to human genes (or zebrafish in the case of the climbing perch) and Blast hits to SwissProt or, as a last resort, Blast hits to nr. Gnomon models with high homology to transposable or retro-transposable elements or models that appear to be single-exon retrocopies of protein-coding genes were excluded from the final set of models. Most Gnomon models with base differences with the genomic assembly introduced to correct frameshift-causing indel were labeled as pseudogenes and annotated without a CDS feature or protein product; but those with a strong unique hit to the SwissProt database or with a human ortholog were marked coding. Such models may indicate underlying defects in the assembly and should be considered lower confidence. Titles for these models are prefixed with "PREDICTED: LOW QUALITY PROTEIN". The resulting annotated assemblies and the annotated products for all 17 assemblies were loaded into RefSeq and are publicly available for download from NCBI Assembly (https://www.ncbi.nlm.nih.gov/assembly/). For each assembly, details of the evidence used for gene prediction and summary statistics of the annotations are available in the annotation reports in **Supplementary Table 20**.

**Chromosome size estimate from Karyotype imaging**

The Anna's hummingbird karyotype image was processed first into a binary representation. Then, the rectangular area surrounding each chromosome image was obtained from the binary representation. The relative chromosome size ratio was estimated compared to the sum of all rectangular heights. Each chromosome size estimate was obtained by multiplying this ratio to the given genome size.

The genome size is given in diploid, as both alleles are present in the Anna's hummingbird karyotype picture. Below is the python code used to generate the karyotype image and chromosome size estimates:

```
# Import relevant libraries and setup matplotlib
import numpy as np
from skimage.measure import regionprops
from skimage.color import rgba2rgb, rgb2gray
from scipy.ndimage import label
import matplotlib.pyplot as plt
import matplotlib.patches as mpatches
import argparse
%matplotlib inline

plt.rcParams['figure.figsize']=20,15

# genome size in bases - if image shows multiple alleles, then the size
has to be adapted accordingly
gs = 2119374518

# be sure to remove anything from the picture (i.e. labels) that is not
to the karyotype
path_img_in = "/Users/pippel/Documents/Calypte_anna_in.png"
path_img_out = "/Users/pippel/Documents/Calypte_anna_out.png"
path_txt_out = "/Users/pippel/Documents/Calypte_anna_out.txt"

# loading the image
img = plt.imread(path_img_in)
plt.imshow(img)

# converting to binary
threshold = 0.5  #<---------- threshold can/should be adjusted
if img.shape[2] == 4:
    gray = 1-rgb2gray(rgba2rgb(img))
else:
    gray = 1-rgb2gray(img)

binary = (gray>threshold).astype(np.int32)
plt.imshow(binary)

# creating labels
labels = label(binary)[0]

plt.imshow(labels)
```

```
# regionprobs on the label image
regions = regionprops(labels)

sizes=sum(r.area for r in regions)

fig, ax = plt.subplots(figsize=(100, 60))
ax.imshow(binary)

plt.rcParams.update({'font.size': 32})

centroid_and_area = np.zeros((len(regions), 3))
c=0
for region in regions:
    x, y = region.centroid
    area = region.area
    minr, minc, maxr, maxc = region.bbox
    # draw rectangle around objects
    rect = mpatches.Rectangle((minc, minr), maxc - minc, maxr - minr,\
     fill=False, edgecolor='red', linewidth=2 \
     plt.text(minc, minr,                     \
     str(np.round(gs*region.area/sizes.sum()/1000000,2)), color='red')
    centroid_and_area[c] = (x, y, gs*region.area/sizes.sum()/1000000)
    c+=1
    ax.add_patch(rect)

# save image to image output file
plt.savefig(path_img_out)

# save centroid and size estimate to text file
np.savetxt(path_txt_out, centroid_and_area, fmt='%1.3f %1.3f %1.3fM')
```

**Weighted read length distributions**
All read lengths or molecule lengths were collected for PacBio CLR and Bionano optical maps over 1kb. The weighted read length distribution was calculated using the read length normalized by the total bases sequenced in reads of that length.

**Pacbio CLR read length distribution:** Read length was extracted for all subreads with samtools[24] 1.9-1.10 `faidx` and filtered for reads over 1kb. Total bases were counted at the end to weight the bases in the read length of X.

```
# Collect read length for each subreads
samtools faidx $subreads.fasta

# length collected per genome at the end
cat *.fasta.fai | awk '{print $1"\t"$2"}' > $genome.len

# Remove data < 1kb and change unit to 1kb
awk '$2>1000 {print $2/1000}' $genome.len > $genome.1kb

# Get the fraction of a read of length LEN over TOTAL_BP
TOTAL_BP=`awk -v sum=0 '{sum+=$1} END {print sum}' $genome.1kb`
awk -v TOTAL_BP=$TOTAL_BP -v PLATFORM="PacBio" \
 '{print $1"\t"($1/TOTAL_BP)"\t"PLATFORM}' $genome.1kb \
  > $dir/$genome.1kb.weighted
```

**10XG molecule length distribution:** 10X Genomics molecule length distribution was estimated using the `molecule_length_mean` (m) from the `summary.csv` produced with `longranger align`. We ran `longranger align` on all curated primary assemblies (**Supplementary Table 10**) except for the Skate, which we used the longranger align output (m) from the 3rd round of polishing. For the two-lined caecilian (aRhiBiv1), the largest chromosomes were broken in two scaffolds and lifted over at the end to bypass the known indexing issue in longranger for large scaffolds. We used the m for length-weighted mean molecule length and computed the exponential distribution of the molecule length following 10X Genomics recommendation on https://support.10xgenomics.com/de-novo-assembly/software/pipelines/latest/output/moleculelen. In brief, when b = 2/m, the function f(x) = b * exp(-bx), where x is the molecule length.

**Bionano raw molecule length distribution:** Molecule length was extracted for all .bnx files over 1kb. Lengths were weighted by total bases.

Following code was applied to Opt1 (BspQI), Opt2 (BssSI), and Opt3 (DLE1) bnx files to extract the molecule length:

```
name=`basename $file | sed 's/.bnx//g' \
       | awk -F "_" '{print $1"_"$2"_"$3}'`
cat $file | grep -v "#" | grep -v "QX" \
   | awk -v name=$name '{printf "%.0f\t%s\n", $3, name}' \
    >> $name.bnx.len
```

```
# Remove data < 1kb and change unit to 1kb
awk '$1>1000 {print $1/1000}' $bn >> $genome.1kb

# Get the fraction of a read of length LEN over TOTAK_BP, add it to the
PacBio results
TOTAL_BP=`awk -v sum=0 '{sum+=$1} END {print sum}' $genome.1kb`
awk -v TOTAL_BP=$TOTAL_BP -v PLATFORM=$PLATFORM          \
  '{print $1"\t"($1/TOTAL_BP)"\t"PLATFORM}' $genome.1kb \
  >> $dir/$genome.1kb.weighted
```

**Hi-C chromatin interaction length distribution:** Interaction distances between any two Hi-C read pairs were collected using the curated primary assemblies (**Supplementary Table 10**) as the reference. Hi-C reads were mapped using Arima mapping pipeline (https://github.com/VGP/vgp-assembly/blob/master/pipeline/salsa/arima_mapping_pipeline.sh) as we did for Salsa scaffolding. After the alignment was finished, 10 million interactions to the longest scaffold was extracted using BEDTools[25]. As with the other datatypes, the sum of all interaction distances was used to normalize each distance.

```
# Largest scaffold is $scaffold and is $len long
len=`awk -v l=0 '$2>l {l=$2} END {print l}' $genome.pri.fasta.fai`
scaffold=`awk -v len=$len '$2==len {print $1}' $genome.pri.fasta.fai`

# Collect intervals >1kb
bedtools bamtobed -bedpe -i $genome.pri.bam \
  | awk -v scaff=$scaffold '$1==scaff && $4==scaff' \
  | awk '{ if($2<$5) {start=$2;} else {start=$5;} \
     if ($3<$6) end=$6;} else { end=$3 } interval=(end-start); \
     if (interval>1000) {print (interval/1000)}}' > $dir/$genome.1kb
head -n 10000000 $genome.1kb > $genome.1kb.10M
TOTAL_BP=`awk -v sum=0 '{sum+=$1} END {print sum}' $genome.1kb.10M`
awk -v TOTAL_BP=$TOTAL_BP -v PLATFORM=$PLATFORM \
  '{print $1"\t"($1/TOTAL_BP)"\t"PLATFORM}' $genome.1kb.10M \
  > $genome.1kb.10M.weighted
```

**Plotting length distribution:** For 10X linked read distances, we made a `linked.len` file which contains the genome id and the m in two columns. In R, we read the data and generate the distribution for plotting.

```
dat.10x=fread("linked.len", header=T)
result <- data.frame()
for (g in dat.10x$genome) {
  dat_genome1=dat.10x[dat.10x$genome==g,]
  dat_genome1 <- data.frame(Count=seq(from = 1, to = 100000, by = 1),\
     Length=rexp(100000, rate=2/dat_genome1$molecule_length_mean))
  totalbp=sum(as.numeric(dat_genome1$Length))
```

```
  print(totalbp)
  head(dat_genome1)
  dat_genome1$Weight=dat_genome1$y/totalbp
  dat_genome1$Platform="Linked_reads"
  dat_genome1$Genome=c(g)
  result <- rbind(result, dat_genome1)
}
```

For other platforms, the `$genome.1kb.weighted` files were concatenated with the genome id at the end in the order we want to display per genomes:

```
for genome in $(cat genome.list.srt);
do
  awk -v genome=$genome '{print $0"\t"genome}' $genome.1kb.weighted \
    >> all.1kb.weighted
done
```

**BUSCO**

Busco 3.02[26,27] was run on all submitted primary assemblies (**Supplementary Table 12**) to assess gene content (C: completeness; D: Duplications; F: fragmented; M: missing), as well as on the benchmark assemblies (**Supplementary Table 13**) to assess duplications using OrthoDB v9 with vertebrata_odb9 database. Integrated software versions used were Hmmer 3.1b2, ncbi-blast-2.2.30+ and augustus-3.3. Command line used is as following:

```
run_BUSCO.py -i asm.fasta -o $out -m genome -l vertebrata_odb9
```

In addition, we performed lineage specific (-l) BUSCO runs with closest available lineages, which uses a gene prediction model trained on human genome annotations (augustus). Specifically, we used 'laurasiatheria' for mammals, 'ave' for birds, 'tetrapoda' for Goode's thornscrub tortoise, and two-lined caecilian, 'actinopterygii' for fishes, and 'vertebrata' for the thorny skate (**Supplementary Table 12**). Command line used is as following with specific `$lineage`:

```
run_BUSCO.py -i asm.fasta -o $out -m genome -l $lineage
```

When investigating duplications, we further tried BUSCO using species specific models (-sp): 'human' for all mammals and the thorny skate; 'chicken' for all birds, Goode's thornscrub tortoise, and two-lined caecilian; and 'zebrafish' for all fishes (**Supplementary Table 12-13**). Command line used is as following with `-sp` set in addition to above example:

```
run_BUSCO.py -i asm.fasta -o $out -m genome -l $lineage -sp $species
```

We observed very similar patterns in duplication level (**Supplementary Table 13**), however fluctuating slightly in the completeness score (**Supplementary Table 12**). This could reflect the quality of the gene models used for training, reference quality used to generate the initial BUSCO gene set, or the availability of the closest / identical lineage and species. With that, throughout this manuscript, we decided to use vertebrata_odb9 with the default human lineage to use the same gene set for investigating relative completeness.

**Mis-joins and missed-joins in assemblies**

The curated hummingbird assembly was mapped to the target assemblies with MashMap2[28] using 5kb segments for CLR assemblies. We used 1kb segments for SR assemblies to compensate for the shorter contig sizes. We ran MashMap using the following command line:

```
# For CLR assemblies
mashmap -r $ref -q $qry -t $SLURM_CPUS_PER_TASK -o $out \
  --filter_mode one-to-one --pi 95 -s 5000

# For SR assemblies
mashmap -r $ref -q $qry -t $SLURM_CPUS_PER_TASK -o $out \
  --filter_mode one-to-one --pi 95 -s 1000
```

The number of mis-joins and missed-joins were identified using a custom script available at https://github.com/jdamas13/assembly_comparison. The assembly_comparison.pl was run using the command line:

```
perl assembly_comparison.pl out.map $ref.fai $segment
```

Note that `assembly_comparison.pl` requires the more continuous assembly to be the `$qry`, and the target assembly being the `$ref` when running `mashmap2`.

Where the `$segment` adjusted accordingly to the `-s` used in MashMap.

From the summary, the number of end-to-end joins (missed-joins), rearrangements, and free-end joins were collected from the diff.summary using the following script:

```
echo -e \
"target\tmissed-joins\trearrangements\tfree-
end_breaks\tnum.scaffolds_affected_by_free-end_breaks" > summary.txt
cat $summary | awk '{print $NF}' | tr '\n' '\t' |\
 awk -v summary=$ref '{print summary"\t"$1"\t"$2"\t"$4"\t"$5}' \
 >> summary.txt
```

The number of mis-joins consist of two error types (**Supplementary Fig. 3**): 1) rearrangements and inconsistency between the curated and target assembly; and 2) free-end breaks where the target assembly has a join not supported by the curated assembly. Missed-joins are contigs or scaffolds in the target assembly that are joined in the curated assembly.

Similarly, to generate comparisons between assembly pipeline steps within a species (**Supplementary Table 14**), each intermediate assembly was mapped to its predecessor using Mashmap2 with parameters --pi 95 -s 10000.

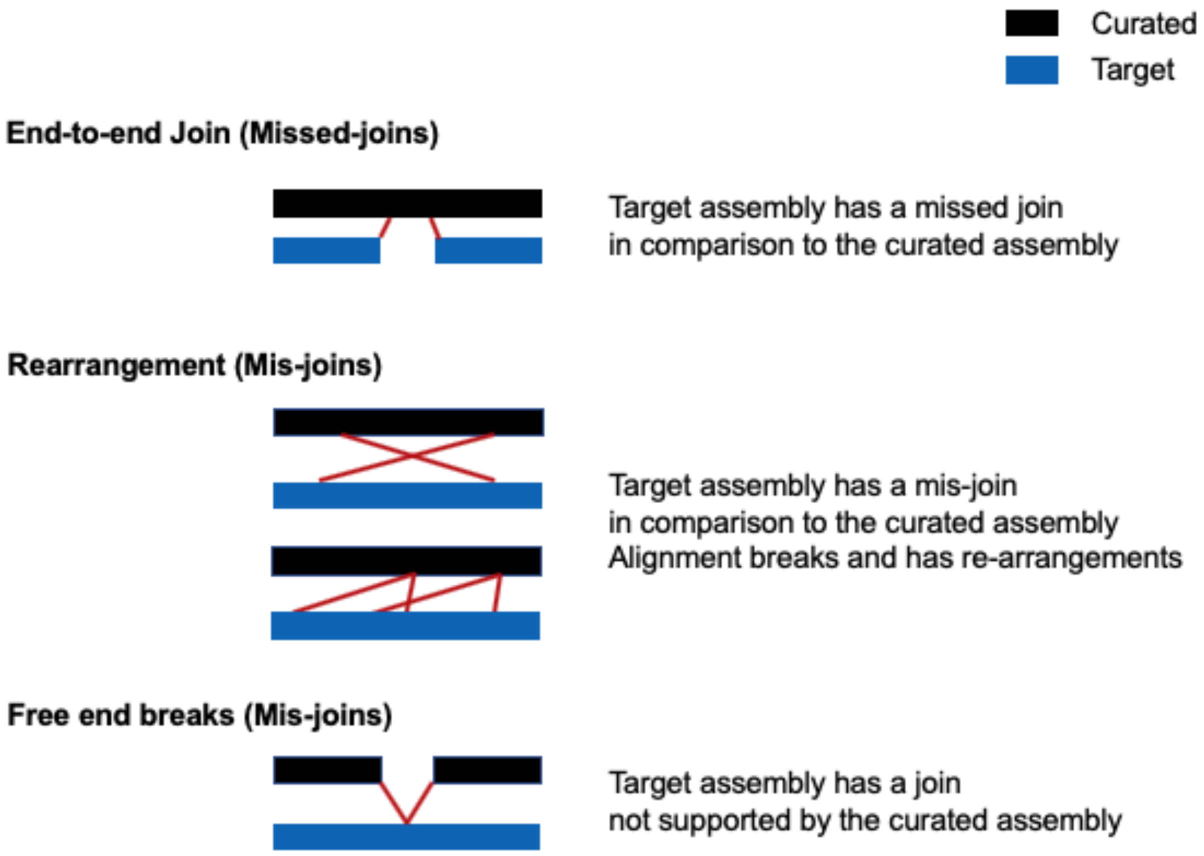


**Supplementary Fig. 3** | Schematic overview of “Mis-joins” and “Missed-joins”.

**Quantifying false duplications with *k-mers***

All *21-mers* were collected from linked reads. The first 23 bp of the first read pair in 10XG reads were trimmed to remove barcode sequences. For the skate, whole genome short read sequencing data was used instead for collecting *k-mers* as the k-mer histogram was abnormal in the linked reads. Using the k-mer histogram of the reads as a truth set for the genome, we compared *21-mers* collected from the primary and alternate assemblies.

Overlaying *k-mers* intersecting with a primary assembly in a read set is informative for inferring artificial duplications. All *k-mers* collected from single and two copies of the genome are expected to be found once in an assembly, assuming all two-copy *k-mers* are from the homozygous part of the genome with no haplotype (allele) specific duplication. A cutoff threshold was determined from the k-mer histogram of the reads as the maximum peak x 1.5 of the *k-mers* found in the assembly once. All distinct *k-mers* found more than once in the assembly within this cutoff of the k-mer counts found in reads are assembled more than expected. As an example, *k-mers* found in the assembly once peaked at 57x (**Supplementary Fig. 4**). The cutoff is therefore 86 (57 x 1.5). All *k-mers* found twice (blue), three (green), four (purple), or more (orange) times under 86x are considered and counted as falsely duplicated *k-mers*. Barcode trimming, *k-mer* counting, copy number spectrum, and false duplication counting was all performed with Merqury[16] `spectra-cn`.

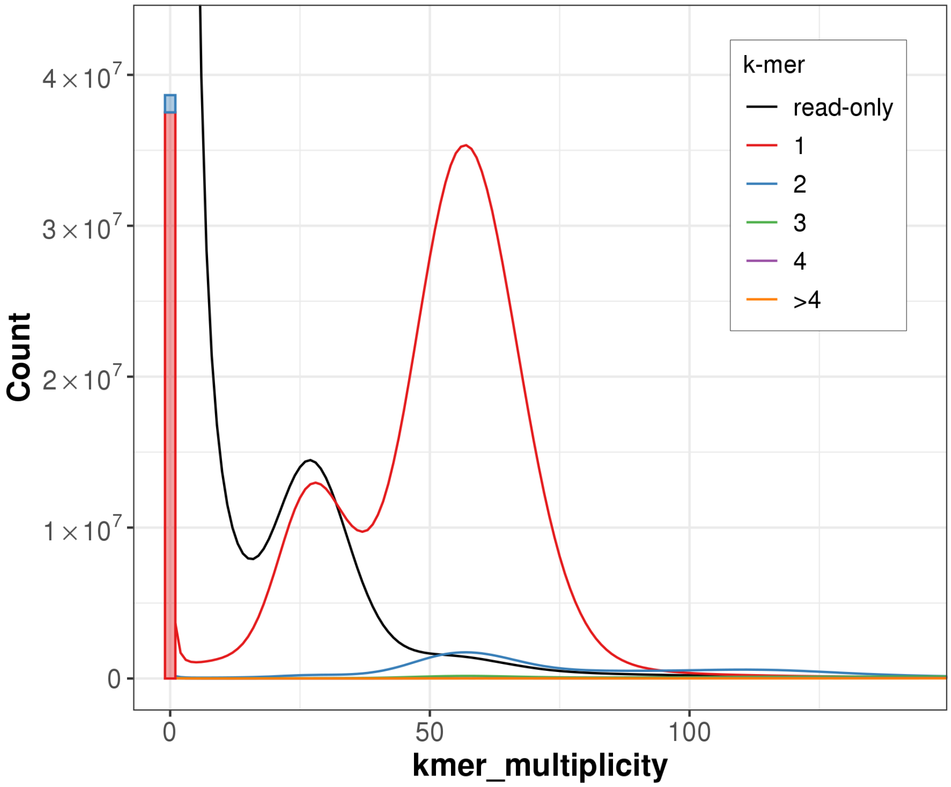


**Supplementary Fig. 4 |** Example histogram of the k-mer counts.

Once the *k-mers* were collected, the histogram (e.g. **Supplementary Fig. 4**) was prepared with spectra-cn of the Merqury code and shown in **Extended Data Fig. 3**:
https://github.com/marbl/merqury/blob/master/eval/spectra-cn.sh

Which generates histogram of the primary assembly in the following format:

```
Copies <tab> kmer_multiplicity <tab> Count
```

Where `Copies` are copies found in the assembly, `kmer_multiplicity` is the k-mer multiplicity found in reads, `Count` being the number of *k-mers* at each multiplicity.

The false duplication results were obtained using the following code: https://github.com/marbl/merqury/blob/master/eval/false_duplications.sh

```
cutoff=`cat $hist  | awk '$1==1 {print $2"\t"$3}' | awk -v max=0 'max<$2 {max=$2; mult=$1 } END {printf "%.0f\n", mult*(1.5)}'`
one_cp=`awk -v cutoff=$cutoff '$1==1 && $2<cutoff {sum+=$NF} END {print sum}' $hist`
two_cp=`awk -v cutoff=$cutoff '$1==2 && $2<cutoff {sum+=$NF} END {print sum}' $hist`
thr_cp=`awk -v cutoff=$cutoff '$1==3 && $2<cutoff {sum+=$NF} END {print sum}' $hist`
fou_cp=`awk -v cutoff=$cutoff '$1==4 && $2<cutoff {sum+=$NF} END {print sum}' $hist`
mor_cp=`awk -v cutoff=$cutoff '$1==">4" && $2<cutoff {sum+=$NF} END {print sum}' $hist`
DUPS_TOTAL=`echo "$one_cp $two_cp $thr_cp $fou_cp $mor_cp" | awk '{dup=$2+$3+$4+$5; all=dup+$1} END {print $1"\t"$2"\t"$3"\t"$4"\t"$5"\t"dup"\t"all"\t"(100*dup/all)}'`
echo -e "$hist\t$DUPS_TOTAL"
```

More details on the interpretation of the k-mer histogram can be found in the KAT[29] and Merqury[16] papers.

**Reliable blocks**

Regions with support from CLR subreads, linked reads, raw molecule and label information (bnx) of the optical maps, and Hi-C maps from the same individual were collected. Low or high coverage regions were excluded, which are indicators of mis-assemblies. To overcome mapping biases, we required at least two independent platforms to agree for being structurally ‘reliable’. For example, a repeat region longer than a CLR read may cause abnormal high coverage in CLR and linked reads from mapping biases, even if the region was well assembled locally. Longer range data such as optical maps and Hi-C interactions can complement this bias and indicate structural reliance.

Read mapping was performed individually for each platform, and coverage support information was collected on the primary assembly with Asset v1.0.2 (https://github.com/dfguan/asset). Here we include a brief description of each parameter for each platform as well as codes used to generate supporting regions. A manuscript for Asset will follow with more detailed information.

Regions with not enough support (hereby “low support”) were merged when less than 100 bp apart. Cutoff for defining low support differs per platform, as noted below per platforms. Reliable regions were calculated by excluding these low support regions from the assembly. Because sequencing coverage naturally drops at the end of scaffolds for optical maps and Hi-C, we included any low support region as “reliable” that overlaps 1 kbp of each ends in scaffolds (**Supplementary Table 4)**. All available optical maps were used, including those not used for hybrid scaffolding (**Supplementary Table 9**). Below are the command lines used to obtain these reliable blocks.

*Gaps in the reference*

Gaps and start, end of each scaffold information was obtained with `detgaps` for the primary assembly. Scaffold length was obtained with samtools. Length was then converted to region file, `asm.bed` and 1 kbp end coordinates was obtained as `asm.ends.bed` as following:

```
# Get gaps
$asset/bin/detgaps $name.fasta > gaps.bed

# Get scaffold length
samtools faidx $name.fasta

# Get scaffold region
awk '{print $1"\t0\t"$2}' $name.fasta.fai > asm.bed

# Get scaffold ends while ignore scaffolds <2kb
cat asm.bed | \
 awk '($3-$2) > 2000 {print $1"\t0\t1000\n"$1"\t"($3-1000)"\t"$3}' \
 > asm.ends.bed
```

*CLR coverage*

CLR reads were aligned to the primary assembly using `minimap2` with `-x map-pb`. Once all `.paf` files were collected, `ast_pb` was run with `-M $max`, which the maximum threshold was identified from `(sequencing mean coverage) x 2.5`. The mean coverage was inferred from the estimated haploid genome size / total bases. By default, `ast_pb` only includes read alignments with a minimum of 600 bases. Where `r` is a read, `s` the starting and `e` the ending coordinate of the alignment of `r`, any read alignment with `r(s+300, e-300)` is used to avoid errors at read ends. Regions with a minimum of 10 read alignments are excluded.

Brief help message is as follows:

```
Usage: aa_pb [options] <PAF_FILE> ...
Options:
  -m INT minimum coverage [10]
  -M INT maximum coverage [400]
  -l INT bases clipped at start and end coordinates of an alignment [300]
  -h  help
```

Command lines used are as follows:

```
# Align each qry subread .fasta file to the reference index
minimap2 -x map-pb -t $cpus $ref.idx $qry > $out.paf

# Accumulate coverage and exclude low and high coverage
pafs=`ls *.paf`
max=`echo $mean_cov | awk '{printf "%.0f\n", $1*2.5}'`
$asset/bin/ast_pb -M $max $pafs > pb_M.bed"
```

*Linked read coverage*

The `aligned.bam` file was re-used, which was generated to obtain mapping-based QV estimates using `longranger align`. To get the $max threshold, `ast_10x` was run in two rounds. The first round was run to get the average molecule coverage. The second round was run with `-C $max`, which is (average molecule coverage) x 3.5. Note this is set higher than what was used in CLR coverage as the linked reads were aligned to both haplotypes, thus here the average molecule coverage is closer to the haploid coverage.

By default, `ast_10x` requires regions to have at least 0.15 x (average molecule coverage) or 10 molecules, whichever is higher. Molecules are only considered when the average mapping quality of the reads in it is over 20, with the inferred molecule size being longer than 1 kbp. A molecule requires shared barcodes among at least 20 reads, where any two adjacent reads are less than 20 kbp apart. The maximum number of reads in a barcode is restricted to at most 1 million; however, based on the number of reads in barcodes, most barcodes meet this filtering criteria.

Brief help message is as follows:

```
Usage: aa_10x [options] <GAP_BED> <BAM_FILEs> ...
Options:
  -x BOOL use longranger bam [False]
```

```
  -b INT  minimum number of reads for each barcode [20]
  -B INT  maximum number of reads for each barcode [1M]
  -c INT minimum molecule coverage. This or -r will be used, whichever
is higher. [10]
  -r FLOAT minimum coverage ratio to the average coverage [.15]
  -C INT maximum coverage [inf]
  -q INT minimum average read mapping quality for each molecule [20]
  -l INT minimum length for a molecule [1000]
 -S INT maximum distance allowed between two adjacent reads with
identical barcode to be grouped as a molecule [20000]
  -a INT minimum number of barcodes for each molecule [5]
  -h     help
```

Command lines used are as follows:

```
# First round: accumulate molecule coverage to get the mean
$asset/bin/ast_10x -x gaps.bed aligned.bam > 10x.bed

# Avg. molecule coverage and max cutoff
mean_cov=`awk '{sum+=$1*$2; total+=$2} END {printf "%.0f\n", sum/total}'
TX.stat`
max=`echo $mean_cov | awk '{printf "%.0f\n", $1*3.5}'`

# Second round
$asset/bin/ast_10x -x -C $cutoff $gaps aligned.bam > 10x_C.bed
```

*Optical map raw molecule (bnx) coverage*

The curated primary assembly was first converted to in-silico reference cmaps for each label (Opt.1, 2, and 3) to align available bnx maps accordingly. The bnx were merged prior to alignment when multiple bnx were available from the same sequencing platform (Irys or Saphyr) and label. The bnx was aligned using `RefAligner` (Solve3.3_10252018) from Bionano Solve 3.3 using non-haplotype option of the sequencing platform (Irys or Saphyr), as we align bnx from both haplotypes to a pseudo-haplotype assembly. Molecule coverage was obtained with `ast_bion_bnx` using default options, which requires regions to have at least 10 molecule coverage or 0.5 x (average molecule coverage), whichever is higher. When multiple bnx files were used, all molecule coverage was gathered using the `union` function of Asset.

Brief help message is as follows:

```
Usage: ast_bion_bnx [options] <REF_CMAP> <QUERY_CMAP> <XMAP> <KEY_FN>
Options:
  -m INT   minimum molecule coverage [10]
  -M INT   maximum molecule coverage [inf]
  -r INT   minimum coverage ratio to mean coverage [.5]
  -s FLOAT minimum alignment confidence [0.0]
  -O STR   output directory [.]
```

```
  -h       help
```

Command lines used are as follows:

```
# Convert primary reference assembly fasta to cmap
perl  $solve_dir/HybridScaffold/10252018/scripts/fa2cmap_multi_color.pl
-e $enzyme 1 -i $ref -o $output_dir/fa2cmap

# Merge if multiple bnx are available from the same platform and label,
prefix is for example mLynCan4_Saphyr_BspQI
$tools/bionano/Solve3.3_10252018/RefAligner/7915.7989rel/RefAligner -if
$prefix.list -merge -o $prefix -bnx -stdout -stderr

# Align bnx to the reference cmap
python    $solve_dir/Pipeline/10252018/align_bnx_to_cmap.py    --prefix
$enzyme      --mol      $query_map      --ref      $ref_cmap       --ra
$solve_dir/RefAligner/7915.7989rel/    --nthreads    $cpus    --output
$output_dir/align                                            --optArgs
$solve_dir/RefAligner/7915.7989rel/optArguments_nonhaplotype_"$platfor
m".xml --pipeline $solve_dir/Pipeline/10252018/

# Convert to support regions of this .bnx.
# $rmap_fn, $qmap_fn, $xmap_fn, and $key_fn are the output files
# of the above fa2cmap
$asset/bin/ast_bion_bnx $rmap_fn $qmap_fn $xmap_fn $key_fn \
  > $output_dir/bionano_"$tech"_"$enzyme".bed \
  2>ast_bion_bnx_"$tech"_"$enzyme".log

# Merge support regions when multiple enzymes were used
$asset/bin/union bnx_*/bionano_*.bed > bn.bed
```

*Hi-C interaction coverage*

The `$genome.pri.bam` was re-used which was generated to plot the weighted length distribution of the Hi-C interactions. Coverage information was obtained using `ast_hic` with default options, which excludes regions with less than seven interactions. An interaction is inferred from the distance of a read pair, using the starting coordinates of each read while excluding N-base gaps. Only interactions less than 15 kbp were considered in coverage to avoid noisy long-range interactions for inferring structural reliability.

Brief help message is as follows:

```
Usage: aa_hic [options] <GAP_BED> <BAM_FILEs>
Options:
  -c INT minimum coverage [7]
  -C INT maximum coverage [inf]
  -q INT minimum alignment quality [0]
```

```
  -L INT maximum insertion length, gap excluded [15000]
  -h     help
```

Command lines used is as follows:

```
# Convert alignments to support information
$asset/bin/ast_hic gaps.bed *.bam > hic.bed
```

*Merging supportive regions*

Once all the support information for each platform is generated, low and high coverage regions are merged and good supporting regions of each platform are obtained using BEDTools[25] 2.92.2 with the following command lines:

```
# Get too-low and too-high coverage low support regions by merging blocks less than 100bp away
bedtools subtract -a asm.bed -b ${platform}.bed | bedtools merge -d 100 -i - > $platform.low_high.bed

# Trim off regions overlapping 1kb
bedtools subtract -a $platform.low_high.bed -b asm.ends.bed -A > $platform.low_high.trim1k.bed

# Get supporting region, using the low_high.bed to exclude <100bp blocks in between other blocks
bedtools subtract -a asm.bed -b $platform.low_high.bed > $platform.support.bed
```

In the last step, we accumulate the supporting evidence of all platforms and obtain supporting regions where >= 2 platforms agree using the following:

```
# Accumulate supports
$asset/bin/acc gaps.bed */*.support.bed > acc.bed 2> acc.log

# Merge to get reliable blocks
awk '$4>1' acc.bed | bedtools merge -i - > acc.gt2.mrg.bed

# Get low support regions by merging blocks <100bp apart
bedtools subtract -a asm.bed -b acc.gt2.mrg.bed | bedtools merge -d 100 -i - > low_support.bed

# Get the final support region as reliable blocks
bedtools subtract -a asm.bed -b low_support.bed > reliable.bed

# Exclude low supports in <1kb scaffold boundaries for excluding end-scaffold effects
```

```
bedtools subtract -A -a low_support.bed -b asm.ends.bed > low_support.trim1k.bed
```

The scripts used here are available on: https://github.com/VGP/vgp-assembly/tree/master/pipeline/asset.

**Telomere motifs**

Telomeric motif 6-mer AATCCC and its reverse complement TTAGGG were searched in the curated assemblies using a custom script. Once the motif sites were collected, regions of enriched telomeric motif signals in 1 kbp windows were collected over threshold *S*, corrected with the *k-mer* survival rate. The corrected threshold becomes *S* x *identity^k*, where *identity* is the approximate base accuracy, which we set as 99.9%. Because the spacing of enriched telomeric motifs varies across species and assembly quality, we collected windows using various thresholds from 10% to 25%. Windows were merged when closer than 100 bp in chromosome assigned scaffolds and reported in **Supplementary Table 5**. Number of windows within the beginning or ending scaffold coordinates were collected using bedtools[25]. Scaffold ends with windows found at a 15% threshold within 1 kbp end coordinates were reported in **Supplementary Table 4**. The following code was used to generate the data:

```
# Find telomere motifs, outputs pri.telomere
$VGP_PIPELINE/telomere/find_telomere.sh pri.fasta

# Find windows using variable $thresholds
java -cp $VGP_PIPELINE/telomere/telomere.jar FindTelomereWindows
pri.telomere 99.9 $threshold > pri.windows.$threshold

# Merge telomere windows when 100bp apart
cat pri.windows.$threshold | awk '{print $2"\t"$(NF-2)"\t"$(NF-1)}' |
sed 's/>//g' | bedtools merge -d 100  > pri.windows.$threshold.bed

# Get scaffold ends with variable $ends
cat pri.lens | awk -v ends=$ends '{if ($2>(ends*2)) {print
$1"\t0\t"ends"\n"$1"\t"($NF-ends)"\t"$NF} else {print $1"\t0\t"$NF}}' >
asm.ends.bed

# Get windows intersecting ends
bedtools intersect -wa -a pri.windows.$threshold.bed -b asm.ends.bed >
pri.windows.$threshold.$ends.ends.bed

# Get unique scaffold ends with a window
bedtools intersect -u -a asm.ends.bed -b pri.windows.$threshold.bed >>
pri.windows.$threshold.$ends.ends.u.bed
```

The full telomere motif finding script used is available on:
https://github.com/VGP/vgp-assembly/tree/master/pipeline/telomere/

**Base pair accuracy (QV) estimate**
We generated base level accuracy estimates (QVs) from the widely used mapping based approach as well as the newly developed k-mer based approach (**Fig. 3a** and **Supplementary Table 16**).

**Mapping based approach**
Longranger 2.2.2 was used for generating reference index and alignments. The reference index was generated on the combined primary and alternate assemblies with `longranger mkref`, and 10XG linked reads were aligned with `longranger align`.

```
longranger-2.2.2/longranger mkref $ref.fasta
longranger-2.2.2/longranger align \
--id=$genome \
--fastq=/data/rhiea/genome10k/$genome/genomic_data/10x/ \
--sample=$genome \
--reference=refdata-$ref \
--jobmode=slurm \
--maxjobs=500 \
--jobinterval=5000 \
--disable-ui \
--nopreflight
```

For reference assembly size larger than 4G, the following memory options were applied as Longranger was tuned for human genomes with `--override=$pipeline/longranger/override_4G.json` option.

The `override_4G.json` looks as following:
```
{
"ALIGNER_CS.ALIGNER._LINKED_READS_ALIGNER.BARCODE_AWARE_ALIGNER":
{ "chunk.mem_gb": 48 },
"ALIGNER_CS.ALIGNER._LINKED_READS_ALIGNER.MERGE_POS_BAM":
{ "join.mem_gb": 48 },
"ALIGNER_CS.ALIGNER._REPORTER.FILTER_BARCODES": { "join.mem_gb": 48 },
"ALIGNER_CS.ALIGNER._REPORTER.REPORT_LENGTH_MASS": { "chunk.mem_gb":
32 }
}
```

From the summary.csv that longranger align outputs, the mean coverage was obtained. Then, variants were called with freebayes 1.3.1 from the possorted_bam.bam file, which are indicative of base-pair errors as we are aligning reads from the same individual. This is the same step used for finding target bases to polish. We use --skip-coverage (mean_cov*12) to avoid variant calls in regions with excessive coverage depth, as the mapping results in this region is not reliable for variant calling. Basic filtering was applied on the called variants, (1) filter out low quality (>1) and (2) only select target sites called as homozygous-like variant calls (all reads support base change to one allele) and heterozygous-like variant calls when both suggestive alleles do not match the reference. In the latter case, the longest allele will be chosen. (https://github.com/VGP/vgp-assembly/tree/master/pipeline/freebayes-polish)

```
# Variant call
freebayes --bam $bam --skip-coverage $((mean_cov*12)) -f $fasta | bcftools view --no-version -Ou > bcf/$.bcf

# Filtering
bcftools view -i 'QUAL>1 && (GT="AA" || GT="Aa")' -Oz --threads=$threads $sample.bcf > $sample.changes.vcf.gz

# Collect number of bases affected (NUM_VAR)
bcftools view -H -Ov $genome.changes.vcf.gz | awk -F "\t" '{print $4"\t"$5}' | awk '{lenA=length($1); lenB=length($2); if (lenA < lenB ) {sum+=lenB-lenA} else if ( lenA > lenB ) { sum+=lenA-lenB } else {sum+=lenA}} END {print sum}' > $genome.numvar
NUM_VAR=`cat $genome.numvar`
echo "Total num. bases subject to change: $NUM_VAR"
```

Mappable region was obtained by excluding low (<3x) and high (mean_cov x 12) coverage. (https://github.com/VGP/vgp-assembly/tree/master/pipeline/qv)

```
# Num. of bases in mappable region (NUM_BP)
l_filter=3
h_filter=$((mean_cov*12))
samtools view -F 0x100 -u $bam | bedtools genomecov -ibam - -split > aligned.genomecov
awk -v l=$l_filter -v h=$h_filter '{if ($1=="genome" && $2>l && $2<h) {numbp += $3}} END {print numbp}' aligned.genomecov > $genome.numbp
NUM_BP=`cat $genome.numbp`

# QV calculation
QV=`echo "$NUM_VAR $NUM_BP" | awk '{print (-10*log($1/$2)/log(10))}'`
echo "QV of this genome $genome: $QV"
```

Number of bases affected (NUM_VAR) and bases in mappable regions (NUM_BP) are obtained per primary and alternate assemblies at the end, according to the reference sequence name.

**K-mer based approach**

Similarly to the mapping based approach, number of k-mers associated with base error (similar to NUM_VAR in mapping based approach) and total number of k-mers in the assembly (NUM_BP, respectively) are obtained and used for QV calculation. This assumes all k-mers occurring in the genome are observed in the short reads, and any k-mers not found in the short read but only in the assembly is considered to be originated from base error(s). More details regarding implementation is described in the Merqury paper[17]. This approach is independent from mapping, and is able to estimate QV across all assembled bases.

Once the k-mers were obtained as described in the **Quantifying false duplications with k-mers** section, Merqury `spectra-cn` was run using the following commands:

```
merqury.sh $read.meryl $pri.fasta $alt.fasta $out
```

Which generates spectra-asm and spectra-cn histograms shown in **Extended Data Fig. 3** as well as QV stats.

**RNA-seq and ATAC-seq Mappability**

RNAseq data from 44 zebra finch brain tissues (11 distinct regions, 4 adult male individuals) were trimmed for adaptors using fastq-mcf as part of the ea-utils package[30] 1.05 and mapped to the Sanger (TaeGut3.2.4) and VGP (bTaeGut1) genomic assemblies using STAR[31] 2.7.1a using default options. Reads were considered uniquely mapped if they matched only one location in the assembly, while multi mapped (<20) or mapped to too many (>20) were also counted. Total mapped was a summary of these categories. Mapping reports were summarized using MultiQC[32] 1.7 and compiled in R for significance testing. Following are the command lines used:

```
STAR
--runMode alignReads \
--runThreadN 8 \
--genomeDir ${REFERENCE} \
--readFilesIn                  ${TRIMDIR}/${SAMPLE}_R1_trimmed.fastq.gz
${TRIMDIR}/${SAMPLE}_R2_trimmed.fastq.gz \
--readFilesCommand zcat \
--outFileNamePrefix ${OUTPUT}/vgp_${SAMPLE} \
--outSAMtype BAM SortedByCoordinate
```

ATAC-seq libraries from 12 zebra finch brain tissues (4 distinct regions, 3 male birds per region) were prepared using the Omni-ATAC-seq method (https://protocolexchange.researchsquare.com/article/nprot-6107/v1) and were sequenced on the Illumina NextSeq 500 with 75 bp paired end reads. The reads were then trimmed with Trim Galore[33] 0.6.5. Next, the reads were aligned to each assembly using Bowtie2[34] 2.4.1. The average of each mapping statistic summary log (mapped zero times, mapped exactly 1 time and mapped multiple times) were calculated for each assembly. Following are the command lines used:

```
# Trimming adapters
trim_galore --paired --nextera myRead_R1.fastq myRead_R2.fastq

# Assembly alignment
bowtie2 -x genome_assembly.fa --sensitive -1 my_reads_trimmed_1.fq.gz -
2     my_reads_trimmed_2.fq.gz     -S     my_reads_trimmed.sam     2>
my_reads_trimmed.log
```

Mapping statistics for both RNA-seq and ATAC-seq on each assembly were tested for significant difference in means using a paired two sample t-test (alpha=0.05).

**False gene annotation in previous assemblies**

We detected evidence of erroneous coding sequences in previous assemblies of the zebra finch, platypus, and climbing perch for the genes which are related to the important trait of the animal[35,36] or, included in the BUSCO gene set[26,27]. To identify the erroneous annotation such as false duplication or truncated sequences which would affect the coding sequences, we collected exon sequences from VGP annotation of the genes and performed blastn v2.6.0+ search[37] against both previous and VGP assembly with those options -task blastn, -perc_identity 90, and -evalue 0.00001 (**Supplementary table 21**). Among the hits found from blast search, we defined false duplication of an exon as a case when duplicated hits within the same scaffold were found on the previous assembly only. Also, we detected the truncated exons where the length of blast hit was shorter than the length of query exon. For visualization, we used Gene Structure Display Server 2.0+[38] and manually modified the display in order to handle small discrepancies between elements. For the intuitive visualization of platypus' vitellogenin-2 gene, we visualized only the scaffolds with more than three blast hits of the previous assembly.

### Summplementary References


1. Chin, C.-S. *et al.* Phased diploid genome assembly with single-molecule real-time sequencing. *Nat. Methods* **13**, 1050–1054 (2016).
2. Kronenberg, Z. N. *et al.* Extended haplotype phasing of de novo genome assemblies with FALCON-Phase. *bioRxiv* 327064 (2019) doi:10.1101/327064.
3. Koren, S. *et al.* De novo assembly of haplotype-resolved genomes with trio binning. *Nat. Biotechnol.* (2018) doi:10.1038/nbt.4277.
4. Durand, N. C. *et al.* Juicer Provides a One-Click System for Analyzing Loop-Resolution Hi-C Experiments. *Cell Syst.* **3**, 95–98 (2016).
5. Zhang, G. *et al.* Comparative genomics reveals insights into avian genome evolution and adaptation. *Science* **346**, 1311–1320 (2014).
6. Dudchenko, O. *et al.* De novo assembly of the Aedes aegypti genome using Hi-C yields chromosome-length scaffolds. *Science* **356**, 92–95 (2017).
7. Ghurye, J. *et al.* Integrating Hi-C links with assembly graphs for chromosome-scale assembly. *PLOS Comput. Biol.* **15**, e1007273 (2019).
8. Bradnam, K. R. *et al.* Assemblathon 2: evaluating de novo methods of genome assembly in three vertebrate species. *GigaScience* **2**, (2013).
9. Korlach, J. *et al.* De novo PacBio long-read and phased avian genome assemblies correct and add to reference genes generated with intermediate and short reads. *GigaScience* **6**, 1–16 (2017).
10. Walker, B. J. *et al.* Pilon: An Integrated Tool for Comprehensive Microbial Variant Detection and Genome Assembly Improvement. *PLOS ONE* **9**, e112963 (2014).
11. Garrison, E. & Marth, G. Haplotype-based variant detection from short-read sequencing. *ArXiv12073907 Q-Bio* (2012).
12. Roach, M. J., Schmidt, S. A. & Borneman, A. R. Purge Haplotigs: allelic contig reassignment for third-gen diploid genome assemblies. *BMC Bioinformatics* **19**, 460 (2018).
13. O'Quin, C. T., Drilea, A. C., Conte, M. A. & Kocher, T. D. Mapping of pigmentation QTL on an anchored genome assembly of the cichlid fish, Metriaclima zebra. *BMC Genomics* **14**, 287 (2013).
14. Albertson, R. C. *et al.* Genetic basis of continuous variation in the levels and modular inheritance of pigmentation in cichlid fishes. *Mol. Ecol.* **23**, 5135–5150 (2014).
15. Yunis, J. J. & Hoffman, W. R. Nuclear Enzymes, Fragile Sites, and Cancer. *J. Gerontol.* **44**, 37–44 (1989).
16. Rhie, A., Walenz, B. P., Koren, S. & Phillippy, A. M. Merqury: reference-free quality, completeness, and phasing assessment for genome assemblies. *bioRxiv* 2020.03.15.992941 (2020) doi:10.1101/2020.03.15.992941.
17. Ondov, B. D. *et al.* Mash: fast genome and metagenome distance estimation using MinHash. *Genome Biol.* **17**, 132 (2016).
18. Morgulis, A., Gertz, E. M., Schäffer, A. A. & Agarwala, R. WindowMasker: window-based masker for sequenced genomes. *Bioinforma. Oxf. Engl.* **22**, 134–141 (2006).
19. Altschul, S. F., Gish, W., Miller, W., Myers, E. W. & Lipman, D. J. Basic local alignment search tool. *J. Mol. Biol.* **215**, 403–410 (1990).
20. Kapustin, Y., Souvorov, A., Tatusova, T. & Lipman, D. Splign: algorithms for computing spliced alignments with identification of paralogs. *Biol. Direct* **3**, 20 (2008).
21. Li, H. Minimap2: pairwise alignment for nucleotide sequences. *Bioinforma. Oxf. Engl.* **34**, 3094–3100 (2018).
22. Lowe, T. M. & Eddy, S. R. tRNAscan-SE: a program for improved detection of transfer RNA genes in genomic sequence. *Nucleic Acids Res.* **25**, 955–964 (1997).
23. Nawrocki, E. P. *et al.* Rfam 12.0: updates to the RNA families database. *Nucleic Acids Res.* **43**, D130-137 (2015).

24. Li, H. *et al.* The Sequence Alignment/Map format and SAMtools. *Bioinformatics* **25**, 2078–2079 (2009).
25. Quinlan, A. R. & Hall, I. M. BEDTools: a flexible suite of utilities for comparing genomic features. *Bioinformatics* **26**, 841–842 (2010).
26. Simão, F. A., Waterhouse, R. M., Ioannidis, P., Kriventseva, E. V. & Zdobnov, E. M. BUSCO: assessing genome assembly and annotation completeness with single-copy orthologs. *Bioinforma. Oxf. Engl.* **31**, 3210–3212 (2015).
27. Waterhouse, R. M. *et al.* BUSCO Applications from Quality Assessments to Gene Prediction and Phylogenomics. *Mol. Biol. Evol.* **35**, 543–548 (2018).
28. Jain, C., Koren, S., Dilthey, A., Phillippy, A. M. & Aluru, S. A fast adaptive algorithm for computing whole-genome homology maps. *Bioinformatics* **34**, i748–i756 (2018).
29. Mapleson, D., Garcia Accinelli, G., Kettleborough, G., Wright, J. & Clavijo, B. J. KAT: a K-mer analysis toolkit to quality control NGS datasets and genome assemblies. *Bioinformatics* **33**, 574–576 (2017).
30. Aronesty, E. Comparison of Sequencing Utility Programs. *Open Bioinforma. J.* **7**, (2013).
31. Dobin, A. *et al.* STAR: ultrafast universal RNA-seq aligner. *Bioinformatics* **29**, 15–21 (2013).
32. Ewels, P., Magnusson, M., Lundin, S. & Käller, M. MultiQC: summarize analysis results for multiple tools and samples in a single report. *Bioinformatics* **32**, 3047–3048 (2016).
33. Krueger, F. Trim galore. *Wrapper Tool Cutadapt FastQC Consistently Apply Qual. Adapt. Trimming FastQ Files* **516**, 517 (2015).
34. Langmead, B. & Salzberg, S. L. Fast gapped-read alignment with Bowtie 2. *Nat. Methods* **9**, 357–359 (2012).
35. Zebra Finch Expression Brain Atlas. http://www.zebrafinchatlas.org/.
36. Robinson, R. For mammals, loss of yolk and gain of milk went hand in hand. *PLoS Biol.* **6**, e77 (2008).
37. Camacho, C. *et al.* BLAST+: architecture and applications. *BMC Bioinformatics* **10**, 421 (2009).
38. Hu, B. *et al.* GSDS 2.0: an upgraded gene feature visualization server. *Bioinforma. Oxf. Engl.* **31**, 1296–1297 (2015).
